# In-solution buffer-free digestion for the analysis of SARS-CoV-2 RBD proteins allows a full sequence coverage and detection of post-translational modifications in a single ESI-MS spectrum

**DOI:** 10.1101/2021.05.10.443404

**Authors:** Luis Ariel Espinosa, Yassel Ramos, Ivan Andújar, Enso Onill Torres, Gleysin Cabrera, Alejandro Martín, Diamilé González, Glay Chinea, Mónica Becquet, Isabel González, Camila Canaán-Haden, Elías Nelson, Gertrudis Rojas, Beatriz Pérez-Massón, Dayana Pérez-Martínez, Tamy Boggiano, Julio Palacio, Sum Lai Lozada-Chang, Lourdes Hernández, Kathya Rashida de la Luz Hernández, Saloheimo Markku, Vitikainen Marika, Yury Valdés-Balbín, Darielys Santana-Medero, Daniel G. Rivera, Vicente Vérez-Bencomo, Mark Emalfarb, Ronen Tchelet, Gerardo Guillén, Miladys Limonta, Eulogio Pimentel, Marta Ayala, Vladimir Besada, Luis Javier González

## Abstract

Subunit vaccines based on the receptor-binding domain (RBD) of the spike protein of SARS-CoV-2, are among the most promising strategies to fight the COVID-19 pandemic. The detailed characterization of the protein primary structure by mass spectrometry (MS) is mandatory, as described in ICHQ6B guidelines. In this work, several recombinant RBD proteins produced in five expression systems were characterized using a non-conventional protocol known as in-solution buffer-free digestion (BFD). In a single ESI-MS spectrum, BFD allowed very high sequence coverage (≥ 99 %) and the detection of highly hydrophilic regions, including very short and hydrophilic peptides (2-8 amino acids), the His_6_-tagged C-terminal peptide carrying several post-translational modifications at Cys^538^ such as cysteinylation, glutathionylation, cyanilation, among others. The analysis using the conventional digestion protocol allowed lower sequence coverage (80-90 %) and did not detect peptides carrying some of the above-mentioned post-translational modifications. The two C-terminal peptides of a dimer [RBD(_319-541_)-(His)_6_]_2_ linked by an intermolecular disulfide bond (Cys_538_-Cys_538_) with twelve histidine residues were only detected by BFD. This protocol allows the detection of the four disulfide bonds present in the native RBD and the low-abundance scrambling variants, free cysteine residues, O-glycoforms and incomplete processing of the N-terminal end, if present. Artifacts that might be generated by the in-solution BFD protocol were also characterized. BFD can be easily implemented and we foresee that it can be also helpful to the characterization of mutated RBD.

## Introduction

The new coronavirus SARS-CoV-2 reported for the first time in Wuhan, China (1) has rapidly spread all over the world, causing more than 3.3 million deaths and 159 million of infected people until May 2021 (2). Currently, this pandemic represents a major threat for the health, the economy and the whole society.

The development of effective vaccines as well as the universal access for their massive introduction are urgently needed to control the COVID-19 pandemic worldwide (3). Nowadays, there are several vaccine platforms being evaluated according to the draft landscape published by the World Health Organization (4), including inactivated and live attenuated virus, non-replicating and replicating viral vectors, DNA-, mRNA-, virus-like particles, and protein subunit vaccines. Some of them have already been approved by the WHO and regulatory authorities and introduced with favorable results in the clinic (5).

SARS-CoV-2 uses the receptor-binding domain (RBD) of the spike (S) protein for entry into the host cells through a high affinity interaction with its cell-surface expressed receptor, the angiotensin-converting enzyme 2 (ACE2) (6, 7). The RBD has been proposed for the rational development of protective vaccines against SARS-CoV-2 (8, 9) and nowadays subunit vaccines are well-represented among the candidates investigated in preclinical studies and clinical trials, according a recent WHO report (4). For a successful introduction of vaccines into the clinic, the immunogens need to be produced at scale and prices affordable for all, including middle- and low-income countries (3).

Probably this is one of the reasons why RBD of SARS-CoV-2, besides its production in mammalian cells (10), has also been produced in several systems such as insect cells (11), yeast (12, 13), C1 *Thermothelomyces heterothallica* (formerly *Myceliophthora thermophila*) (14, 15), baculovirus-silkworm system (16) and bacteria (17) despite of the challenge that represents the expression of a non-globular protein with four disulfide bonds and the requirement of the N-glycosylation for its proper expression and folding (12).

The ICHQ6B guidelines (18), harmonized among most important regulatory agencies worldwide, state the key role of mass spectrometry (MS) to develop well-characterized products in the biotechnology industry. MS is the analytical tool of choice for the verification of the amino acid sequence, to demonstrate the integrity of the N- and C-terminal ends, and to detect post-translational modifications (PTMs) in natural and recombinant proteins. The presence of PTMs may modify the physico-chemical, and immunological properties of the proteins. In particular, a disulfide bonds arrangement identical to the present in the native protein, is mandatory for biotherapeutics as well as for vaccine development in cases where the antigen should be well folded to raise conformational and topological neutralizing antibodies (19, 20).

Sample processing prior to MS analysis also plays a determinant role in the quality of the results. In the characterization of recombinant proteins, an efficient proteolytic digestion and the recovery of the proteolytic peptides is mandatory to obtain the highest sequence coverage, including the identification of PTMs. In particular, if electrospray ionization mass spectrometry (ESI-MS) is used, a desalting step is needed to ionize properly the proteolytic peptides. This step, although necessary, often comprises the recovery of highly hydrophilic and hydrophobic peptides when, either commercial or in-house prepared micro-columns, based on reverse phase chromatography are used. Arbeitman et al (12) analyzed by MALDI-MS the in-solution tryptic digests of two reduced and S-alkylated recombinant RBD of SARS-CoV-2 expressed in *P. pastoris* and in a mammalian cell line (HEK-293T). The tryptic peptides were desalted by C18-ZipTips prior to mass spectrometric analysis and although sequence identity was unequivocally demonstrated in both cases, the sequence coverage was only 40 and 60% for the proteins expressed in *P. pastoris* and HEK-293T cells, respectively. In that experiment, some of the internal short and/or hydrophilic peptides (^318^SR^319^, ^379^CYGVSPTK^386^, ^418^IDIAYNYK^424^, ^455^LFR^457^, and ^530^STNLVK^535^) were not verified for any proteins. The C-terminal peptide containing the tandem repeat of six histidine residues (LPETGHHHHHH), was only detected for the recombinant protein expressed in *P. pastoris,* suggesting variable results in the desalting step. The unmodified peptide located at the N-terminal region (^320^VQP***<u>T</u>***E***<u>S</u>***IVR^328^) was only assigned for the case of RBD expressed in *P. pastoris*. Probably the presence of O-glycosylation site(s) at Thr^323^/Ser^325^ (21) makes this peptide more hydrophylic when RBD was expressed in HEK-293T cell line and in consequence the peptide was not retained in the desalting step and not detected in MALDI-MS analysis (12). Also two internal peptides (F^329^-R^346^ and D^467^-R^509^) containing three (Cys^336^, Cys^480^ and Cys^488^) out of the eight cysteine residues were not detected in any protein. The peptide ^329^FP**N**ITNL*<u>C</u>*PFGEVF**N**ATR^346^ was not detected propably because it contains two potential N-glycosylation sites at Asn^331^ and Asn^343^ (21) fully occupied by N-glycans and a deglycosylation step with PNGase F was not conceived in the sample processing to facilitate its detection. On the other hand, the long and hydrophobic peptide D^467^-R^509^ containing two cysteine residues (Cys^480^-Cys^488^) probably was not efficiently eluted from the C18-Ziptips (12). In the same manuscript, the arrangement of disulfide bonds and the presence of free cysteine residues were not verified by mass spectrometry because of the reduction and S-alkylation steps included in the sample processing, probably introduced to make more efficient the tryptic digestion, impaired their detection (12). The detection of non-desired free cysteine residues in the molecule, even present as low-abundance species, is also a matter of great importance because they may promote disulfide exchange and generate scrambling variants (22).

In our laboratory, we initially demonstrated that proteins separated by SDS-PAGE can be efficiently in-gel desalted and digested in water with trypsin in absence of traditional saline buffers (23). This procedure avoids a desalting step of the proteolytic peptides and allows their direct analysis by ESI-MS (23). This non-conventional procedure moves away from the conventional canons of biochemistry, but it avoids the loss of hydrophobic tryptic peptides and guaranteed sequence coverage higher than the achieved by the traditional in-gel digestion protocol. Our procedure has allowed full-sequence coverage of ATPase subunit 9, the most hydrophobic protein of *S. cerevisiae* proteome by an in-gel digestion procedure (23).

In the biotechnology industry, for the quality control of recombinant proteins, in-solution protein digestion is more frequently used than in-gel digestion procedures due to its greater efficiency and simplicity and also because sample amount is generally not a concern.

Recently, the principles of the in-gel buffer-free digestion protocol (23) were extended to in-solution buffer-free digestion (BFD) of other proteins (24). In-solution BFD protocol improved the sequence coverage of certain regions of proteins represented by short and hydrophilic peptides including some N-glycopeptides, short peptides linked by disulfide bonds as well as hydrophilic C-terminal peptides of proteins that contain a tandem repeat of six histidine residues (24).

In particular, the introduction of a tandem repeat of six histidine residues at the N- or at the C-terminal end of the recombinant proteins is frequently used to facilitate the purification by immobilized metal affinity chromatography. This procedure although efficient, can turn the N- and C-terminal peptides, into more hydrophilic species and it makes more difficult their recovery from the desalting step prior ESI-MS analysis. The integrity of the N-and C-terminal ends is an aspect requested by the ICHQ6B due to their propensity to be proteolytic processed by the host cell proteases and at the same time they carry PTMs (18).

Considering that subunit vaccines based on the recombinant RBD are very well-represented among the different strategies for the development of vaccines against COVID-19, in this work, we adapted the in-solution BFD protocol (24) to the analysis of the products of six gene constructs containing the RBD sequence from SARS-CoV-2 obtained in five different expression systems, such as mammalian cells (CHO and HEK-293T), yeast (*P. pastoris*), bacteria (*E. coli*) and fungus (*Thermothelomyces heterothallica*). Unlike the standard protocol that uses salt buffers and desalting through reverse phase microcolumns, the implemented BFD method avoids buffers and desalting is carried out by precipitation, allowing very high sequence-coverage (≥ 99 %) of these RBDs, and the detection of PTMs including those located at the N- and the C-terminal end. The BFD protocol allowed the identification, in a single mass spectrum, of the four native disulfide bonds as well as scrambled disulfide bonds, the presence of free cysteine residues, N- and O-glycosylation, and other PTMs of known and unknown nature linked to an unpaired cysteine residue present in some of the analyzed RBD molecules.

## Materials and methods

### Cloning expression and purification of RBD variants

Seven RBD recombinant proteins, produced at laboratory scale in a wide range of host cells, were used as model antigens to develop and refine suitable analytical methods for RBD characterization. **Table 1** summarizes their sequences. The procedures for cloning, expression and purification are further described below.

**Table 1.**
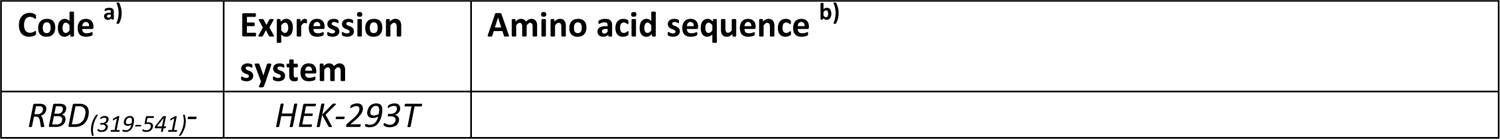
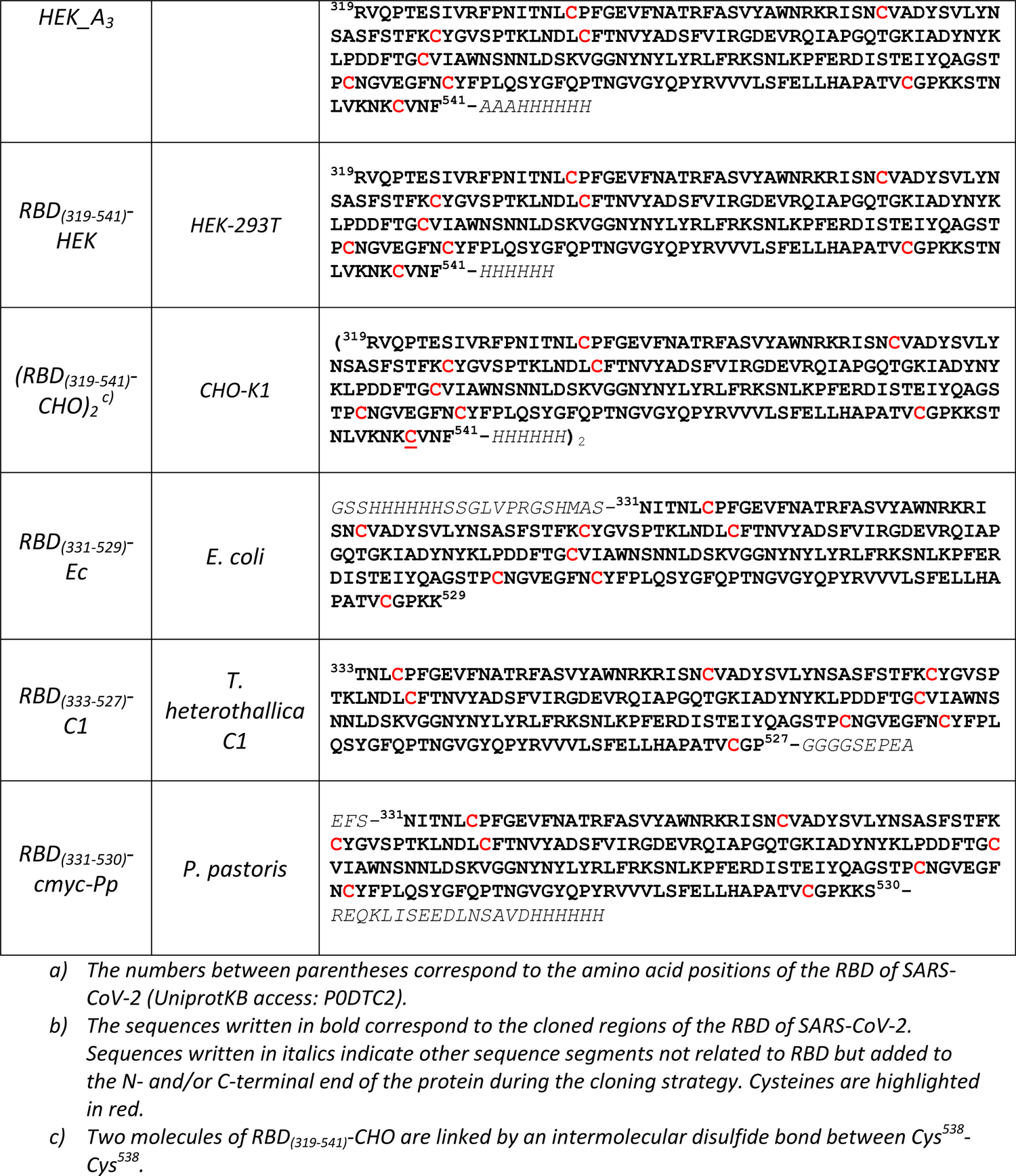
Sequences of the recombinant receptor binding domain of SARS-CoV-2 characterized in this work.

### *RBD*(*_319-541_*)*-HEK_A_3_* and *RBD_(319-541)_-HEK* from transient transfection of HEK-293T cells

The coding sequence for RBD_319-541_ was optimized for mammalian cell expression (hamster, *Cricetulus griseus*), using the online gene optimization tools provided by Eurofins (Germany). The optimized nucleotide sequence was assembled and amplified by PCR using gene fragments synthesized by Eurofins and oligonucleotides synthesized at CIGB (Cuba), and cloned into pCMX/His vector through BssHII and NotI restriction sites. The use of NotI site resulted in the introduction of three alanine residues between RBD and His_6_ tag in the coded protein. A second version of the genetic construct (coding for a recombinant protein without additional Ala residues) was obtained by assembling RBD_319-541_ sequence directly followed by the gene ofHis_6_ tag and a stop codon, and cloning that insert in the same expression vector. HEK-293T cells adapted to grow in suspension in Freestyle F-17 medium were transfected with the resulting genetic constructs, using linear polyethyleneimine as the transfection agent. Transfected cells were fed with fresh media plus tryptone (0.5% final concentration) 48 h post-transfection. Supernatant was harvested five days after feeding, and the recombinant protein was purified from cell culture supernatant by IMAC with Ni-NTA Sepharose. Proteins were buffer-exchanged with PBS pH 7.4 using PD-10 columns (GE Healthcare).

### (RBD_(319-541)_-CHO)_2_ from transduced CHO-K1 cells

The whole expression cassette including the gene coding for RBD_319-541_ (from CMV promoter to His_6_ tag and stop codon) was amplified by PCR from pCMX/His (see the previous section) and re-cloned into the lentiviral vector pL6WBlast (CIGB, Cuba) with the restriction enzymes XhoI and EcoRV. Lentiviral particles containing the gene of interest were produced and used for stable transduction of CHO-K1 cells. Cells producing the recombinant protein were grown in protein-free medium (a mixture of a proprietary Center for Molecular Immunology medium with PFHMII) with shaking. Recombinant RBD_319-541_ was purified from cell culture supernatant by IMAC with Ni-IDA Sepharose, followed by SP cation exchange chromatography. Dimeric and monomeric fractions were subsequently isolated by size exclusion chromatography with Superdex 200 in PBS pH 7.4.

### RBD_(331-529)_-Ec from Escherichia coli

A sequence coding for residues 331-530 of the Spike protein from SARS-CoV-2 was obtained by reverse transcription from nasopharyngeal swabs of COVID-19 patients and amplification by PCR, adding a *Nhe I* site at the 5’ end and a stop codon and a *Sal I* site at the 3’ end. The resulting amplicon, after digestion with these enzymes, was cloned in-frame into *Nhe I/Sal I*-digested pET28a+ (Novagen, USA). Transformation of the resulting construct into NiCo21(DE3) cells (New England Biolabs, USA) and culture in a ZY5052 medium (25) resulted in the expression of an N-terminally His-tagged RBD, which was purified from inclusion bodies using immobilized metal affinity chromatography (IMAC) under denaturing conditions. Protein was eluted using 16 mM phosphate-buffered, 300 mM NaCl, 150 mM imidazole, pH 6.8. Fractions were pooled and buffer-exchanged to sodium carbonate/hydrogen carbonate buffer pH 10 using a Sephadex G-25 column (GE Healthcare, UK). Recombinant protein was stored at −20 °C until required. An aliquot of 200 μg was buffer-exchanged to PBS (12 mM Na_2_HPO_4_.2H_2_O, 3 mM NaH_2_PO_4_.2H_2_O, 150 mM NaCl pH 7.4) using Sephadex PD-10 columns (Pharmacia).

### RBD_(333-527)_-C1 expressed in thermophile C1, Thermothelomyces heterothallica

A sequence coding for a C1 endogenous signal sequence, the residues 333-527 of the Spike protein from SARS-CoV-2 (RBD), a GlySer-linker and the C-tag flanked by PmeI sites was amplified by PCR from a synthetic fragment provided by GenScript (USA), codon-optimized for expression in a low protease background C1, *Thermothelomyces heterothallica* (formerly *M. thermophila*) strain. The fragment was cloned into a C1 expression vector under a C1 endogenous promoter. The ready expression vector and a mock vector partner needed for a complete marker gene were digested with PmeI and co-transformed to a low protease background C1 strain. The transformation was done with protoplast/PEG method (14) and transformants were selected for nia1+ phenotype and hygromycin resistance. Transformants were streaked once to selective medium and screened by Western blotting of culture supernatant samples from 24-well plate cultures.

C1 transformants producing RBD-C-tag were purified by single colony plating and analyzed by PCR for correct integration of the expression cassette and clone purity. A purified clone was then selected for cultivation in a 1 L bioreactor in a fed-batch process with a medium containing glucose as the carbon source and (NH_4_)_2_SO_4_ and yeast extract as the nitrogen sources. The fermentation was carried out for 5 days at pH 6.8 at 38 °C. RBD-C-tag was purified from 116 h time point of the fermentation by C-tag affinity chromatography with the CaptureSelect C-tagXL resin (Thermo Fisher Scientific) according to the manufacturer’s protocols. The eluted product was dialyzed against PBS.

### RBD_(331-530)_-cmyc from Pichia pastoris

A sequence coding for residues 331-530 of the Spike protein from SARS-CoV-2 flanked by *EcoR I* and *Xba I* sites was amplified by PCR from a synthetic fragment provided by Eurofins (Germany), codon-optimized for expression into *Saccharomyces cerevisiae* (*S. cerevisiae*). After digestion with these enzymes, the fragment was cloned in-frame with the *S. cerevisiae* alpha factor pre-pro peptide into *EcoR I/Xba I*-digested pPICK-αA (a pPICZ-αA derivative bearing a G418 resistant selection marker). Transformation of the resulting construct into *P. pastoris* GS115, selection of clones resistant to high G418 concentrations, and culture of one representative clone in BMGY/BMMY (Invitrogen, USA) resulted in the secretion into the supernatant of an RBD variant tagged C-terminally with *C-myc* and His_6_ sequences, which was further purified by IMAC on Ni-NTA agarose and gel filtration.

### In-solution buffer-free digestion (BFD) protocol

Fifty micrograms of the protein dissolved in PBS (pH 7.4) containing 0.5 M guanidine hydrochloride reacted with 5 mM N-ethylmaleimide (NEM) during 30 min at room temperature (22 °C). One microliter of PNGase F (New England Biolabs) was added and the deglycosylation reaction proceeded during two hours at 37 °C. Also, N-glycosylated *RBD_(333-527)_-C1* was reduced with 10 mM dithiotreitol and 0.2 M Tris-HCl buffer pH 8.0 for 1 h at 37 °C, and then S-alkylated with 25 mM iodoacetamide in the dark for 20 min at 22 °C. All samples were cooled at room temperature and proteins were precipitated with ten volumes of cold acetone (−20 °C) or 80% ethanol (v/v) and the solution was kept at −80±5 °C during 1 h. The sample was centrifuged at 10000 rpm during 5 minutes and the supernatant was discarded. The precipitate was washed by vortexing with 75% cold acetone or ethanol (−20 °C), centrifuged at 10000 rpm during 5 min and the supernatant was discarded. This procedure was repeated twice and the final precipitate was dried up in a vacuum centrifuge during 15 min. The precipitate was dissolved in 50 μL of 20% (v/v) acetonitrile in water solution with 1 min vortex and 10 min of sonication in a water bath. One microgram of sequencing grade trypsin (Promega) dissolved in water was added to the protein solution and the specific proteolytic digestion proceeded for 16 h at 37 °C in a thermomixer (Thermo Fisher Scientific). Digestion was centrifuged at 10000 rpm during 1 min and 4 μL of the resultant mixture of tryptic peptides was mixed with 0.3 μL of 90% formic acid and it was loaded into a metal coated borosilicate nanocapilar for mass spectrometric analysis.

### Standard digestion (SD) protocol

Fifty micrograms of the protein dissolved in PBS (pH 7.2) containing 0.5 M guanidine hydrochloride reacted with 5 mM NEM during 30 min at room temperature (22 °C). One microliter of PNGase F (New England Biolabs) was added and the deglycosylation reaction proceeded during two hours at 37 °C. The sample was four-fold diluted and the protein digested in presence of 0.2 M Tris-HCl buffer pH 8.0 and 1 μg of sequencing grade trypsin (Promega) previously dissolved in 20 mM acetic acid. Tryptic digestion proceeded for 16 h at 37 °C and digestion was stopped with a final 5% formic acid (v/v). The resulting peptides were desalted with ZipTip C18 (Millipore, USA), washed with 0.2% (v/v) formic acid solution, and eluted in 4 μL of 60% acetonitrile in water containing 0.2% formic acid (v/v).

### Electrospray ionization mass spectrometry analysis

Seven micrograms of the proteins deglycosylated with PNGase F as described above and the N-glycosylated *RBD_(333-527)_-C1,* were mixed with equal volume of 6 M guanidine hydrochloride solution and desalted by using ZipTip C18 (Millipore, USA). The protein was extensively washed with 0.2% (v/v) formic acid solution and finally eluted in 3 μL of 60% acetonitrile in water containing 0.2% formic acid (v/v). The elution was loaded into the metal coated nanocapillary for ESI-MS analysis.

The mixture of tryptic peptides was analyzed in a hybrid orthogonal QTof-2 tandem mass spectrometer (Micromass, Manchester, UK) by spraying the sample into the ion source using 1200 and 35 volts for the capillary and the entrance cone, respectively. The ESI-MS were acquired from *m/z* 200-2000 and the multiply-charged ions were manually fragmented by collision induced dissociation using appropriated collision energies (20-50 eV) to obtain structural information in the MS/MS spectra. Argon was used as a collision gas and the mass spectra were processed by using MassLynx v4.1 (Micromass, UK). The ESI-MS/MS of tryptic peptides with z≥3+ were deconvoluted by MaxEnt 3.0. The multiply-charged ESI-MS spectrum (*m/z* 400-3000) of the protein deglycosylated with PNGase F was deconvoluted (mass 5000-70000) by using the MaxtEnt1.0 tool. The theoretical *m/z* for tryptic peptides as well as for the intact protein was calculated by using the peptide and protein editor available in the MassLynx v4.1 software (Micromass, UK).

### SDS-PAGE analysis

*RBD_(319-541)_-HEK_A_3_*, *(RBD_(319-541)_-CHO)_2_* and *RBD_(331-529)_-Ec* proteins were separated by SDS-PAGE as described by Laemmli (26), under reducing and non-reducing conditions. Two micrograms of N-glycosylated and deglycosylated proteins were applied in a 12.5%T, 3%C acrylamide-bisacrylamide separating gel at 30 mA/gel until the tracking dye left the gel. Proteins were detected by silver staining (27) or Coomassie Brilliant Blue G-250, gel images were analyzed with a GS-900 calibrated imaging densitometer (Bio-Rad) and processed with Image Lab v6.0 software (Bio-Rad).

### NP-HPLC analysis

N-glycosylation profile was determined by using the procedure described by Guile et al. (28). Briefly, the N-glycans released by PNGase F treatment were derivatized with 2-amino benzamide (2AB) by reductive amination. The chromatographic separation was carried out in an HPLC Prominence-Shimadzu (Japan) using a linear gradient from 20% to 53% of 50 mM, pH 4.4 ammonium formate (solution A) and pure acetronitrile (solution B). 2AB N-glycans separation was performed on an Amide-80 column (TSKgel 250×46 mm, 5 µm, Tosohaas, Japan) and the derivatized oligosaccharides were detected on-line by fluorescence using an excitation and detection wavelengths of 330 nm and 420 nm, respectively. The structural assignment was performed by comparing the experimental GU values with the GlycoStore database (https://glycostore.org/). GU values were calculated from the retention time of each peak using as reference an HPLC separation ran under similar conditions for the 2AB derivatives of a dextran ladder generated by acid partial hydrolysis. Glycans structures were represented according to GlycoStore nomenclature.

## Results and discussion

### Comparison between the standard digestion (SD) and the in-solution buffer-free digestion (BFD) protocols

Both, the SD (**Fig. 1a**) and the in-solution BFD (24) (**Fig. 1b**) protocols, start with the S-alkylation of free cysteine residues by adding an excess of N-ethylmaleimide (NEM) or iodoacetamide (IAA). This step blocks the free thiol groups that can be present either because the RBD contains an odd number of cysteine residues or these groups were not quantitatively linked and thus remain partially free by a non-correct folding. At the same time, the alkylating agent added at the beginning of the protocol avoids artifacts due to the disulfide bond exchange during the subsequent steps (22). This could be more critical in the conventional protocol using a basic pH during tryptic digestion (29, 30). The use of a slightly acidic pH for trypsin digestion (pH 5.5-6.0) with BFD minimizes artificial modifications introduced during sample preparation such as scrambling due to the presence of free Cys in the analyzed protein. The S-alkylating agent introduce an artificial mass tag that facilitates the assignment when any Cys is partially free and differentiate them from species modified with natural thiol-blocking groups due to alkylating species present in the culture media (31).

**Fig. 1.**
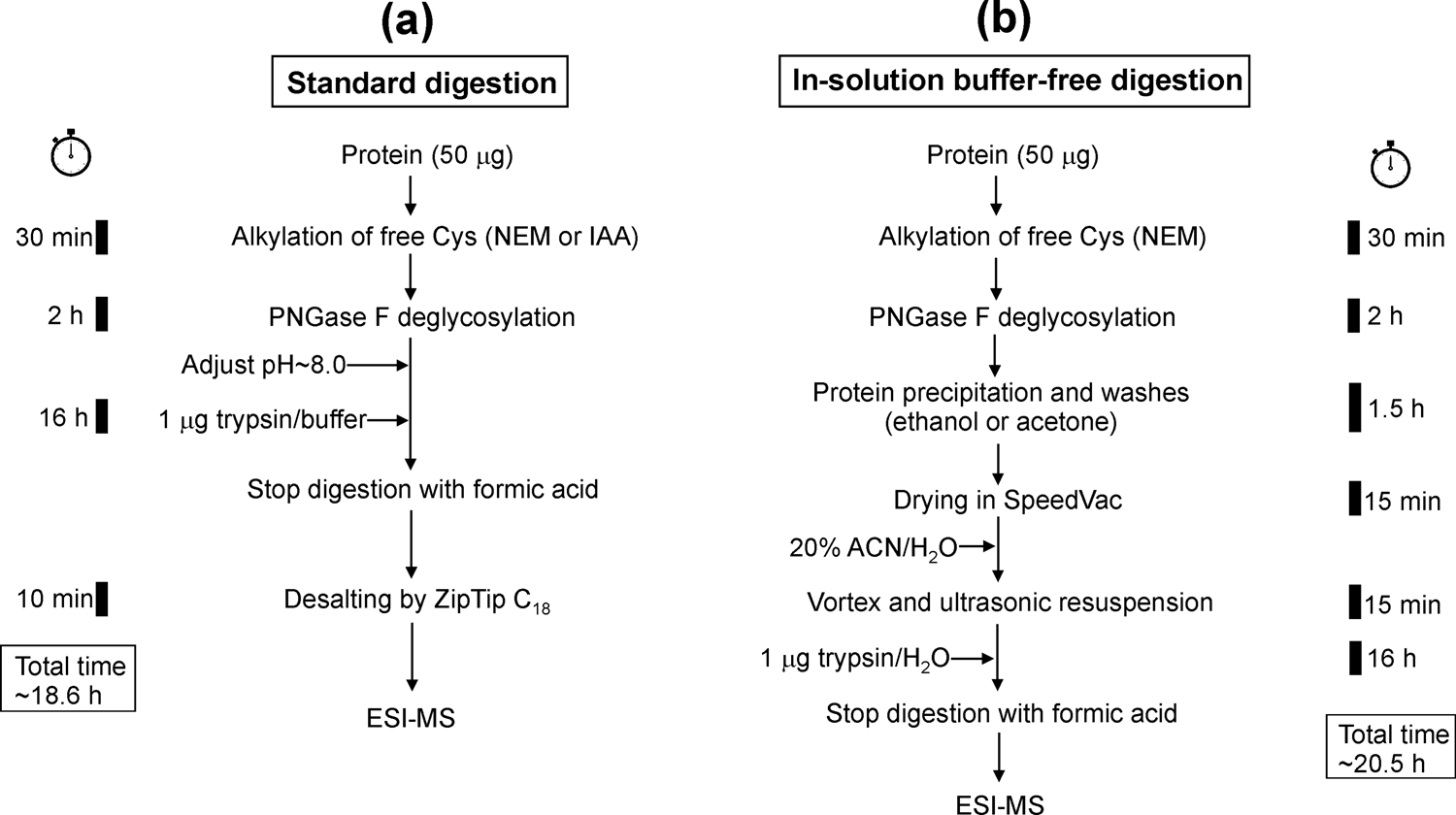
A comparison between the in-solution standard digestion **(a)** and buffer-free digestion (24) **(b)** protocols for the ESI-MS analysis of the tryptic digests. Black rectangles at the left and right sides of figure indicate the time required for the individual steps in each protocol. Square boxes at the bottom-left and bottom-right in the figure indicate the total time consumed for each protocol. NEM and IAA mean N-ethylmaleimide and iodoacetamide, respectively.

As a second step, both protocols comprise the deglycosylation with PNGase F of the recombinant RBDs and convert the fully glycosylated asparagines (Asn_331_/Asn_343_) into aspartic acids. This step also facilitates the detection and sequencing of two peptides (Phe_329_-Arg_346_) and (Ile_358_-Lys_378_) linked by an intermolecular disulfide bond between Cys_336_ and Cys_361_. For the particular cases of RBD_(333-527)_-C1 and RBD_(331-530)_-cmyc-Pp the peptide with the disulphide bond Cys_336_-Cys_361_ at the same time contains the N-terminal end of the protein. The identification of the disulfide bonds as well as the N-terminal sequencing of the protein are aspects inquired by regulatory agencies to develop well-characterized products according the ICHQ6B guidelines (18).

For the in-solution SD protocol (**Fig. 1a**), the pH of the solution is adjusted at basic pH and the deglycosylated RBD is digested with trypsin during 16 hours due to our interest to guarantee an efficient digestion. Also note that even after disulfide reduction this protein has been digested overnight by other authors (12, 32). Finally, the digestion is quenched by adding formic acid and the resultant tryptic peptides are desalted by using C_18_-ZipTips and eluted in a solution compatible with ESI-MS analysis.

For the in-solution BFD protocol (**Fig. 1b**), a desalting step is achieved at the protein level by conventional precipitation protocols using either cold acetone (33) or ethanol (34). Here, washing steps are included to minimize inorganic ions that may provoke adduct signals in the mass spectra. Protein resuspension is guaranteed by vigorous vortex and ultrasonic bath in 20% acetonitrile, before adding trypsin previously dissolved in water. There are no appreciable differences on both workflows (**Fig. 1a** and **1b**) respect to the processing time before MS analysis.

### Characterization of RBD_(319-541)_-HEK_A_3_ and RBD_(319-541)_-HEK proteins

*RBD_(319-541)_-HEK_A_3_* (**Table 1**) expressed in HEK-293T mammalian cell line has four disulfide bonds and a free cysteine residue (Cys_538_) located towards the C-terminal region of the protein. The high reactivity of Cys_538_ can be used for either site-directed chemical conjugation to highly immunogenic carrier proteins such as tetanus toxoid (35).

*RBD*(*_319-541_*)*-HEK_A_3_* was analyzed by SDS-PAGE in non-reducing conditions (**Fig. 2a**, lane 2) showing an intense and diffuse band at 33.3 kDa corresponding to the monomer with the heterogeneity of the N-glycosylation. Also, a band detected at 59.7 kDa representing approximately ∼13% was assigned to the dimer. After treatment with PNGase F, and analyzed under non-reducing conditions these bands migrated at 29.3 and 43.9 kDa (**Fig. 2a**, lane 3) confirming that *RBD_(319-541)_-HEK_A_3_* is *N*-glycosylated. The presence of *O*-glycosylation was not excluded because PNGase F does not hydrolyze *O*-glycans covalently linked to serine or threonine. When the same samples were analyzed by SDS-PAGE under reducing conditions only protein bands corresponding to the glycosylated monomer (**Fig. 2a**, lane 5) and the deglycosylated monomer (**Fig. 2a**, lane 6) were detected. No evidence of the dimer were observed suggesting that dimerization of the molecule was mediated by disulfide bonds and not due to an aggregation artifact.

**Fig. 2.**
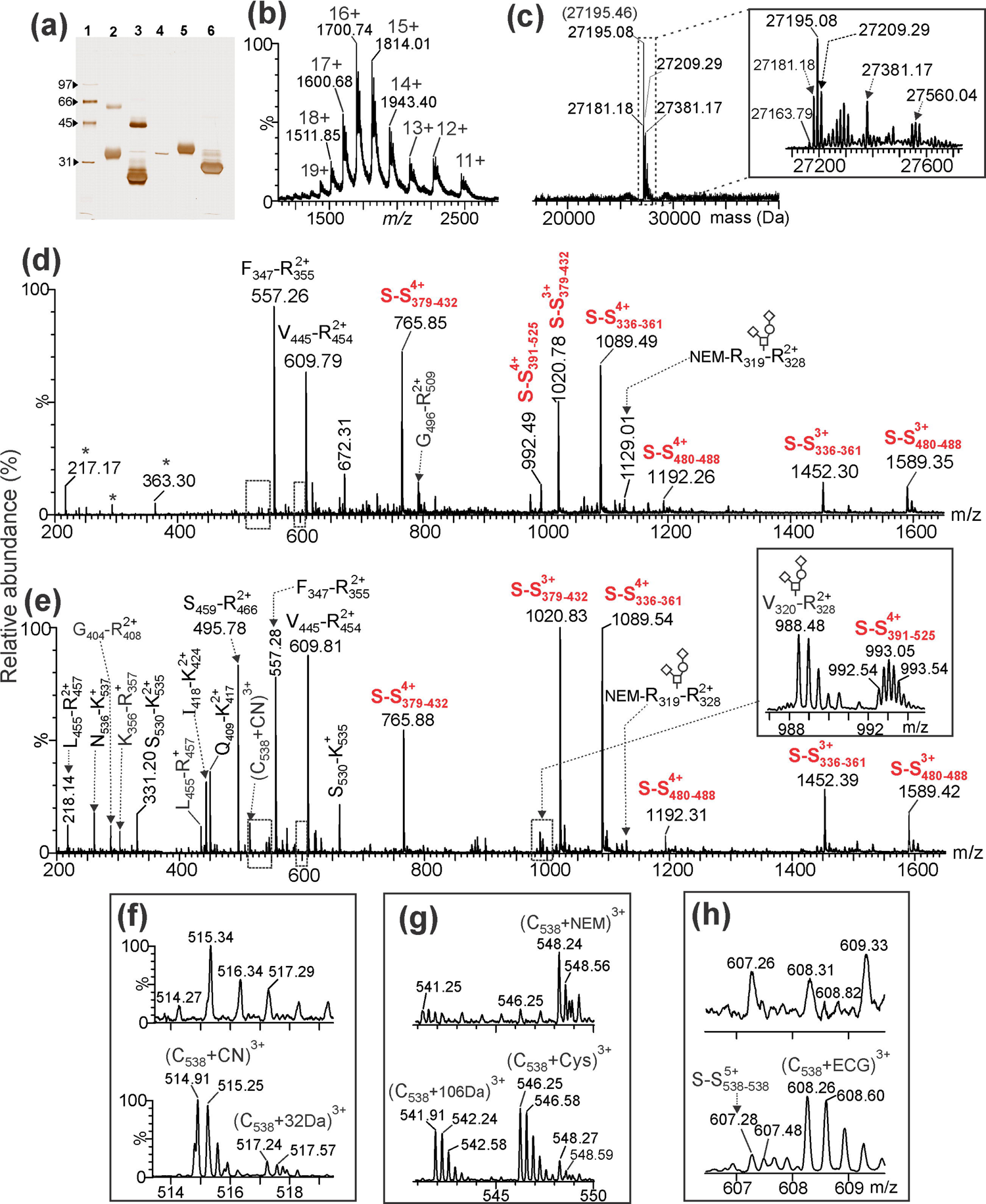
(a) SDS-PAGE analysis in reducing and non-reducing conditions of N-glycosylated and deglycosylated *RBD_(319-541)_-HEK_A_3_* and detected with silver staining. Lane 1: Molecular weight markers of low-range from 31 to 97 kDa (Bio-Rad). Lane 2-3: N-glycosylated and deglycosylated protein in non-reducing conditions detecting the monomer and a low-abundance (13%) dimer specie of *RBD_(319-541)_-HEK_A_3_*. Lane 4: Control of PNGase F used in the N-deglycosylation. Lane 5-6: N-glycosylated and deglycosylated protein in reducing conditions. **(b)** ESI-MS analysis of the *RBD_(319-541)_-HEK_A_3_* deglycosylated with PNGase F. **(c)** Resultant ESI-MS spectrum after deconvolution with MaxEnt v 1.0 software. The inset shown in **(c)** corresponds to the expanded ESI-MS spectrum in the range delimited by a broken line rectangle. The masses between parentheses indicate the expected molecular masses of the detected species. A detailed assignment of this ESI-MS spectrum is shown in **Table 2**. The ESI-MS spectra shown in **(d)** an **(e)** correspond to the ESI-MS analysis of the resultant tryptic peptides of *RBD_(319-541)_-HEK_A_3_* digested with trypsin following the SD and in-solution BFD (with ethanol precipitation) protocols shown in **Fig. 1(a)** and **(b)**. Asterisks in **(d)** correspond to background signals, not assigned to tryptic peptides. The inset shown in **(e)** corresponds to an expanded region where the O-glycosylated N-terminal end peptide (Val^320^-Arg^328^+[HexNAc:Hex:NeuAc_2_])^2+^ and two disulfide bonded peptides (assigned as S-S_391-525_) were detected. Monosaccharide symbols follow the SNFG system (60) and the O-glycans structures as previously reported (36). The upper and lower mass spectra shown in **(f)**, **(g)** and **(h)** correspond to expanded regions of the ESI-MS spectra shown in **(d)** and **(e)**, respectively. A detailed assignment for all tryptic peptides in this figure is summarized in **Table 3**.

**Table 2.**
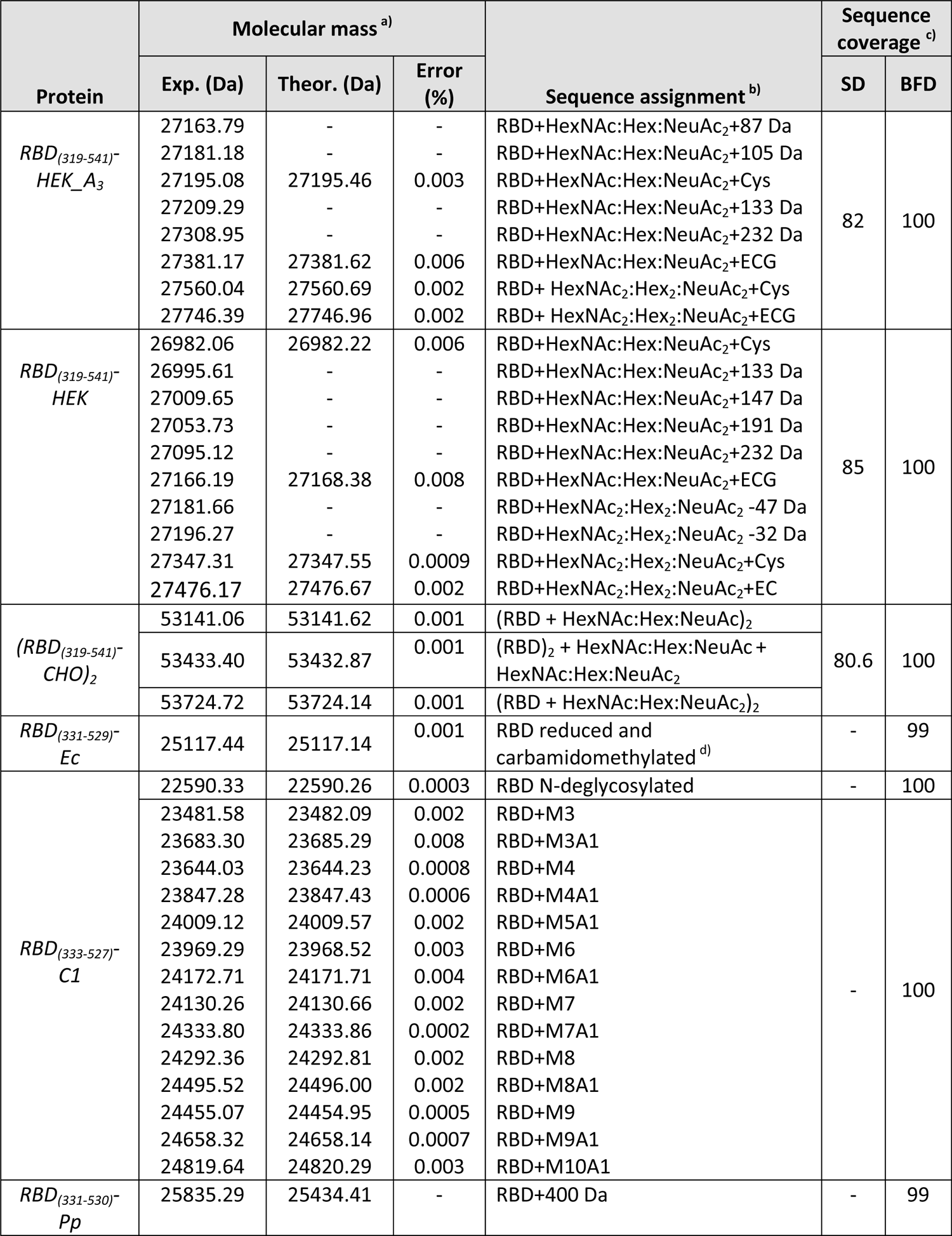
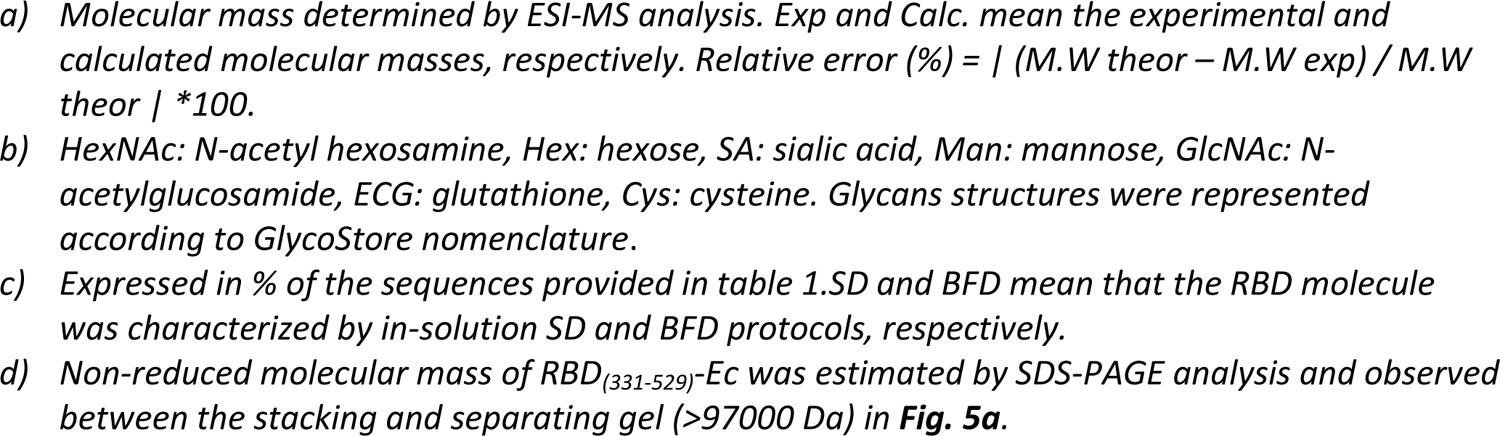
Summary of the ESI-MS analysis of in-solution SD and BFD protocol and sequence coverage of the RBD characterized in this work.

| Protein | Molecular mass <sup>a)</sup> |  |  | Sequence assignment <sup>b)</sup> | Sequence coverage <sup>c)</sup> |  |
| --- | --- | --- | --- | --- | --- | --- |
|  | Exp. (Da) | Theor. (Da) | Error (%) |  | SD | BFD |
| *RBD<sub>(319-541)</sub>-HEK_A<sub>3</sub>* | 27163.79 | - | - | RBD+HexNAc:Hex:NeuAc<sub>2</sub>+87 Da | 82 | 100 |
|  | 27181.18 | - | - | RBD+HexNAc:Hex:NeuAc<sub>2</sub>+105 Da |  |  |
|  | 27195.08 | 27195.46 | 0.003 | RBD+HexNAc:Hex:NeuAc<sub>2</sub>+Cys |  |  |
|  | 27209.29 | - | - | RBD+HexNAc:Hex:NeuAc<sub>2</sub>+133 Da |  |  |
|  | 27308.95 | - | - | RBD+HexNAc:Hex:NeuAc<sub>2</sub>+232 Da |  |  |
|  | 27381.17 | 27381.62 | 0.006 | RBD+HexNAc:Hex:NeuAc<sub>2</sub>+ECG |  |  |
|  | 27560.04 | 27560.69 | 0.002 | RBD+ HexNAc<sub>2</sub>:Hex<sub>2</sub>:NeuAc<sub>2</sub>+Cys |  |  |
|  | 27746.39 | 27746.96 | 0.002 | RBD+ HexNAc<sub>2</sub>:Hex<sub>2</sub>:NeuAc<sub>2</sub>+ECG |  |  |
| *RBD<sub>(319-541)</sub>-HEK* | 26982.06 | 26982.22 | 0.006 | RBD+HexNAc:Hex:NeuAc<sub>2</sub>+Cys | 85 | 100 |
|  | 26995.61 | - | - | RBD+HexNAc:Hex:NeuAc<sub>2</sub>+133 Da |  |  |
|  | 27009.65 | - | - | RBD+HexNAc:Hex:NeuAc<sub>2</sub>+147 Da |  |  |
|  | 27053.73 | - | - | RBD+HexNAc:Hex:NeuAc<sub>2</sub>+191 Da |  |  |
|  | 27095.12 | - | - | RBD+HexNAc:Hex:NeuAc<sub>2</sub>+232 Da |  |  |
|  | 27166.19 | 27168.38 | 0.008 | RBD+HexNAc:Hex:NeuAc<sub>2</sub>+ECG |  |  |
|  | 27181.66 | - | - | RBD+HexNAc<sub>2</sub>:Hex<sub>2</sub>:NeuAc<sub>2</sub> -47 Da |  |  |
|  | 27196.27 | - | - | RBD+HexNAc<sub>2</sub>:Hex<sub>2</sub>:NeuAc<sub>2</sub> -32 Da |  |  |
|  | 27347.31 | 27347.55 | 0.0009 | RBD+HexNAc<sub>2</sub>:Hex<sub>2</sub>:NeuAc<sub>2</sub>+Cys |  |  |
|  | 27476.17 | 27476.67 | 0.002 | RBD+HexNAc<sub>2</sub>:Hex<sub>2</sub>:NeuAc<sub>2</sub>+EC |  |  |
| *(RBD<sub>(319-541)</sub>-CHO)<sub>2</sub>* | 53141.06 | 53141.62 | 0.001 | (RBD + HexNAc:Hex:NeuAc)<sub>2</sub> | 80.6 | 100 |
|  | 53433.40 | 53432.87 | 0.001 | (RBD)<sub>2</sub> + HexNAc:Hex:NeuAc + HexNAc:Hex:NeuAc<sub>2</sub> |  |  |
|  | 53724.72 | 53724.14 | 0.001 | (RBD + HexNAc:Hex:NeuAc<sub>2</sub>)<sub>2</sub> |  |  |
| *RBD<sub>(331-529)</sub>-Ec* | 25117.44 | 25117.14 | 0.001 | RBD reduced and carbamidomethylated <sup>d)</sup> | - | 99 |
| *RBD<sub>(333-527)</sub>-C1* | 22590.33 | 22590.26 | 0.0003 | RBD N-deglycosylated | - | 100 |
|  | 23481.58 | 23482.09 | 0.002 | RBD+M3 | - | 100 |
|  | 23683.30 | 23685.29 | 0.008 | RBD+M3A1 |  |  |
|  | 23644.03 | 23644.23 | 0.0008 | RBD+M4 |  |  |
|  | 23847.28 | 23847.43 | 0.0006 | RBD+M4A1 |  |  |
|  | 24009.12 | 24009.57 | 0.002 | RBD+M5A1 |  |  |
|  | 23969.29 | 23968.52 | 0.003 | RBD+M6 |  |  |
|  | 24172.71 | 24171.71 | 0.004 | RBD+M6A1 |  |  |
|  | 24130.26 | 24130.66 | 0.002 | RBD+M7 |  |  |
|  | 24333.80 | 24333.86 | 0.0002 | RBD+M7A1 |  |  |
|  | 24292.36 | 24292.81 | 0.002 | RBD+M8 |  |  |
|  | 24495.52 | 24496.00 | 0.002 | RBD+M8A1 |  |  |
|  | 24455.07 | 24454.95 | 0.0005 | RBD+M9 |  |  |
|  | 24658.32 | 24658.14 | 0.0007 | RBD+M9A1 |  |  |
|  | 24819.64 | 24820.29 | 0.003 | RBD+M10A1 |  |  |
| *RBD<sub>(331-530)</sub>-Pp* | 25835.29 | 25434.41 | - | RBD+400 Da | - | 99 |

**Table 3.**
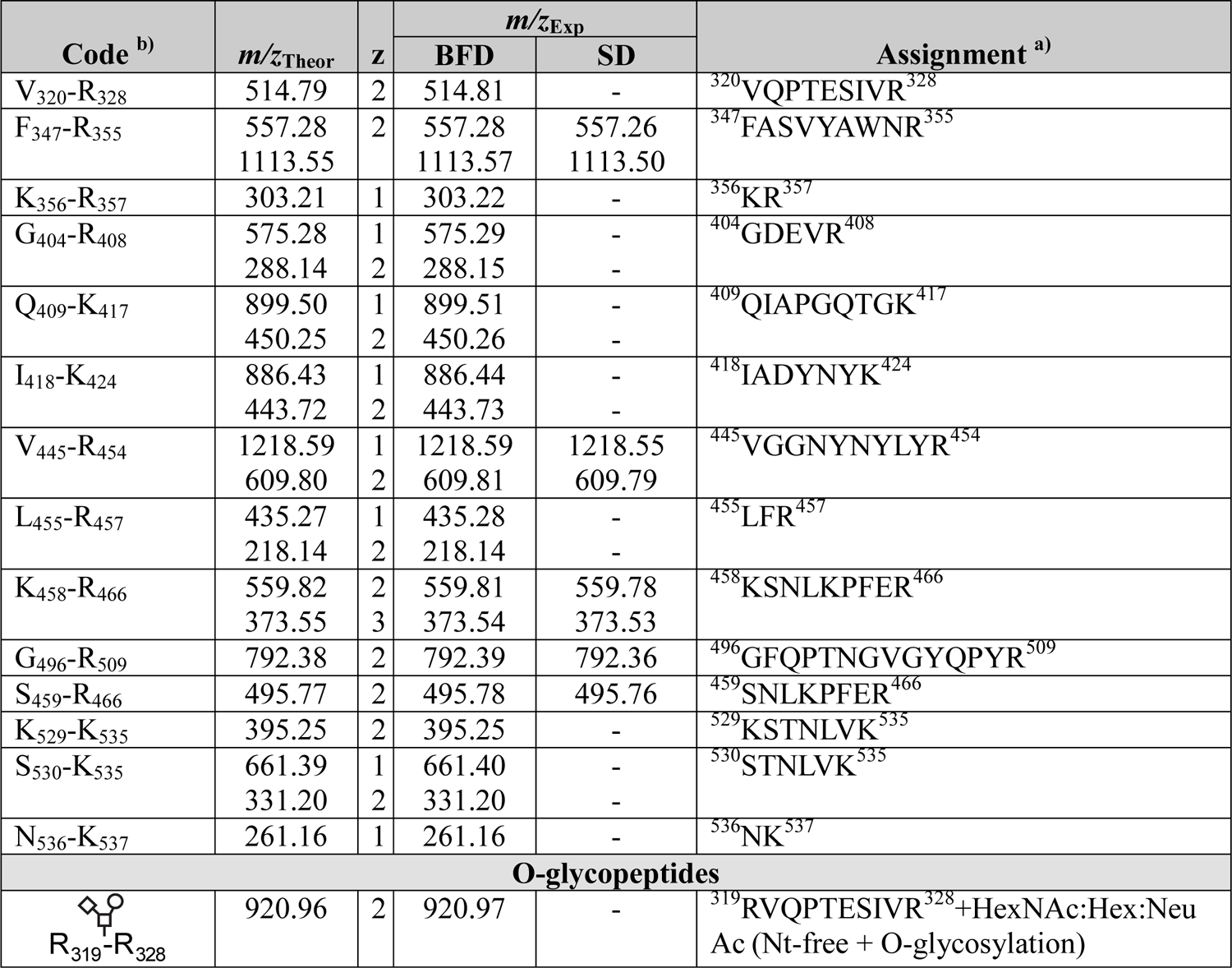
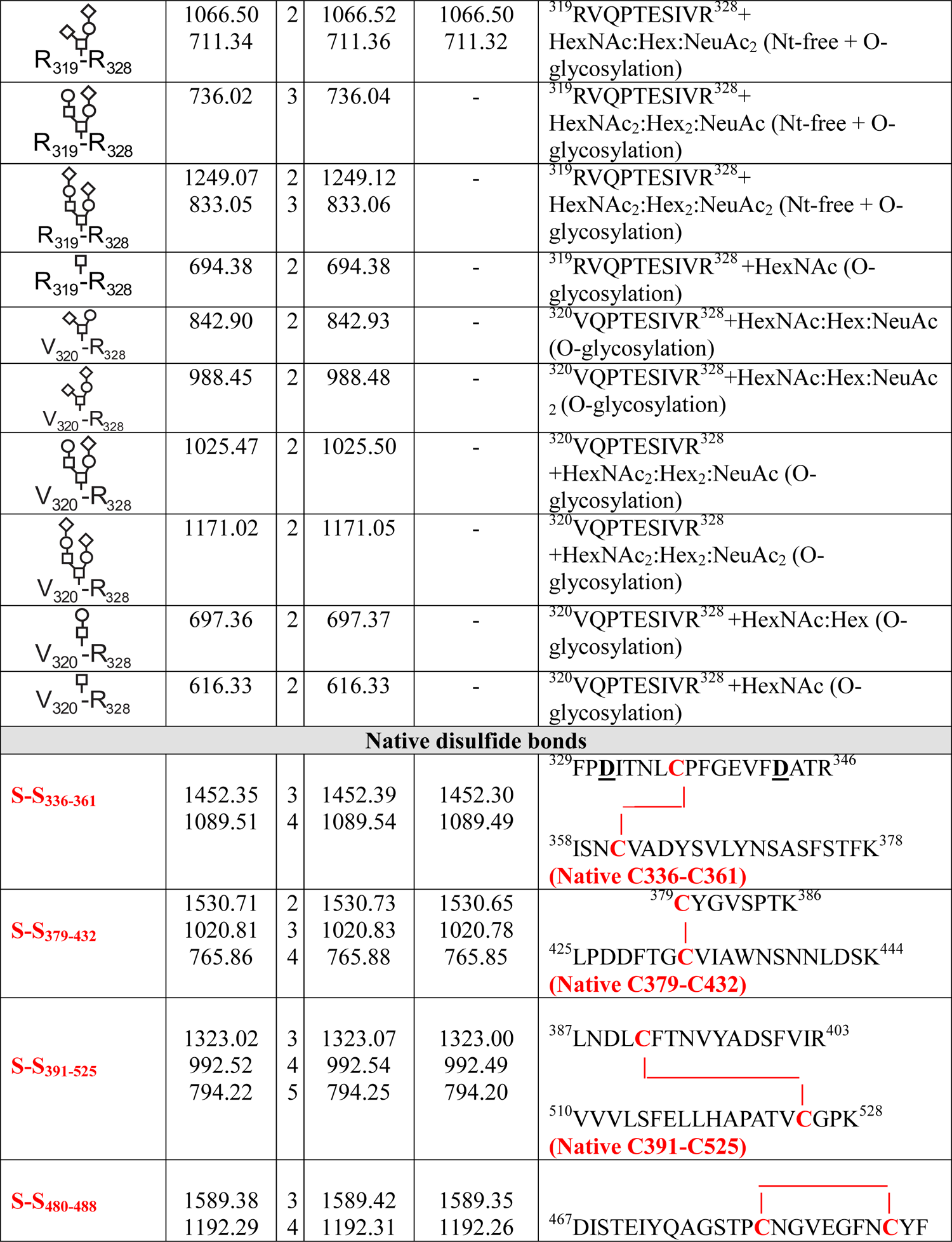
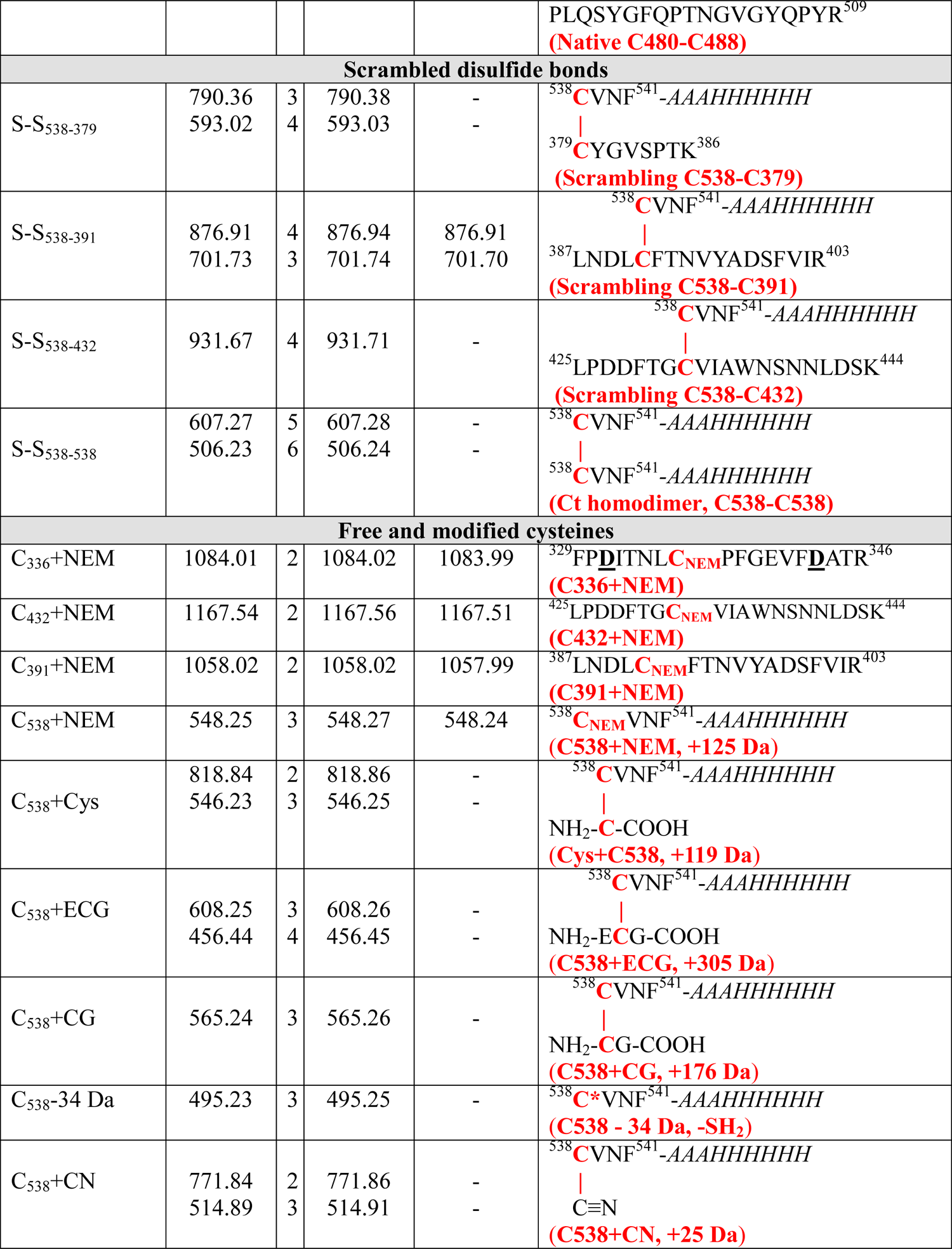
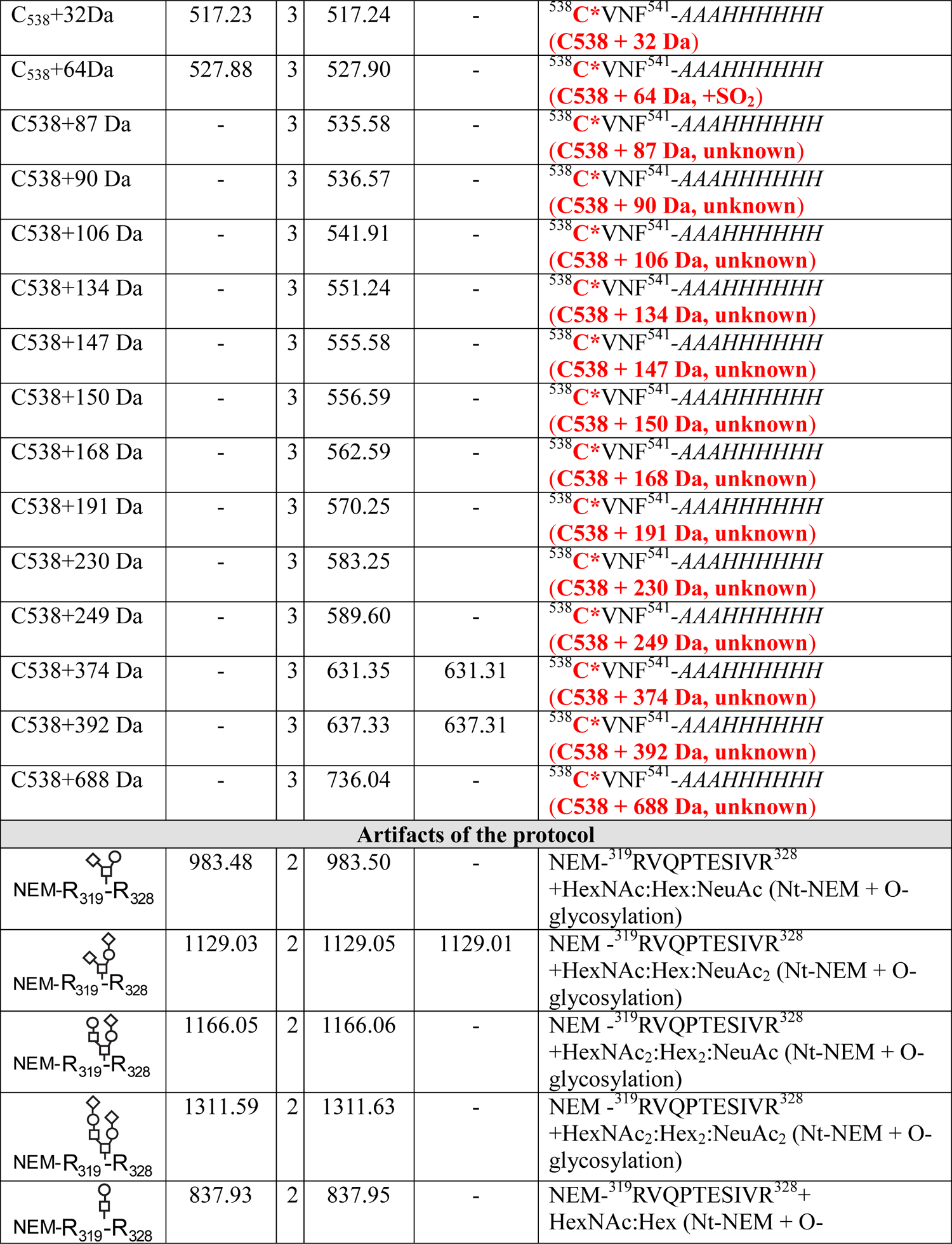
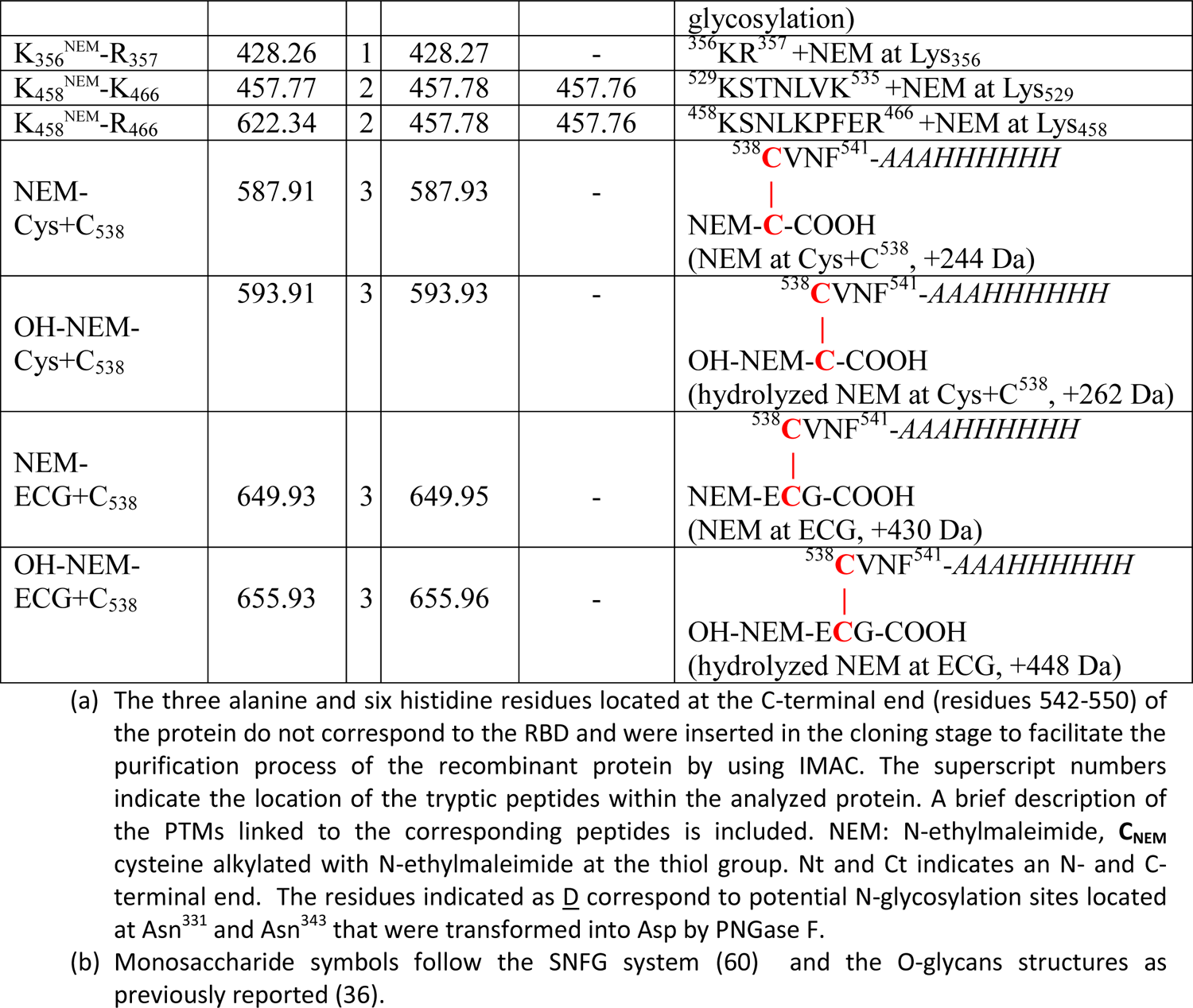
Summary of the 100% sequence coverage assignment by ESI-MS of the tryptic digestion using the in-solution buffer-free (BFD) and 82% by the standard digestion (SD) protocol of *RBD_(319-541)_-HEK_A_3_* expressed in HEK293T.

To confirm the integrity, the N-deglycosylated protein was analyzed by ESI-MS (**Fig. 2b**) and showed intense multiply-charged signals corresponding to z=11+ to 19+ of the protein. The deconvoluted ESI-MS spectrum (**Fig. 2c**) shows the most intense signal with molecular mass of 27195.08 Da that agreed with the expected mass (27195.46 Da) considering the *N*-deglycosylated monomer of *RBD_(319-541)_-HEK_A_3_*, cysteinylated and *O*-glycosylated with HexNAc:Hex:NeuAc_2_ (**Table 2**). Other groups that also expressed RBD molecules in HEK-293 with an odd number of cysteine residues reported cysteinylation (32, 35). O-glycosylation has been reported for the native RBD of SARS-Cov-2 (21, 36) as well as for several RBD expressed in mammalian cells (32, 35).

Also, of other signals observed in **Fig. 2c** (see inset) and summarized in **Table 2** suggest the presence of other modified species of the N-deglycosylated *RBD_(319-541)_-HEK_A_3_*. Separately, the *N*-deglycosylated protein was digested in-solution with trypsin by using the SD (**Fig. 1a**) and BFD (**Fig. 1b**) protocols and the resultant ESI-MS spectra are shown in **Fig. 2d** and **Fig. 2e**, respectively. The sequence assignment based on the agreement between the expected and experimental *m/z* of tryptic peptides are summarized in **Table 3**. The four disulfide bonds present in the native RBD of S protein of SARS-CoV-2 were identified by both protocols (**Fig. 2d and 2e**) and confirmed by MS/MS analysis (**Fig. S1a-S1d**).

In the SD protocol only the N-terminal peptide R_319_-R_328_ containing HexNAc:Hex:NeuAc_2_ was detected (**Fig. 2d**, **Table 3**), presumably linked to either at Thr_323_ or Ser_325_ according to previous reports (32, 36). However, in the in-solution BFD protocol the peptides R_319_-R_328_ and V_320_-R_328_ linked to HexNAc; HexNAc:Hex; HexNAc:Hex:NeuAc; HexNAc:Hex:NeuAc_2_; HexNAc_2_:Hex_2_:NeuAc and HexNAc_2_:Hex_2_:NeuAc_2_ were detected (**Fig. 2e**, **Table 3**). Five out of the six *O*-glycans structures were detected only by the in-solution BFD protocol and these six *O*-glycans structures agree very well with the previous reports of *O*-glycosylation of Thr_323_/Ser_325_ in the SARS-CoV-2 spike protein (21, 36). MS/MS spectra of these *O*-glycopeptides confirmed this assignment (**Fig. S2a-S2c**) by showing intense neutral losses of glycans from the precursor ions fragmented by CID according to previous reports (37).

Full-sequence of *RBD_(319-541)_-HEK_A_3_* was verified by using in-solution BFD protocol, while using the SD protocol 82% of sequence coverage was achieved (**Table 2**).

Several signals in the low-mass region (*m/z* 200-700) were exclusively detected when *RBD_(319-541)_-HEK_A_3_* was analyzed by the BFD protocol and they were assigned to short and hydrophilic internal peptides (^356^KR^357^, ^536^NK^537^, ^455^LFR^457^, ^404^GDEVR^408^, ^409^QIAPGQTGK^417^, ^418^IADYNYK^424^, ^529^KSTNLVK^535^ and ^530^STNLVK^535^, **Table 3**). These peptides represent the 18% of the *RBD_(319-541)_-HEK_A_3_* sequence. Most of them (^356^KR^357^, ^455^LFR^457^, ^409^QIAPGQTGK^417^, ^418^IADYNYK^424^, ^529^KSTNLVK^535^ and ^530^STNLVK^535^) were not detected by Arbeitman *et al.* (12) when the same RBD protein expressed in *P. pastoris* and HEK-293T cell line was digested with a protocol similar to the in-solution SD protocol and analyzed by MALDI-MS. Two of them (^404^GDEVR^408^ and ^536^NK^537^) were only detected in the RBD expressed in HEK-293T, probably included in the sequences of more hydrophobic peptides containing missed cleavages sites.

The C-terminal peptide with the C_538_ alkylated with NEM (^538^CVNF^541^-*AAAHHHHHH, m/z* 548.24, 3+, **Fig. 2g** and **Table 3**) was detected by both protocols (**Fig. 1a-b**). It confirmed that a fraction of this RBD contains an unpaired free C_538_ residue. However, the low-intensity of the signal assigned to the C-terminal peptide with a C_538_ alkylated with NEM (*m/z* 548.27, 3+; **Fig. 2g** and **Table 3**) when BFD protocol was applied suggested that Cys_538_ should be modified with other chemical groups.

Cyanylation (*m/z* 541.91, 3+; (C_538_+CN)^3+^, **Fig. 2f**), cysteinylation (*m/z* 546.25, 3+; (C_538_+Cys)^3+^, **Fig. 2g**) and glutathionylation (*m/z* 608.26, 3+; (C_538_+ECG)^3+^, **Fig. 2h**) of the unpaired Cys_538_ in the C-terminal peptide of *RBD_(319-541)_-HEK_A_3_* was detected exclusively when using the BFD protocol. The assignment of these modified peptides was confirmed by MS/MS analysis (**Fig. 3a-3c**). A signal detected at *m/z* 565.26, 3+ was also only observed when *RBD_(319-541)_-HEK_A_3_* was analyzed by BFD (**Table 3**). MS/MS analysis demonstrated that it corresponded to the same C-terminal peptide (C_538_+CG)^3+^ with the C_538_ linked to a truncated variant of glutathion (**Fig. S3a**).

**Fig. 3.**
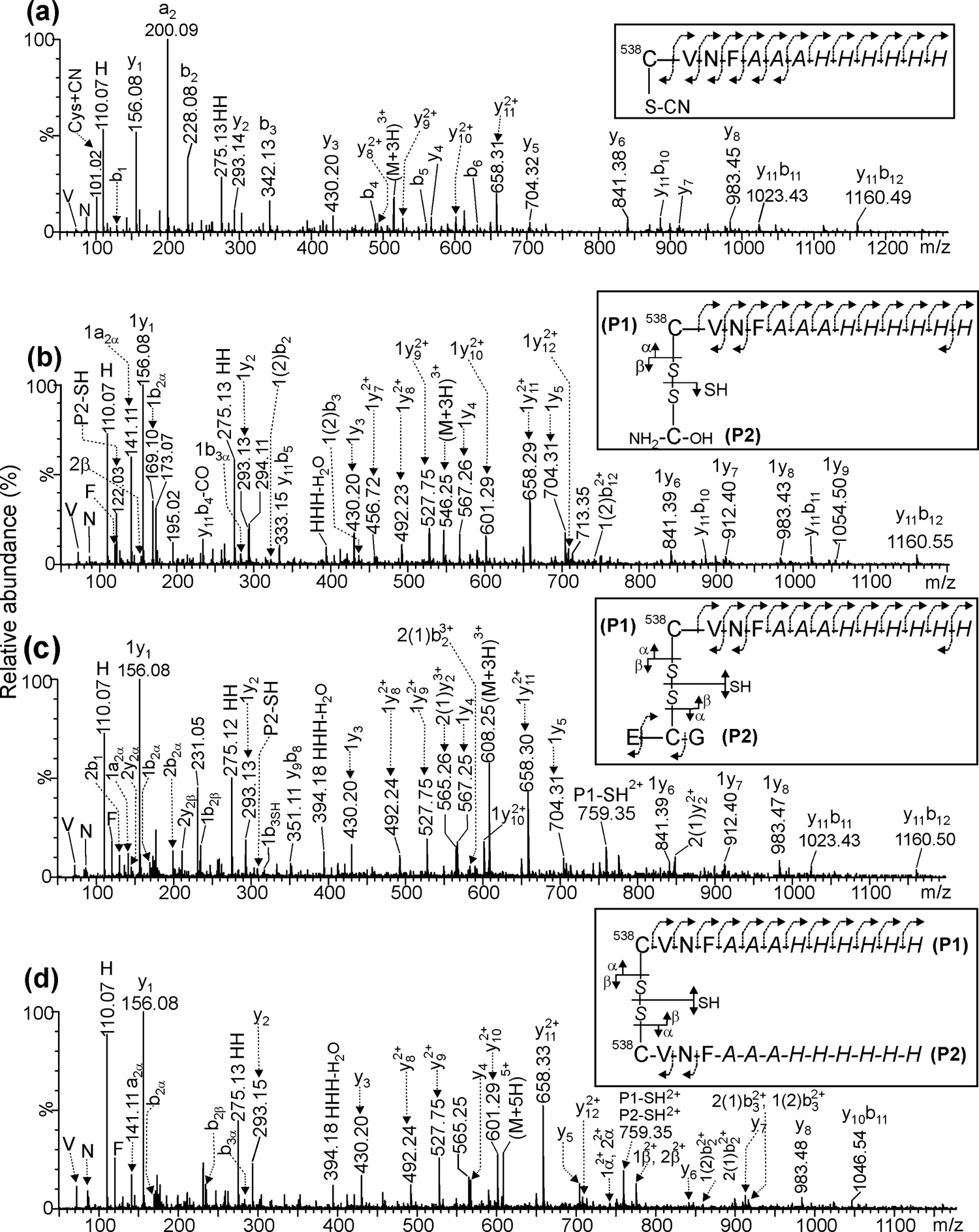
ESI-MS/MS spectra of peptides containing modified cysteine by **(a)** cyanylation, **(b)** cysteinylation, **(c)** glutathionylation and **(d)** dimerization containing Cys_538_ within the C-terminal end peptide of two linked subunits. The nomenclature of fragment ions is in agreement with the proposed by Mormann *et al* (61).

The alkylation with NEM, inserted in our protocols (**Fig. 1a, b**), transformed the hydrophilic C-terminal peptide (containing the unpaired C_538_) in a more hydrophobic species and in consequence it was detected even using the SD protocol. On the contrary, the remaining Cys-capping modifications (31) mentioned above (cyanylation, glutathionylation, cysteinylation and truncated glutathionylation) did not increase the hydrobicity of the C-terminal peptide sufficiently to be retained by ZipTip-C_18_ and they were detected exclusively when in-solution BFD was applied.

Signals detected at *m/z_Exp_* 517.24, 3+ (**Fig. 2f**) and *m/z_Exp_* 541.91, 3+ (**Fig. 2g**) were assigned as (C_538_+32 Da)^3+^ and (C_538_+106 Da)^3+^, corresponding to the C-terminal peptide with C_538_ linked to modifying groups of unknown chemical nature. These signals were only detected when in-solution BFD was applied to the characterization of *RBD_(319-541)_-HEK_A_3_*. We also found thirteen other different variants of the C-terminal peptide (confirmed by MS/MS, see **Fig. S3**) that were not assigned to a defined chemical structure of Cys^538^ (see **Table 3**).

Opossed to the thesis proposing that oxidoreductase-mediated protein disulfide bonding with free cysteine or glutathione in the lumen of endoplasmic reticulum (38–40) as the source of these modifications, Zhong et al. have demonstrated that these modifications are generated outside mammalian cells and are sensitive to the culture medium composition (31). Interestingly, the authors also proposed the addition of chemically modified drugs to the culture medium to obtain homogeneous drug conjugates in a single step avoiding the need for an uncapping step prior to conjugation (31).

Cysteinylation at Cys_538_ has been reported by other authors [31, 34], but to our knowledge the other modifying groups (**Table 3**) have not previously been reported for RBD. The species with Cys_538_ modifications and *O*-glycoforms detected at protein level (**Table 2**) were further confirmed at tryptic peptide level by the in-solution BFD (**Table 3**).

The usage of culture media with defined composition and a well-characterized downstream process would avoid unexpected modifications of free cysteine residues (41, 42), otherwise, the structural assignment for these PTMs would be more difficult according to our knowledge (43).

Although Cys-capping modifications protect the molecule from aggregation and scrambling mediated by inter- and intra-molecular disulfide bonds, respectively, it need to be adressed if the final outcome is to use the unpaired Cys for further modification as for example drug conjugation process (44, 45). Another issue also to be adressed is the potential protein heterogeneity if the final intention is the use of the dimer molecule through disulphide bonds (38, 39, 46, 47).

A low-intensity signal at *m/z_Exp_* 607.28, 5+ and assigned to (S-S^5+^) in **Fig. 2h**, was exclusively detected when in-solution BFD protocol was applied. It suggests that a minor fraction of this molecule is a dimer mediated by an intermolecular disulfide bond between two Cys_538_ residues. MS/MS of this signal confirmed this assignment (**Fig. 3d**). This result match with SDS-PAGE of *RBD_(319-541)_-HEK_A_3_* ran under reducing and non reducing conditions showing that a dimer of this molecule repesents 13% of this preparation (**Fig. 2a**).

The presence of three low abundance scrambling variants (C_538_-C_379_, C_538_-C_391_, C_538_-C_432_) and the homodimer (C_538_-C_538_) of this molecule agrees with the presence of a free Cys_538_ detected in this preparation (**Table 3**). All the previous scrambled species were detected by using the in-solution BFD protocol, while only the scrambling C_538_-C_391_ was detected by the SD protocol. Also, a low-abundance population of the protein with free C_336_, C_391_, C_432_ and C_538_ was detected by both protocols. All the above-mentioned assignments of scrambled and free Cys variants were confimed by the MS/MS spectra (**Fig. S4-S5**). The presence of an unpaired Cys residues may also promote disulfide exchange (22) and in consequence generates low-abundance scrambling variants of the desired molecule.

Our results indicate that Cys reduction and S-alkylation of the RBD protein before MS analysis is not convenient as important information is lost. The most striking results obtained with the BFD protocol are the detection of the disulfide-containing peptides (including low-abundance scrambled variants) and the finding of several modifications linked to free cysteines that probably, some of them would be missed if reduction of disulfides takes place during sample preparation.

The analysis of the same gene construct (*RBD_(319-541)_-HEK)* for the expression in HEK-293T of the same protein without the C-terminal spacer arm of three alanines (**Table 1**) by in-solution SD and BFD protocol (**Fig. S6, Table 2 and S1**) yields similar results to that described here for *RBD_(319-541)_-HEK_A_3_*, at protein and peptide level (**Fig. 2, Table 2 and 3**). Full-sequence coverage was achieved in the analysis of *RBD_(319-541)_-HEK* by using in solution BFD protocol while using the SD protocol 85% was achieved (**Table 2**). C-terminal peptide containing the six histidine tail was only detected by using BFD protocol (data not shown).

### Characterization of (RBD_(319-541)_-CHO)_2_

The RBD dimer *(RBD_(319-541)_-CHO)_2_* resulting from an intermolecular disulfide bond Cys_538_-Cys_538_ was originally obtained as a by-product during the attempt to obtain *RBD_(319-541)_-CHO.* Unpaired Cys^538^ was introduced in order to use it for site selective conjugation to tetanus toxoid (35). The increased immunogenicity of RBD-dimer (9, 48) promoted their use in at least two vaccines currently in clinical trials.

The *(RBD_(319-541)_-CHO)_2_* protein non-treated (**lane 2**, **Fig. 4a**) and treated (**lane 3**, **Fig. 4b**) with PNGase F and analyzed by SDS-PAGE under reducing conditions only the presence of a glycosylated and deglycosylated monomer, respectively, were observed. When the same samples were analyzed by non-reducing conditions the glycosylated (**lane 4**, **Fig. 4a**) and the deglycosylated (**lane 5**, **Fig. 4a**) dimer were observed. This result confirmed the dimeric and N-glycosylated nature of *(RBD_(319-541)_-CHO)_2_*.

**Fig. 4.**
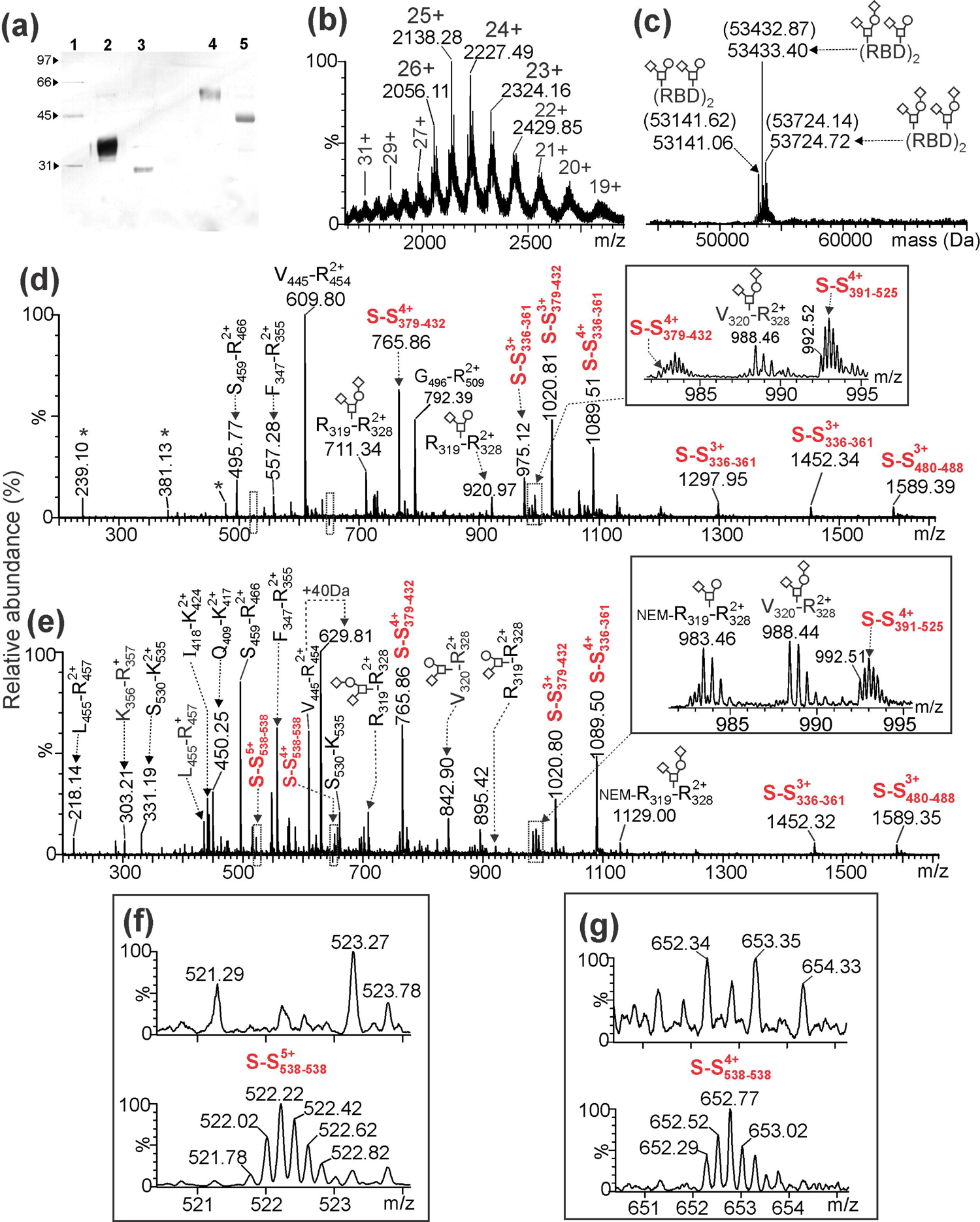
(a) SDS-PAGE analysis in reducing and non-reducing conditions of N-glycosylated and deglycosylated *(RBD_(319-541)_-CHO)_2_* and detected with silver staining. Lane 1: Molecular weight markers of low-range from 31 to 97 kDa (Bio-Rad). Lane 2-3: N-glycosylated and deglycosylated protein in reducing conditions detecting the reduced monomer. Lane 4-5: N-glycosylated and deglycosylated protein in non-reducing conditions detecting the dimer species [*(RBD_(319-541)_-CHO)_2_*]. **(b)** ESI-MS spectrum of a dimeric RBD deglycosylated with PNGase F. **(c)** Deconvolution of the ESI-MS spectrum shown in **(b)** reveals the presence of the three major *O*-glycoforms of *(RBD_(319-541)_-CHO)_2_*. Between parentheses the expected molecular masses of the different *O*-glycoforms are shown. (RBD)_2_ represents an abbreviated form for referring to the *(RBD_(319-541)_-CHO)_2_* molecule. Monosaccharide symbols follow the SNFG system (60) and the O-glycans structures as previously reported (36). The ESI-MS spectra shown in **(d)** and **(e)** correspond to the *(RBD_(319-541)_-CHO)_2_* digested with trypsin following the SD and in-solution BFD (precipitated with acetone) protocol, respectively. Asterisks in **(d)** correspond to background signals, not assigned to tryptic peptides and **(S-S)^n+^** were peptides containing a disulfide bonds between the described cysteines. The insets shown in **(d)** and **(e)** correspond to the expanded regions of the mass spectra (*m/z* 981.5-995.5) delimited by rectangles with broken lines showing the O-glycosylated peptides and two disulfide bond peptides (assigned as S-S_391-525_^4+^ and S-S_379-432_^4+^). The upper- and lower-mass spectra shown in **(f)** and **(g)** correspond to two expanded regions (*m/z* 520.4-524.1 and *m/z* 650.5-655.2) of the ESI-MS spectra shown in **(d)** and **(e)**, respectively. A detailed assignment for all tryptic peptides in this figure is summarized in **Table S2**.

The ESI-MS spectrum of the PNGase F deglycosylated dimer showed the typical multiply-charged ions and suggested certain heterogeneity in the molecule (**Fig. 4b**). It was more easily appreciated after the deconvolution process (**Fig. 4c**) due to the presence of three major signals corresponding to the three combinations of two short *O*-glycan chains linked to the dimer as indicated in **Fig. 4c** (21). The good agreement observed between the expected and experimental masses for all these *O*-glycoforms of *(RBD_(319-541)_-CHO)_2_* is summarized in **Table 2.** ESI-MS analysis of the *(RBD_(319-541)_-CHO)_2_* (**Fig. 4c**) suggest that this molecule is more homogeneous than the monomers *RBD_(319-541)_-HEK_A_3_ and RBD_(319-541)_-HEK,* probably because Cys_538_ is fully compromised in the intermolecular disulfide bond and is not linked to other blocking groups present in the culture media.

The N-deglycosylated protein was digested with trypsin by using the in-solution SD and BFD protocols and the resultant ESI-MS spectra are shown in **Fig. 4d** and **4e**, respectively. Full-sequence coverage was achieved for the in-solution BFD protocol while using the SD protocol only 80.6% of the sequence was verified (**Table 2**). The assignments for all tryptic peptides generated by both protocols are summarized in **Table S2**.

The four disulfide bonds present in the native RBD of SARS-CoV-2 were detected by applying both protocols (**Fig 4d** and **Fig 4e**). O-glycosylated N-terminal peptides (R_319_-R_328_ and V_320_-R_328_) with O-glycosylation sites located at Thr_323_/Ser_325_ residues (21) were detected with appreciable intensities (*m/z_Exp_* 711.34, 3+; 842.90, 2+ and 988.44, 2+ in **Fig 4d** and **Fig 4e**). The mass shift provoked by these *O*-glycans observed for the N-deglycosylated protein (**Fig 4c**) agreed with the one observed at the peptide level (**Fig 4d** and **4e**). Additionally, two low-intensity signals at *m/z_Exp_* 616.33, 2+ and 697.36, 2+ assigned to peptide V_320_-R_328_ linked to HexNAc and HexNAc:Hex were detected only by in-solution BFD protocol (**Table S2**).

The most striking differences between both ESI-MS spectra (**Fig 4d** and **Fig 4e**) are observed in the low-mass region where short and hydrophilic peptides [L_455_-R_457_ (*m/z_Exp_* 218.14, 2+), G_404_-R_408_ (*m/z_Exp_* 288.14, 2+), K_356_-R_357_ (*m/z_Exp_* 303.21, 1+) and S_530_-K_535_ (*m/z_Exp_* 331.19, 2+)] were only detected by applying the in-solution BFD protocol.

The ESI-MS signals that confirm the dimeric nature of *(RBD_(319-541)_-CHO)_2_* are the corresponding to the peptide [C_538_-H_547_]-S-S-[C_538_-H_547_] containing Cys_538_ and Cys_538_ linked by intermolecular disulfide bond (*m/z_Exp_* 522.02, 5+ and *m/z_Exp_* 652.29, 4+ in **Fig. 4f** and **Fig. 4g**). These signals that also enabled the verification of the C-terminal end of this molecule were exclusively detected by applying the in-solution BFD protocol. Probably, the presence of twelve histidine residues in the structure of [C_538_-H_547_]-S-S-[C_538_-H_547_] (assigned as S_538_-S_538_ and S_538_-S_538_ and in **Fig. 4e** and **4f**, respectively) makes difficult its retention in the C_18_-ZipTip during the desalting step when the in-solution SD protocol is applied. The verification of the C-terminal end of proteins is a very important aspect included in the ICHQ6B guidelines and requested by regulatory authorities (18).

### Characterization of RBD_(331-529)_-Ec

The expression of RBD as inclusion bodies in *E. coli* constitutes a challenge due to the presence of eight cysteines and the absence of the native N-glycosylation required for the appropriate expression and folding of the molecule. The non-correctly folded RBD is not useful for a vaccine against SARS-CoV-2 because a tridimensional structure identical to the native protein is required to generate upon vaccination neutralizing antibodies recognizing conformational epitopes (19, 20) in the S protein. For this reason the detection of non-native disulfide bonds is of tremendous importance for the development of biologically active molecules and well-characterized products (18).

SDS-PAGE analysis under reducing conditions (**Fig. 5a**, lane 2) of *RBD_(331-529)_-Ec* shows a band that migrates with an estimated molecular mass of 27.3 kDa. The good agreement between the expected (25117.14 Da) and the experimental (25117.44 Da) molecular masses for the reduced and S-alkylated protein determined by ESI-MS analysis confirmed this result (**Fig 5b,** and **Table 2**). However, when *RBD_(331-529)_-Ec* was analyzed by SDS-PAGE under non-reducing conditions (**Fig. 5a**, Lane 3) a band with a very high molecular mass between the stacking and separating gel was observed. Also an smear in the stacking gel suggested the presence of *RBD_(331-529)_-Ec* aggregates mediated by multiple and probably random disulfide bonds. Probably this is the reason why, we did not succeed in measuring the intact molecular mass of this protein by ESI-MS analysis.

**Fig. 5.**
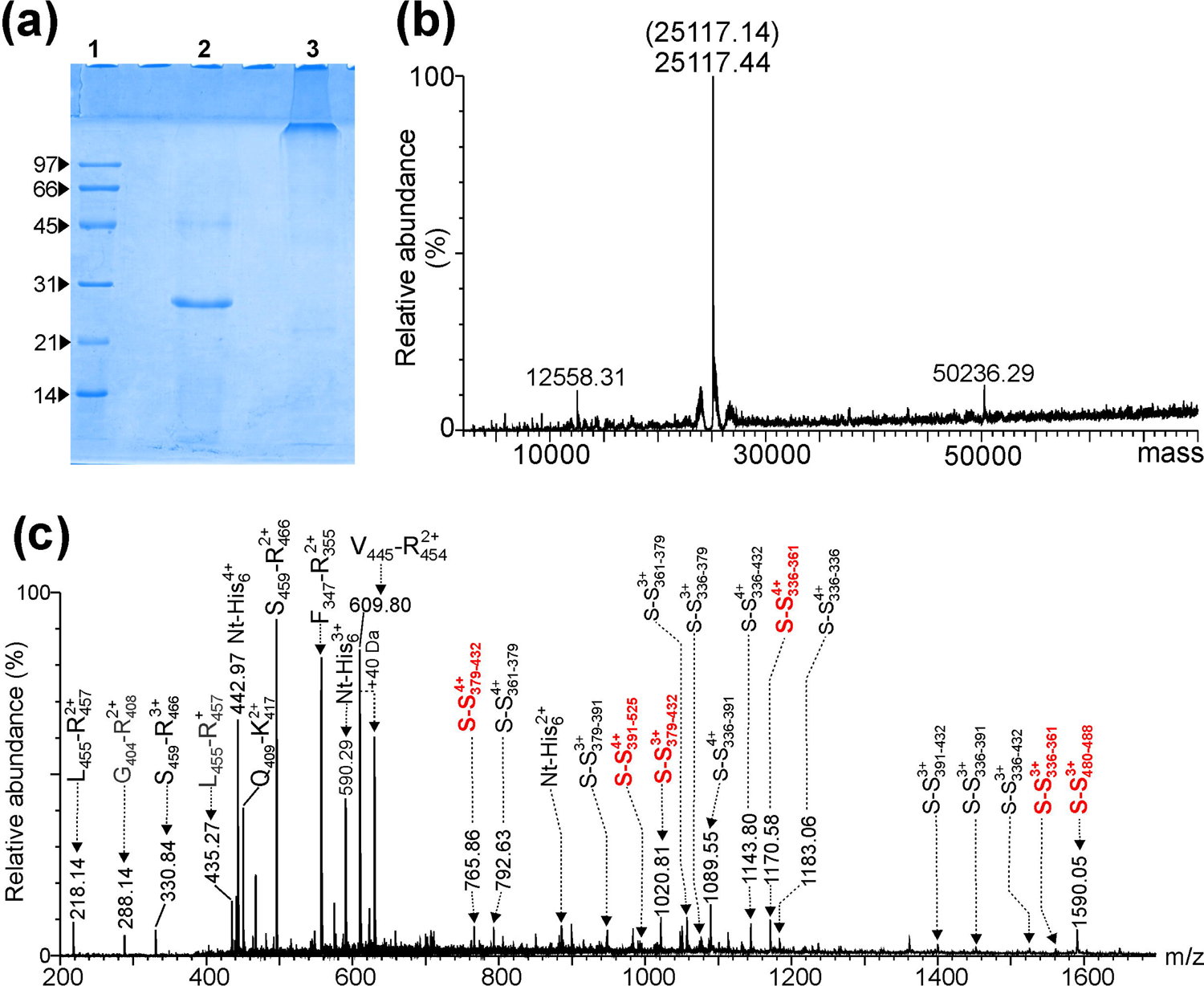
(a) SDS-PAGE analysis of the recombinant *RBD_(331-529)_-Ec* analyzed under reducing (Lane 2) and non-reducing (Lane 3) conditions and detected with Coomassie staining. Lane 1 corresponds to the molecular weight markers of low-range from 14 to 97 kDa (Bio-Rad). **(b)** Deconvoluted ESI-MS spectrum of the reduced and S-carbamidomethylated protein. The expected molecular mass is indicated between parentheses. **(c)** ESI-MS analysis of the recombinant expressed in *E. coli* and digested with trypsin by using in-solution BFD protocol. Signals assigned as (S-S)^n+^ correspond to the peptides containing disulfide bonds between the cysteines described. The signals labeled with (Nt-His_6_)^n+^ correspond to the N-terminal peptide containing the tandem repeat of six histidine residues in its amino acid sequence. A detailed assignment for all tryptic peptides in this figure is summarized in **Table S3**.

For this reason *RBD_(331-529)_-Ec* was considered as a positive control to evaluate if the in-solution BFD protocol (**Fig. 1b**) might allow the identification of the native and a wide variety of non-native disulfide bonds in the same ESI-MS spectrum.

ESI-MS analysis of *RBD_(331-529)_-Ec* digested with trypsin by using the in-solution BFD protocol showed several multiply-charged ions signals assigned to peptides containing Cys corresponding to the four native disulfide bonds (signals written in red and assigned as S-S_#-#_, **Fig. 5c**). The good agreement between the expected and experimental molecular masses of other signals written in black and assigned as S-S_#-#_, (**Fig. 5c**) were assigned to tryptic peptides containing scrambled disulfide bonds of *RBD_(331-529)_-Ec* (**Table S3**). The MS/MS spectra that confirmed these assignments are shown in **Fig. S11**. **Table S3** shows a summary for the assignment of all signals observed in the ESI-MS spectrum of **Fig. 5c**.

The charge state (z) of all these signals assigned to peptides linked by scrambled disulfide bonds was equal or higher than 3+ (**S-S**, in **Fig. 5c**). It agrees with a previous observation (49, 50) showing that in ESI-MS analysis cross-linked peptides generated by tryptic digestion are ionized preferably with z≥ 3+. Intermolecular disulfide linked peptides are a particular case of cross-linked peptides.

The results shown here demonstrated that in-solution BFD protocol (24) in combination with ESI-MS analysis of RBD enabled in a single mass spectrum the detection of native disulfide bonds, the scrambled variants, as well as free cysteine residues that might be responsible for promoting disulfide exchange and protein aggregation (22). This example is important for a future validation of this technique according to the ICHQ2R1 guidelines (51). These aspects fulfills the requirements of regulatory agencies for the development of a well-characterized product taking into account the importance of this PTM for the ICHQ6B guidelines (18). Ninety-nine percent of sequence coverage for *RBD_(331-529)_-Ec* was achieved when used the in-solution BFD protocol.

### Characterization of *RBD_(333-527)_-C1*

*Thermothelomyces heterothallica*, was engineered to develop an industrialized protein production host expression system with high yields (> 10 g/L) of several recombinant proteins including Mabs, Fc-fusion proteins, virus like particles, etc (14).

This engineered *T. heterothallica* C1 host expression system has a very significant reduction of the protease load thus minimizing unwanted degradation during fermentation (14). For these reasons, the *RBD_(333-527)_-C1* (see **Table 1**) was expressed in *T. heterothallica C1* host expression system and characterized using the in-solution BFD protocol. Unlike other proteins characterized in this work, *RBD_(333-527)_-C1* has only one N-glycosylation site located at Asn_343_. NP-HPLC profile showed the structural assignment based on the GU indexes for the individual N-glycans released with PNGase F and labeled with 2AB (**Fig. 6a**). ESI-MS spectrum deconvoluted with MaxEnt1 of the intact *RBD_(333-527)_-C1* confirmed the typical heterogeneity of N-glycoproteins indicating the presence of several non-fucosylated glycoforms (**Fig. 6b**). The experimental and expected molecular masses agreed very well (**Fig. 6b, Table 2**). In both analyses, the three predominant glycoforms were Man4, Man4A1 and Man5A1, being the last one the most-abundant.

**Fig. 6.**
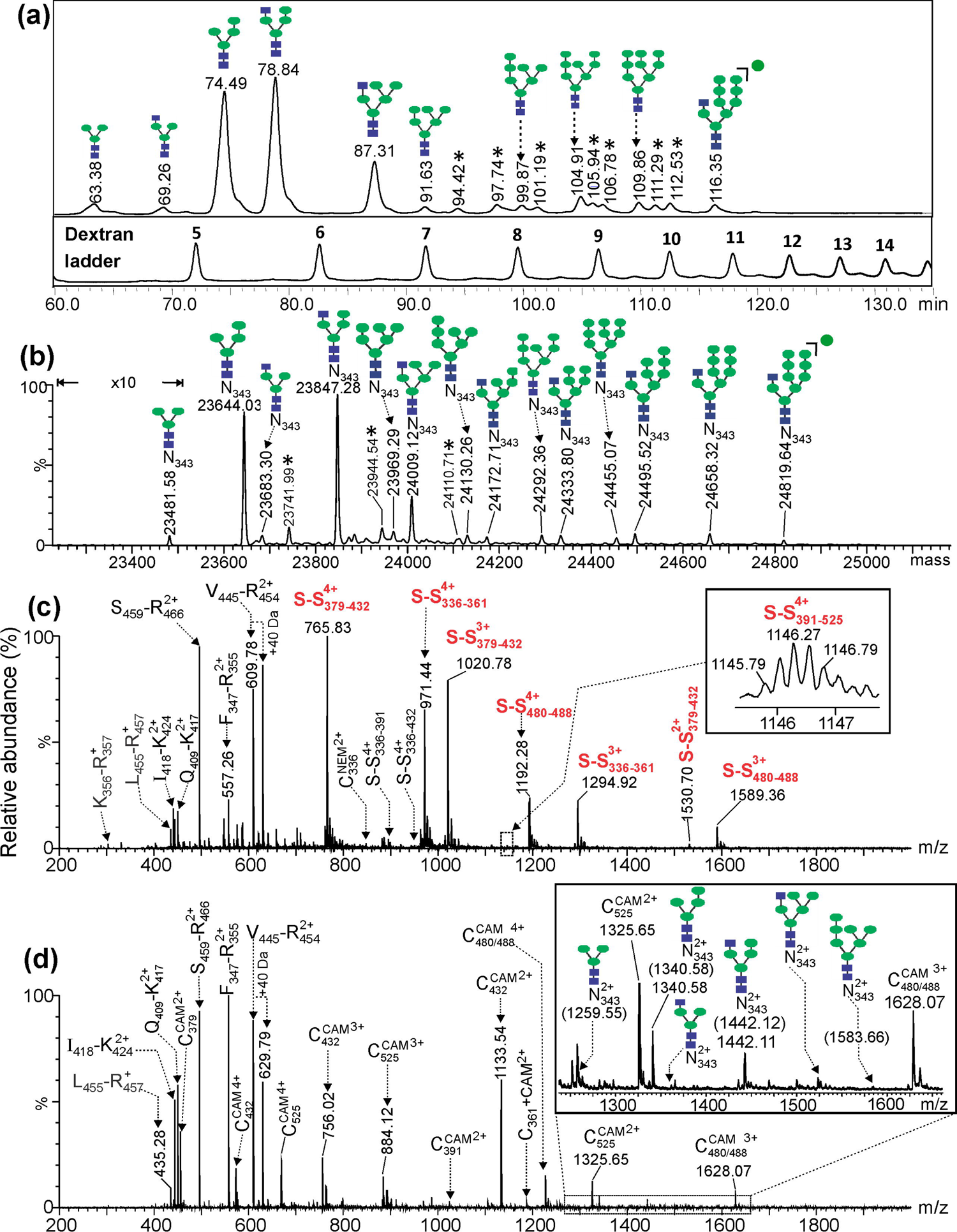
(a) NP-HPLC profile (upper chromatogram) of the 2AB-N-glycans released by PNGase F treatment of the recombinant *RBD_(333-527)_-C1* and corresponding dextran ladder (lower chromatogram) used to calculate the GU indexes for all 2AB-N-glycans and to perform for the structural assignment. The asterisks correspond to non-assigned glycoforms. The numbers above peaks in the dextran ladder indicate the corresponding glucose units. The nomenclature used in the structural assignment of the 2-AB *N*-glycans agrees with the proposed by the SNFG system (60). The deconvoluted ESI-MS spectrum shown in **(b)** corresponds to the intact with potential N-glycosylation site located at the Asn^343^ occupied to several glycoforms. A magnification of 10x is shown in the low molecular mass region of **(b)**. The ESI-MS spectrum shown in **(c)** correspond to the *RBD_(333-527)_-C1* treated with PNGase F and digested following the in-solution BFD protocol shown in **Fig. 1b**. The ESI-MS spectrum shown in **(d)** correspond to the reduced and S-alkylated glycosylated *RBD_(333-527)_-C1*. Signals assigned as (C#+cam)^n+^ correspond to tryptic peptides containing carbamidomethyl cysteine residues at position #. The inset shown in **(d)** corresponds to an expanded region (*m/z* 1237-1662) showing the presence of several signals assigned to the N-terminal end glycopeptides (T_333_-R_346_) with several N-glycans linked to the glycosylated Asn^343^. Signal assigned as (C_480/488_+cam)^3+^ correspond to the peptide D_467_-R_509_ containing the Cys_480_ and Cys_488_ S-alkylated with iodoacetamide. A detailed assignment for all tryptic peptides in this figure is summarized in **Table S4**.

This protein, after N-deglycosylation with PNGase F and ESI-MS analysis (**Fig. S8a**), also showed several multiply charged signals with z=8+ to 15+, that once deconvoluted with MaxEnt1 (**Fig. S8b**) yields an intense and unique signal with an experimental molecular mass of 22590.33 Da (**Table 2**). This result agrees very well with the expected (22590.26 Da) assuming the *RBD_(333-527)_-C1* monomer N-deglycosylated and with four disulfide bonds.

The N-deglycosylated protein digested with trypsin by in-solution BFD protocol (**Fig. 6c**) and analyzed by the ESI-MS allowed a full-sequence coverage (**Table 2**), confirmed the integrity of the N- and C-terminal ends and allowed the identification of the four native disulfide bonds (S-S_379-432_, S-S_336-361_, S-S_480-488_ and S-S_391-525_, **Table S4**). Very low-abundance signals (**Fig. 6c**) were detected at *m/z_Exp_* 847.87 (2+) and 1167.52 (2+) and assigned to the peptides T_333_-R_346_ and L_425_-K_444_ containing the Cys_336_ and Cys_432_ modified with NEM (**Fig. S9**). It indicates that a minor fraction of *RBD_(333-527)_-C1* contains Cys_336_ and Cys_432_ with free thiols in the original molecule. In addition, the same Cys_336_ and Cys_432_ were also detected in three low-intensity signals detected at *m/z_exp_* 889.72, 4+; *m/z_exp_* 944.43, 4+ and *m/z_exp_* 1131.26, 4+ (see **Table S4**) that were assigned to (T_333_-R_346_)-S-S-(L_387_-R_403_), (T_333_-R_346_)-S-S-(L_425_-K_444_) and (I_358_-K_378_)-S-S-(L_425_-K_444_) linked by the scrambled disulfide bonds between Cys_336_-Cys_391_, Cys_336_-Cys_432_ and Cys_361_-Cys_432_, respectively (**Fig. S9**). Scrambled Cys_361_-Cys_525_ was also detected and the MS/MS spectra supporting these assignments are shown in **Fig. S9**. The presence of free cysteine in the molecule probably is the responsible for the generation of these two low-abundance scrambling variants according to the mechanisms proposed for the generation of this modification (22).

The size heterogeneity of N-glycans linked to Asn_343_ in *RBD_(333-527)_-C1* was not revealed by the trypsin digestion and the ESI-MS analysis of **Fig. 6c** due to the treatment with PNGase F at the initial steps of the in-solution BFD protocol. A variant of the in-solution BFD protocol without the PNGase F treatment did not provided this information either because the N-terminal peptide (T_333_-R_346_) of the *RBD_(333-527)_-C1* containing the glycosylated Asn_343_ is linked at the same time to (I_358_-K_378_) by a disulfide bond (Cys_336_-Cys_361_). Probably, the heterogeneity typical of N-glycopeptides, combined with its high molecular mass (over 4 kDa) and its ionization suppressed at the same time by the presence of shorter proteolytic peptides make difficult it detection by ESI-MS analysis.

However, when the N-glycosylated *RBD_(333-527)_-C1* was reduced and S-alkylated with iodoacetamide and digested using the in-solution BFD all cysteine containing peptides were detected (**Fig. 6d**, **Table S4**) including the N-terminal peptide T_333_-R_346_ containing Cys_336_ and several glycoforms as shown in the inset of **Fig. 6d**. MS/MS spectra supporting these assignments are shown in **Fig. S10**.

The in-solution BFD protocol achieved 100% sequence coverage (**Table 2**) when the *RBD_(333-527)_-C1* was N-deglycosylated under non-reduced conditions (**Fig. 6c**) and also when the N-glycoprotein was reduced and carbamidomethylated (**Fig. 6d**).

### Characterization of *RBD_(331-530)_-Cmyc-Pp*

RBD of SARS-CoV-2 was also expressed in *P. pastoris* with a tandem repeat of six histidine residues and the Cmyc tag fused at the C-terminal end (*RBD_(331-530)_-Cmyc-Pp,* see **Table 1**) to be used for analytical purposes.

The ESI-MS spectrum of *RBD_(331-530)_-Cmyc-Pp* deglycosylated with PNGase F (**Fig. 7a**) with charge-states from 9+ to 17+ was deconvoluted (**Fig. 7b**), yielding an intense signal with a molecular mass of 25835.29 Da that is 400.88 Da higher than expected (25434.41 Da) considering the sequence of *RBD_(331-530)_-Cmyc-Pp* in **Table 1** with four disulfide bonds and two N-deglycosylated sites. The N-deglycosylated protein was digested with trypsin by the in-solution BFD protocol (**Fig. 1b**) and the resultant ESI-MS spectrum (**Fig. 7c**) showed an unexpected signal of appreciable intensity at *m/z_Exp_* 1219.32, 4+. The MS/MS spectrum of this signal (**Fig. 7d**) confirmed that two peptides [*EAEAEFS-*(D^331^-R^346^)-S-S-(I^358^-R^378^)] were linked by an intermolecular disulfide bond between Cys_336_ and Cys_361_. One of these peptides [*EAEAEFS*-(D_331_-R_346_)] contains an incomplete processed fragment of the alpha mating factor signal peptide (*EAEA-*) (52) linked to the expected N-terminal end EFS-(D_331_-R_346_) of the mature *RBD_(331-530)_-Cmyc-Pp*. The expected molecular mass of the residues (*EAEA-*) linked to the N-terminal end (400.39 Da) agrees with the mass difference observed between the experimental and calculated molecular mass for the N-deglycosylated protein (400.88 Da, **Fig 7b**).

**Fig. 7.**
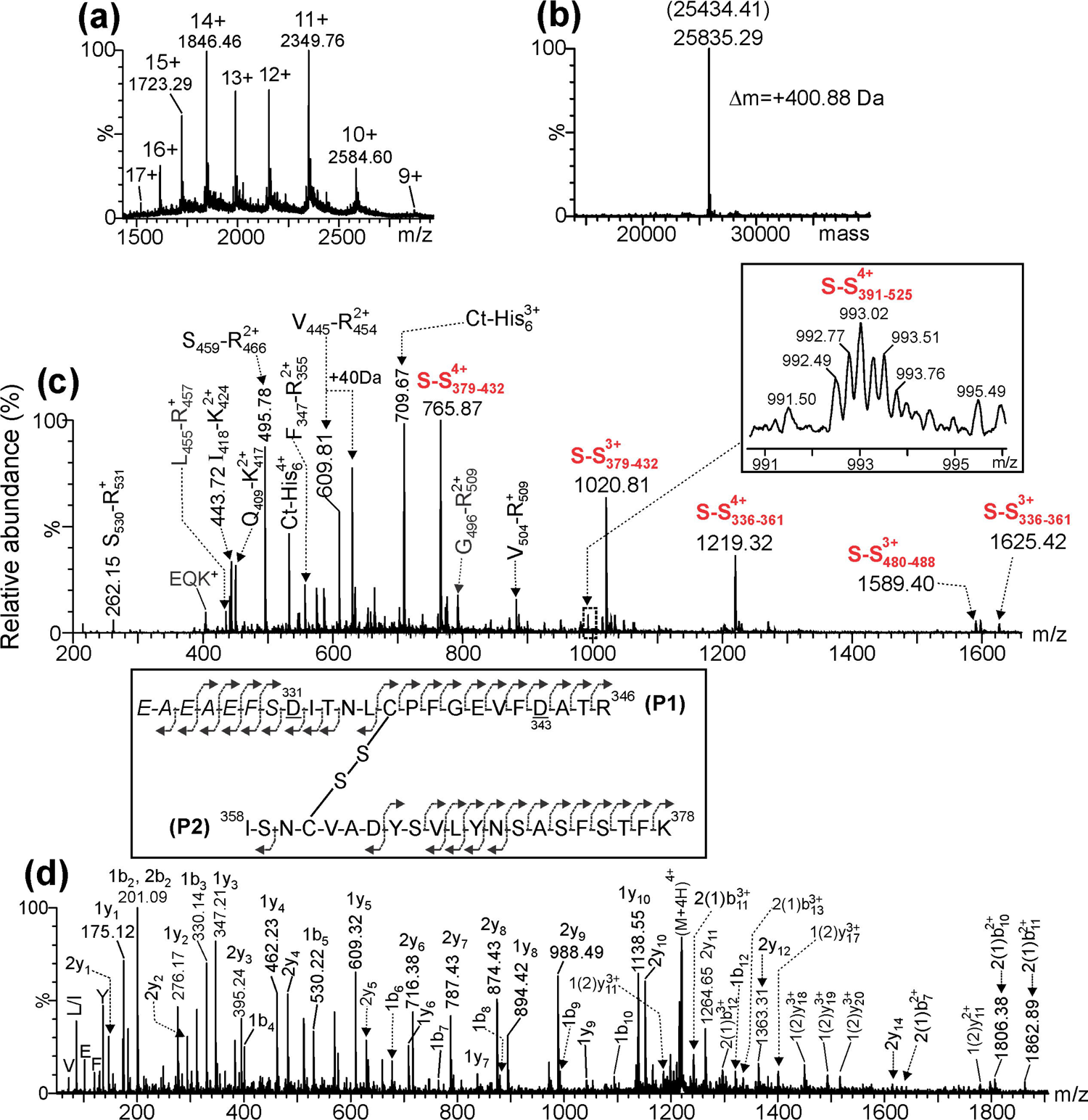
(a) ESI-MS analysis of the deglycosylated *RBD_(331-530)_-cmyc-Pp* expressed in *P. pastoris*. **(b)** Deconvoluted ESI-MS spectrum. Between parentheses the expected mass of the N-deglycosylated protein is indicated. **(c)** ESI-MS analysis of the *in-solution* BFD trypsin digestion of the N-deglycosylated *RBD_(331-530)_-cmyc-Pp.* The inset shows the isotopic ion distribution of a 4+ion corresponding to peptides [Leu_387_-Arg_403_]-*S*-*S*-[Val_510_-Lys_528_] linked by a disulfide bond between C_391_-C_525_. A summary of the above results is shown in **Table 2-3** and the detailed assignment for all signals in **(c)** is shown in **Table S5**. **(d)** ESI-MS/MS spectrum of peptides [*EAEAEFS*-Asn_331_-Arg_346_]-*S*-*S*-[Ile_358_-Lys_378_] linked by a disulfide bond between C_336_ and C_361_. This specie contains an extension of seven amino acids (*EAEAEFS*-) added to the expected N-terminal end [Asn_331_-Arg_346_] due to an incomplete processing of the signal peptide (alpha mating factor) during protein expression. Asn_331_ and Asn_343_ are transformed into Asp residues due to the action of PNGase F. The nomenclature for the fragment ions observed in the MS/MS spectrum agrees with the proposed by Mormann et al (61).

**Table S5** shows a summary for the assignment of all signals observed in the ESI-MS spectrum of **Fig. 7c**. Several short and hydrophilic peptides (^356^KR^357^; ^404^GDEVR^408^; ^455^LFR^457^; ^530^S*R*^531^; *EQK*) were detected and sequence coverage of 99% (**Table 2**) was achieved for the in-solution BFD protocol.

The α-mating factor prepro peptide secretion signal from *Saccharomyces cerevisiae* is still the most commonly used signal sequence for recombinant proteins expressed in *P. pastoris* (53). Processing of the alpha mating factor should occur in three steps, in particular the last step involves the Ste13 protein that cleaves the Glu-Ala repeats in the Golgi (54), therefore we speculate that this step was interrupted in some way, as all the purified protein was detected exclusively with the *EAEA-* linked to the N-terminal end. Probably the high expression level of this protein (40 g/L) impaired the complete processing of the signal peptide.

In-solution BFD enabled the detection of incomplete processing of the signal peptide of RBD expressed in *P. pastoris*, an expression system of choice for vaccine development due to the ease of genetic manipulation, the capability to perform complex post-translational modifications and, at the same time, obtain high expression yield (12). The characterization of the N-terminal end is also one of the aspects requested by the ICHQ6B guidelines (18).

### Artificial modifications introduced during sample processing by the in-solution BFD protocol

In the characterization of all RBDs by using in-solution BFD protocol we initially used acetone for protein precipitation (**Fig. 1b**). We noticed in the ESI-MS spectra an unexpected doubly-charged signal at *m/z_Exp_* 629.81 (**Fig 8a**) having a variable intensity. This signal was not detected when RBDs were processed by using in-solution SD protocol (**Fig. 2d, 4d and S6c**) and when the protein precipitation step (**Fig. 1b**) was carried out with cold ethanol (**Fig. 8b**) instead of acetone (**Fig. 8a**).

**Fig. 8.**
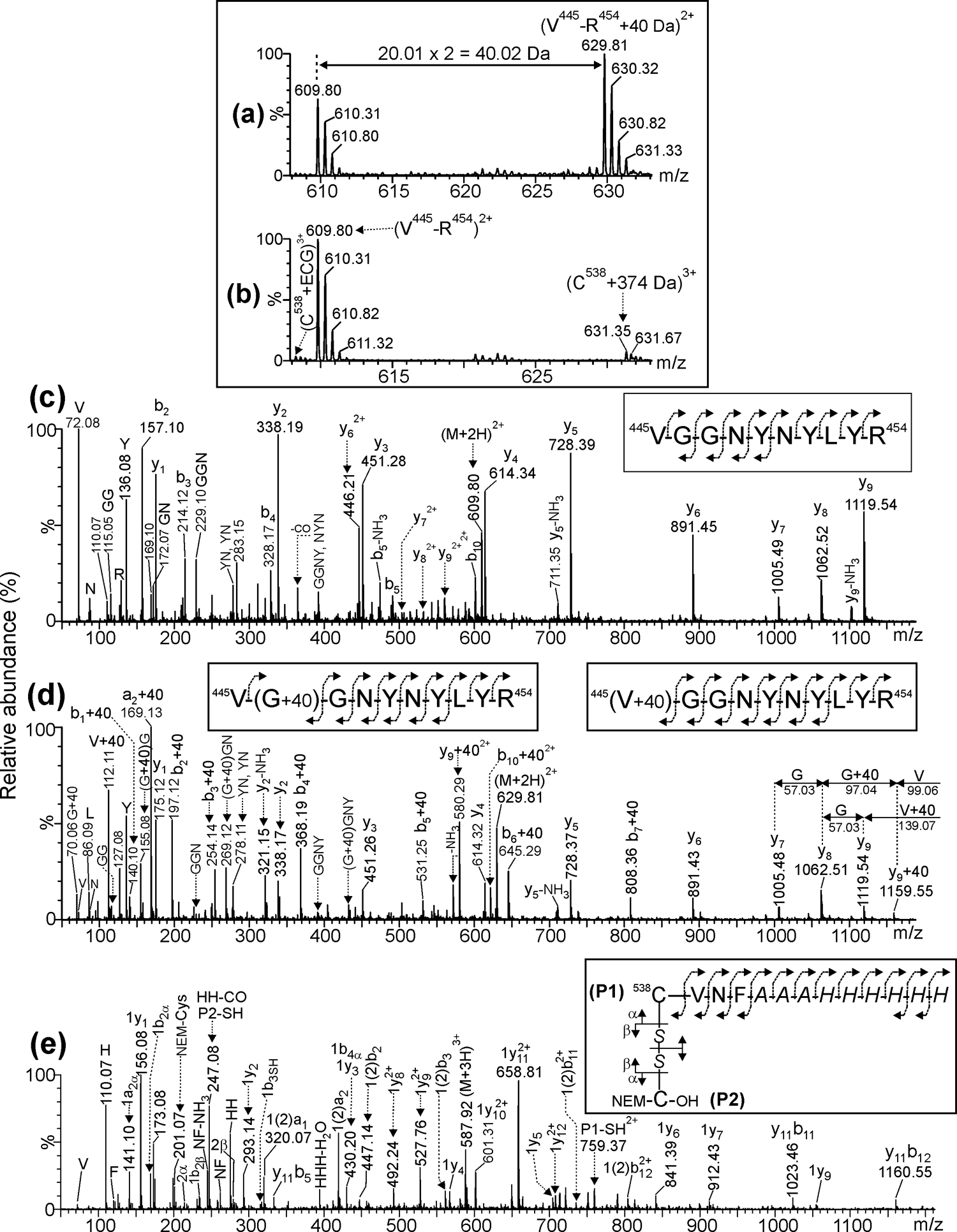
The ESI-MS spectra shown in **(a)** and **(b)** correspond to expanded regions of the tryptic peptides derived from *RBD_(319-541)_-HEK_A_3_* digested by in-solution BFD protocol after precipitation with acetone and ethanol, respectively. The signals assigned in **(b)** as (C^538^+ECG)^3+^ and (C^538^+374 Da)^3+^ correspond to the C-terminal peptide ^538^CVNF^541^-*AAAHHHHHH* with the C^538^ modified with glutathione and a chemical modification of unknown chemical nature that increased its molecular mass in 374 Da, respectively. The MS/MS spectra shown in **(c)** and **(d)** correspond to the internal non-modified Val^445^-Arg^454^ peptide (*m/z* 609.80, 2+) and the same peptide with a modification that increased its molecular mass by 40.02 Da (*m/z* 629.81, 2+), respectively. This chemical modification introduced in the precipitation step with acetone is located alternatively at the N-terminal end (V+40) or at the second position glycine (G+40). The MS/MS spectra shown in **(e)** correspond to the cysteinylated peptide C-terminal end peptide (^538^CVNF^541^-*AAAHHHHHH*) with the C^538^ linked by a disulfide bond (-*S*-*S*-) to a Cys residue (C-OH) modified at the N-terminal end with an N-ethylmaleimide group (NEM-) introduced during the sample processing. Peptide and C-OH have been assigned as P1 and P2, respectively. The nomenclature of fragment ions is in agreement with the proposed by Mormann *et al* (61).

Comparison between the MS/MS spectra of the unmodified peptide (^445^VGGNYNYLYR^454^, *m/z_Exp_* 609.80, 2+, **Fig. 8c**) and the signal detected at *m/z_Exp_* 629.81, 2+ (**Fig. 8d**) revealed that it corresponds to the same internal peptide (Val^445^-Arg^454^) modified by adding 40 Da alternatively at Gly^446^ (^445^V[G+40]GNYNYLYR^454^) and at the N-terminal end (^445^[V+40]GGNYNYLYR^454^).

The ions detected at *m/z_Exp_* 1119.54 (y_9_), *m/z_Exp_* 140.10 (b_1_+40) and *m/z_Exp_* 112.10 (immonium ion of [^445^Val+40]) exclusively observed in the MS/MS spectrum of the modified peptide (**Fig. 8d**) support the presence of the +40 Da at the N-terminal end.

Also, in the same MS/MS spectra (**Fig. 8d**), the ion detected at *m/z_Exp_* 1159.54 (y_9_+40 Da), and the immonium ion of valine (Val) suggest the coexistence of a second specie with the intact N-terminal ^455^Val. In addition, the presence of fragment ions of series b_n+40_ (b_2_+40 Da to b_7_+40 Da) and several internal ions, only detected in **Fig. 8d** at *m/z_Exp_* 70.06 (G+40), *m/z_Exp_* 155. 08 ([G+40]G), m/z*_Exp_* 269.12 ([G+40]GN), *m/z_Exp_* 432.19 ([G+40]GNY), suggest that the modification of 40 Da mass is also present at the second position (G+40) (**Fig. 8d**).

Although in literature a structure for this modification has not been proposed yet, a previous work noticed that it is specific only for those peptides having Gly at position n+2 that were derived from tryptic digests of proteins previously precipitated with acetone (55). All RBDs characterized here have only one internal tryptic peptide ^445^V**G\***GNYNYLYR^454^ with this characteristic.

The acetone traces that remains adhered in the pellet, during trypsin digestion at 37 °C for 16 hours, are the responsible for this modification (55). The intensity of this modified peptide can be reduced considerably if a 15 minutes vacuum drying step is inserted in the protocol after acetone protein precipitation. However, care should be taken because an extensive drying makes more difficult dissolving the protein pellet in water/acetonitrile as required for an efficient in-solution BFD protocol.

In-solution BFD of proteins precipitated with ethanol and acetone yield very similar results and they can be used indistinctively. However, during the analysis of *RBD_(319-541)_-HEK_A_3_* after acetone precipitation, the isotopic ion distributions of the modified ^445^Val-Arg^454^ + 40 Da peptide (*m/z_Exp_* 629.81, 2+, **Fig. 8a**) and the C-terminal peptide (^538^CVNF^541^-*AAAHHHHHH*) carrying a +374 Da modification at Cys^538^ (**Fig. 8b**) were partially overlapped thus it impaired its detection. This modification at Cys^538^ was only detected when the protein *RBD_(319-541)_-HEK_A_3_* was precipitated with ethanol and analyzed by in-solution BFD (**Fig. 8b**). Another artifact originated by the sample processing was the partial addition of NEM to the N-terminal end of the RBD proteins despite maleimide has 1000 fold selectivity for thiols over amine groups at neutral pH (56).

The addition of NEM was verified by ESI-MS analyses of the RBD deglycosylated with PNGase F (**Table 3**) and confirmed by the ESI-MS/MS analysis of the N-terminal tryptic glycopeptides (**Fig. S11**). Despite the abundant fragmentation of glycans in the MS/MS of **Fig. S11**, three b_n_ ions (b_1_, b_3_ and b_4_) containing the N-terminal end of the peptide R_319_-R_328_ increased its masses by in 125 Da due to the addition of NEM were detected. In addition, the cysteinylated *RBD_(319-541)_-HEK_A_3_* also partially added two molecules of NEM, one at the N-terminal end of Arg^319^ and a second one to the N-terminal end of Cys linked to Cys^538^ (**Fig. 8e**).

The ESI-MS/MS spectrum shown in **Fig. 8e** confirm this finding because it corresponds to the C-terminal peptide (^538^CVNF^541^-*AAAHHHHHH, m/z_Exp_* 587.92, 3+) of *RBD_(319-541)_-HEK_A_3_* linked by a disulfide bond to a cysteine residue that contains a modifying NEM group (+125 Da) at its N-terminal end. The symmetric dissociation of the disulfide bond yields the reduced ^538^CVNF^541^-*AAAHHHHHH (m/z_Exp_* 759.37, 2+ assigned as P1-SH^2+^) and the reduced NEM-C-OH *(m/z_Exp_* 247.08, 1+, assigned as P2-SH). Several daughter ions originated by the asymmetric fragmentation of the disulfide bond (1a_2α_, 1b_2α_, 1b_4α_, 1b_2β_, 1b_3SH_) confirmed that the N-terminal end of ^538^CVNF^541^-*AAAHHHHHH* is free while two other fragment ions (2_α_ and 2_β_) confirm that NEM is located at the N-terminal end of the cysteine residue that is linked by an intermolecular disulfide bond to Cys^538^.

Using the in-solution BFD protocol (**Fig. 1b**) the remaining internal tryptic peptides were not modified with NEM at their N-terminal ends because this S-alkylating reagent was eliminated during sample precipitation and the subsequent washing steps before proceeding to the proteolytic digestion. However, three low-intensity signals in the ESI-MS analysis of tryptic digestion corresponding to peptides ^356^KR^357^ (*m/z_Exp_* 428.26, 1+), ^458^KSNLKPFER^466^ (*m/z_Exp_* 622.36, 2+) and ^529^KSTNLVK^535^ (*m/z_Exp_* 457.78, 2+) with the epsilon amino group of Lys_356_, Lys_458_ and Lys_529_ modified with NEM (+125 Da) were detected using the in-solution BFD protocol and confirmed by MS/MS (**Fig. S12**). On the contrary, when the RBD is digested in-solution by using the SD protocol and NEM is present even at a very low concentration (≤5 mM) during all sample processing it will be added to the N-terminal end of most of the internal tryptic peptides (data not shown).

NEM is added in excess at a concentration of 5 mM and it remains during the N-deglycosylation step (2 h at 37 °C) at a pH slightly over neutral (7.2-7.4). It seems that these conditions make favorable this side reaction in a considerable fraction of the N-terminal end of the deglycosylated RBD as well as for the cysteine linked by disulfide bond to Cys^538^. In a minor extension the epsilon amino groups of Lys residues were modified. Therefore, the partial addition of NEM at the N-terminal end of the protein is a side reaction to be considered when in-solution BFD is used.

The hydrolysis of thiosuccinimide ring after adding the NEM to free Cys residues in protein was also observed probably more appreciable when the digestion of the analyzed RBD was carried out by the SD protocol at basic pH (**Fig. 1a**) (57). This side reaction increased the molecular masses of peptides by 18 Da.

## Conclusions

In-solution BFD allowed in a single ES-MS spectrum, the full-sequence coverage for most recombinant RBD sequences characterized in this work and outperformed, in this aspect, the in-solution SD protocol. Hydrophobic and the hydrophilic peptides were simultaneously and efficiently detected in the same ESI-MS spectrum only when BFD protocol was used.

The in-solution BFD protocol implemented in combination with ESI-MS analysis has demonstrated to be sensitive for the detection of PTMs (N- and O-glycosylation, and several modifications linked to the free Cys^538^) present in the recombinant RBDs produced in different expression systems (mammalian cells, fungus, yeast and bacteria). In an RBD with the unpaired Cys^538^ several PTMs were only detected when in-solution BFD was applied.

In particular, the identification of the C-terminal end of proteins containing a tandem repeat of six histidine residues, an important aspect requested in the ICHQ6B guidelines, was always possible by the in-solution BFD while with the SD sample processing the identification was not achieved in all proteins.

Artifacts introduced during the application of the in-solution BFD protocol like the one provoked by acetone (+40 Da) and the N-terminal modification with NEM (+125 Da) were well characterized and the sources that originate them were readily identified. In particular, the artifact of +40 Da in an internal peptide of RBD (^445^VG*GNYNYLYR^454^) can be eliminated if ethanol precipitation is used instead of acetone. Other S-alkylating agents might be also used alternatively depending the preferences of the labs, but was not explored here.

We believe that the results obtained in this study, suggest that in-solution BFD protocol in combination with ESI-MS deserve to be validated for the characterization of RBDs used as active pharmaceutical ingredients of SARS-CoV-2 subunit-based vaccines. This protocol is simple and can be easily implemented because it essentially employs reagents frequently used for the characterization of proteins by MS.

MALDI-MS in principle, can also be used in combination with the in-solution BFD, however the detection of some low molecular mass peptides in somehow might be troublesome due to matrix interference. Probably the usage of blank mass spectra might be useful to obtain reliable results.

We foresee that the current BFD protocol can also be applicable to the analysis of active pharmaceutical ingredients of other recombinant RBD-based subunit vaccines customized for SARS-CoV-2 with point mutations (58) or other mutants generated by virus recombination phenomena (59). In particular, it could be of relevance in a context where vaccination constitutes a tremendous selective pressure for the onset of virus mutants and probably new subunit-based vaccine candidates might be required as it is the case of seasonable vaccines required for immunization against influenza.

## Availability of data and material

available if requested

## References

1. Wang C, Horby PW, Hayden FG, Gao GF. A novel coronavirus outbreak of global health concern. The lancet. 2020;395(10223):470–3.

2. WHO. WHO Coronavirus (COVID-19) Dashboard https://covid19.who.int [

3. Wouters OJ, Shadlen KC, Salcher-Konrad M, Pollard AJ, Larson HJ, Teerawattananon Y, et al. Challenges in ensuring global access to COVID-19 vaccines: production, affordability, allocation, and deployment. The Lancet. 2021.

4. WHO. Target product profiles for COVID-19 vaccines https://www.who.int/publications/m/item/whotarget-product-profiles-for-covid-19-vaccines [updated version 3. April 29, 2020.

5. Encinosa Guzmán PE, Bello Soto Y, Rodríguez-Mallon A. Genetic and biological characterization of a Cuban tick strain from Rhipicephalus sanguineus complex and its sensitivity to different chemical acaricides. International Journal of Acarology. 2016;42(1):18–25.

6. Zhou P, Yang X-L, Wang X-G, Hu B, Zhang L, Zhang W, et al. A pneumonia outbreak associated with a new coronavirus of probable bat origin. nature. 2020;579(7798):270-3.

7. Wrapp D, Wang N, Corbett KS, Goldsmith JA, Hsieh C-L, Abiona O, et al. Cryo-EM structure of the 2019-nCoV spike in the prefusion conformation. Science. 2020;367(6483):1260-3.

8. Yang J, Wang W, Chen Z, Lu S, Yang F, Bi Z, et al. A vaccine targeting the RBD of the S protein of SARS-CoV-2 induces protective immunity. Nature. 2020;586(7830):572-7.

9. Valdes-Balbin Y, Santana-Mederos D, Paquet F, Fernandez S, Climent Y, Chiodo F, et al. Molecular Aspects Concerning the Use of the SARS-CoV-2 Receptor Binding Domain as a Target for Preventive Vaccines. ACS Central Science. 2021.

10. Shang J, Ye G, Shi K, Wan Y, Luo C, Aihara H, et al. Structural basis of receptor recognition by SARS-CoV-2. Nature. 2020;581(7807):221-4.

11. Li T, Zheng Q, Yu H, Wu D, Xue W, Xiong H, et al. SARS-CoV-2 spike produced in insect cells elicits high neutralization titres in non-human primates. Emerging microbes & infections. 2020;9(1):2076–90.

12. Arbeitman CR, Auge G, Blaustein M, Bredeston L, Corapi ES, Craig PO, et al. Structural and functional comparison of SARS-CoV-2-spike receptor binding domain produced in Pichia pastoris and mammalian cells. Scientific reports. 2020;10(1).

13. Pollet J, Chen W-H, Versteeg L, Keegan B, Zhan B, Wei J, et al. SARS-CoV-2 RBD219-N1C1: A yeast-expressed SARS-CoV-2 recombinant receptor-binding domain candidate vaccine stimulates virus neutralizing antibodies and T-cell immunity in mice. Human Vaccines & Immunotherapeutics. 2021:1–11.

14. Visser H, Joosten V, Punt PJ, Gusakov AV, Olson PT, Joosten R, et al. Development of a mature fungal technology and production platform for industrial enzymes based on a Myceliophthora thermophila isolate, previously known as Chrysosporium lucknowense C1. Industrial Biotechnology. 2011;7(3):214–23.

15. Müller F, Fischer L, Chen ZA, Auchynnikava T, Rappsilber J. On the reproducibility of label-free quantitative cross-linking/mass spectrometry. Journal of The American Society for Mass Spectrometry. 2017;29(2):405–12.

16. Fujita R, Hino M, Ebihara T, Nagasato T, Masuda A, Lee JM, et al. Efficient production of recombinant SARS-CoV-2 spike protein using the baculovirus-silkworm system. Biochemical and Biophysical Research Communications. 2020;529(2):257–62.

17. Prahlad J, Struble L, Lutz WE, Wallin SA, Khurana S, Schnaubelt A, et al. Bacterial expression and purification of functional recombinant SARS-CoV-2 spike receptor binding domain. bioRxiv. 2021.

18. Rudge SR, Nims RW. ICH Q6B Specifications: Test Procedures and Acceptance Criteria for Biotechnological/Biological Products. ICH Quality Guidelines: An Implementation Guide. 2017:467.

19. Li Y, Lai D-y, Zhang H-n, Jiang H-w, Tian X, Ma M-l, et al. Linear epitopes of SARS-CoV-2 spike protein elicit neutralizing antibodies in COVID-19 patients. Cellular & Molecular Immunology. 2020;17(10):1095–7.

20. Poh CM, Carissimo G, Wang B, Amrun SN, Lee CY-P, Chee RS-L, et al. Two linear epitopes on the SARS-CoV-2 spike protein that elicit neutralising antibodies in COVID-19 patients. Nature Communications. 2020;11(1):2806.

21. Sanda M, Morrison L, Goldman R. N-and O-glycosylation of the SARS-CoV-2 spike protein. Analytical chemistry. 2021;93(4):2003–9.

22. Lakbub JC, Shipman JT, Desaire H. Recent mass spectrometry-based techniques and considerations for disulfide bond characterization in proteins. Analytical and bioanalytical chemistry. 2018;410(10):2467–84.

23. Castellanos-Serra L, Ramos Y, Huerta V. An in-gel digestion procedure that facilitates the identification of highly hydrophobic proteins by electrospray ionization-mass spectrometry analysis. Proteomics. 2005;5(11):2729–38.

24. Betancourt LH, Espinosa LA, Ramos Y, Bequet-Romero M, Rodríguez EN, Sánchez A, et al. Targeting the hydrophilic regions of recombinant proteins by MS via in-solution buffer-free trypsin digestion. European Journal of Mass Spectrometry. 2020;26(3):230–7.

25. Studier FW. Protein production by auto-induction in high density shaking cultures. Protein Expr Purif. 2005;41(1):207–34.

26. Laemmli UK. Cleavage of structural proteins during the assembly of the head of bacteriophage T4. Nature. 1970;227(5259):680-5.

27. Heukeshoven J, Dernick R. Characterization of a solvent system for separation of water-insoluble poliovirus proteins by reversed-phase high-performance liquid chromatography. Journal of Chromatography A. 1985;326:91–101.

28. Guile GR, Rudd PM, Wing DR, Prime SB, Dwek RA. A rapid high-resolution high-performance liquid chromatographic method for separating glycan mixtures and analyzing oligosaccharide profiles. Analytical biochemistry. 1996;240(2):210–26.

29. Kerr J, Schlosser JL, Griffin DR, Wong DY, Kasko AM. Steric effects in peptide and protein exchange with activated disulfides. Biomacromolecules. 2013;14(8):2822–9.

30. Monahan FJ, German JB, Kinsella JE. Effect of pH and temperature on protein unfolding and thiol/disulfide interchange reactions during heat-induced gelation of whey proteins. Journal of Agricultural and Food Chemistry. 1995;43(1):46–52.

31. Zhong X, He T, Prashad AS, Wang W, Cohen J, Ferguson D, et al. Mechanistic understanding of the cysteine capping modifications of antibodies enables selective chemical engineering in live mammalian cells. Journal of biotechnology. 2017;248:48–58.

32. Gstöttner C, Zhang T, Resemann A, Ruben S, Pengelley S, Suckau D, et al. Structural and functional characterization of SARS-CoV-2 RBD domains produced in mammalian cells. bioRxiv. 2021.

33. Crowell AM, Wall MJ, Doucette AA. Maximizing recovery of water-soluble proteins through acetone precipitation. Analytica chimica acta. 2013;796:48–54.

34. Ma J, Stoter G, Verweij J, Schellens JH. Comparison of ethanol plasma-protein precipitation with plasma ultrafiltration and trichloroacetic acid protein precipitation for the measurement of unbound platinum concentrations. Cancer chemotherapy and pharmacology. 1996;38(4):391–4.

35. Valdes-Balbin Y, Santana-Mederos D, Quintero L, Fernandez S, Rodriguez L, Sanchez-Ramirez B, et al. SARS-CoV-2 RBD-Tetanus toxoid conjugate vaccine induces a strong neutralizing immunity in preclinical studies. bioRxiv. 2021.

36. Shajahan A, Supekar NT, Gleinich AS, Azadi P. Deducing the N- and O-glycosylation profile of the spike protein of novel coronavirus SARS-CoV-2. Glycobiology. 2020;30(12):981–8.

37. Mechref Y. Use of CID/ETD mass spectrometry to analyze glycopeptides. Curr Protoc Protein Sci. 2012;Chapter 12:Unit-12.1.1.

38. Gadgil HS, Bondarenko PV, Pipes GD, Dillon TM, Banks D, Abel J, et al. Identification of cysteinylation of a free cysteine in the Fab region of a recombinant monoclonal IgG1 antibody using Lys-C limited proteolysis coupled with LC/MS analysis. Analytical biochemistry. 2006;355(2):165–74.

39. Buchanan A, Clementel V, Woods R, Harn N, Bowen MA, Mo W, et al., editors. Engineering a therapeutic IgG molecule to address cysteinylation, aggregation and enhance thermal stability and expression. MAbs; 2013: Taylor & Francis.

40. Banks DD, Gadgil HS, Pipes GD, Bondarenko PV, Hobbs V, Scavezze JL, et al. Removal of cysteinylation from an unpaired sulfhydryl in the variable region of a recombinant monoclonal IgG1 antibody improves homogeneity, stability, and biological activity. Journal of pharmaceutical sciences. 2008;97(2):775–90.

41. Bayer M, König S. Abundant cysteine side reactions in traditional buffers interfere with the analysis of posttranslational modifications and protein quantification–how to compromise. Rapid Communications in Mass Spectrometry. 2016;30(15):1823–8.

42. Kim HJ, Ha S, Lee HY, Lee KJ. ROSics: chemistry and proteomics of cysteine modifications in redox biology. Mass spectrometry reviews. 2015;34(2):184–208.

43. Moya G, Gonzalez LJ, Huerta V, Garcıa Y, Morera V, Perez D, et al. Isolation and characterization of modified species of a mutated (Cys125–Ala) recombinant human interleukin-2. Journal of Chromatography A. 2002;971(1-2):129–42.

44. Junutula JR, Bhakta S, Raab H, Ervin KE, Eigenbrot C, Vandlen R, et al. Rapid identification of reactive cysteine residues for site-specific labeling of antibody-Fabs. Journal of immunological methods. 2008;332(1-2):41–52.

45. Stimmel JB, Merrill BM, Kuyper LF, Moxham CP, Hutchins JT, Fling ME, et al. Site-specific conjugation on serine→ cysteine variant monoclonal antibodies. Journal of Biological Chemistry. 2000;275(39):30445–50.

46. Chen X, Nguyen M, Jacobson F, Ouyang J, editors. Charge-based analysis of antibodies with engineered cysteines: from multiple peaks to a single main peak. MAbs; 2009: Taylor & Francis.

47. Tang HY, Speicher DW. Experimental Assignment of Disulfide-Bonds in Purified Proteins. Current protocols in protein science. 2019;96(1):e86.

48. Dai L, Zheng T, Xu K, Han Y, Xu L, Huang E, et al. A universal design of betacoronavirus vaccines against COVID-19, MERS, and SARS. Cell. 2020;182(3):722–33. e11.

49. Giese SH, Fischer L, Rappsilber J. A study into the collision-induced dissociation (CID) behavior of cross-linked peptides. Molecular & Cellular Proteomics. 2016;15(3):1094–104.

50. Chen ZA, Jawhari A, Fischer L, Buchen C, Tahir S, Kamenski T, et al. Architecture of the RNA polymerase II-TFIIF complex revealed by cross-linking and mass spectrometry. The EMBO journal. 2010;29(4):717–26.

51. Borman P, Elder D. Q2 (R1) validation of analytical procedures. ICH Quality guidelines. 2017:127–66.

52. Kurjan J, Herskowitz I. Structure of a yeast pheromone gene (MFα): a putative α-factor precursor contains four tandem copies of mature α-factor. Cell. 1982;30(3):933–43.

53. Lin-Cereghino GP, Stark CM, Kim D, Chang J, Shaheen N, Poerwanto H, et al. The effect of α-mating factor secretion signal mutations on recombinant protein expression in Pichia pastoris. Gene. 2013;519(2):311–7.

54. Brake AJ, Merryweather JP, Coit DG, Heberlein UA, Masiarz FR, Mullenbach GT, et al. Alpha-factor-directed synthesis and secretion of mature foreign proteins in Saccharomyces cerevisiae. Proceedings of the National Academy of Sciences. 1984;81(15):4642–6.

55. Simpson DM, Beynon RJ. Acetone precipitation of proteins and the modification of peptides. Journal of proteome research. 2010;9(1):444–50.

56. Kratz H, Haeckel A, Michel R, Schönzart L, Hanisch U, Hamm B, et al. Straightforward thiol-mediated protein labelling with DTPA: Synthesis of a highly active 111 In-annexin A5-DTPA tracer. EJNMMI research. 2012;2(1):17.

57. Boyatzis AE, Bringans SD, Piggott MJ, Duong MN, Lipscombe RJ, Arthur PG. Limiting the Hydrolysis and Oxidation of Maleimide-Peptide Adducts Improves Detection of Protein Thiol Oxidation. Journal of Proteome Research. 2017;16(5):2004–15.

58. Cele S, Gazy I, Jackson L, Hwa S-H, Tegally H, Lustig G, et al. Escape of SARS-CoV-2 501Y.V2 from neutralization by convalescent plasma. Nature. 2021.

59. Zhu Z, Meng K, Meng G. Genomic recombination events may reveal the evolution of coronavirus and the origin of SARS-CoV-2. Scientific reports. 2020;10(1):1–10.

60. Varki A, Cummings RD, Aebi M, Packer NH, Seeberger PH, Esko JD, et al. Symbol nomenclature for graphical representations of glycans. Glycobiology. 2015;25(12):1323–4.

61. Mormann M, Eble J, Schwöppe C, Mesters RM, Berdel WE, Peter-Katalinić J, et al. Fragmentation of intra-peptide and inter-peptide disulfide bonds of proteolytic peptides by nanoESI collision-induced dissociation. Analytical and bioanalytical chemistry. 2008;392(5):831–8.

