## Supplemental files for "In-solution buffer-free digestion for the analysis of SARS-CoV-2 RBD proteins allows a full sequence coverage and detection of post-translational modifications in a single ESI-MS spectrum"

**SUPPORTING INFORMATION**

**Title:** In-solution buffer-free digestion permits high sequence coverage and the detection of post-translational modifications in ESI-MS analysis of several recombinant receptor-binding domains of SARS-CoV-2.

**Authors:** Luis Ariel Espinosa^1^, Yassel Ramos^1^, Ivan Andújar^1^, Enso Onill Torres^1^, Gleysin Cabrera^1^, Alejandro Martín^1^, Diamilé Roche^1^, Glay Chinea^1^, Mónica Becquet^1^, Isabel González^1^, Camila Canaán-Haden^1^, Elías Nelson^1^, Gertrudis Rojas^2^, Beatriz Pérez-Massón^2^, Dayana Pérez-Martínez^2^, Tamy Boggiano^2^, Julio Palacio^2^, Sum Lai Lozada Chang^2^, Lourdes Hernández^2^, Kathya Rashida de la Luz Hernández^2^, Saloheimo Markku^3^, Vitikainen Marika^3^, Yury **Valdés-Balbín^4^**, Darielys **Santana-Medero**^4^, Daniel G. Rivera^5^, Vicente Vérez-Bencomo^4^, Mark Emalfarb^3^, Ronen Tchelet^3^, Gerardo Guillén^1^, Miladys Limonta^1^, Eulogio Pimentel^1^, Marta Ayala^1^, Vladimir Besada^1^, Luis Javier González^1^*.

**Affilliation**:

^1^ Center for Genetic Engineering and Biotechnology (CIGB), Ave 31, e/ 158 y 190, Cubanacán, Playa, Havana, Cuba.

^2^ Center of Molecular Immunology, P.O. Box 16040, 216 St. Havana, Cuba.

^3^ VTT Technical Research Centre of Finland Ltd. Dyadic International, Inc.

^4^ Finlay Vaccine Institute, 200 and 21 Street, Havana 11600, Cuba.

^5^ Laboratory of Synthetic and Biomolecular Chemistry, Faculty of Chemistry, University of

Havana, Cuba.

***Corresponding author**:

Luis Javier González, PhD

Mass Spectrometry Laboratory,

Department of Proteomics,

Center for Genetic Engineering and Biotechnology

### **Fig. S1.** ESI-MS/MS spectra of the signals at *m/z*_Exp_ 1452.39, 3+ (a); 1020.83, 3+ (b); 1589.42, 3+ (c) and 992.54, 4+ (d) corresponding to the native disulfide bonds S-S_336-361_, S-S_379-432_, S-S_480-488_ and S-S_391-525_, respectively detected in *RBD_(319-541)_-HEK_A_3_*.

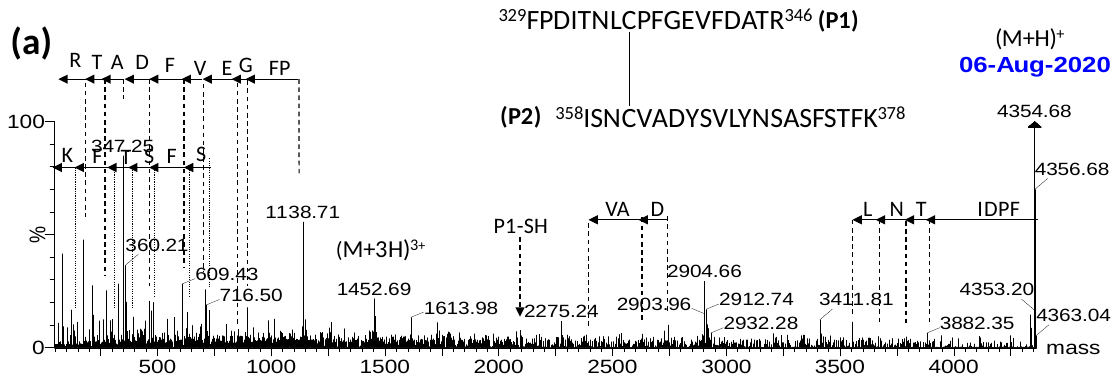

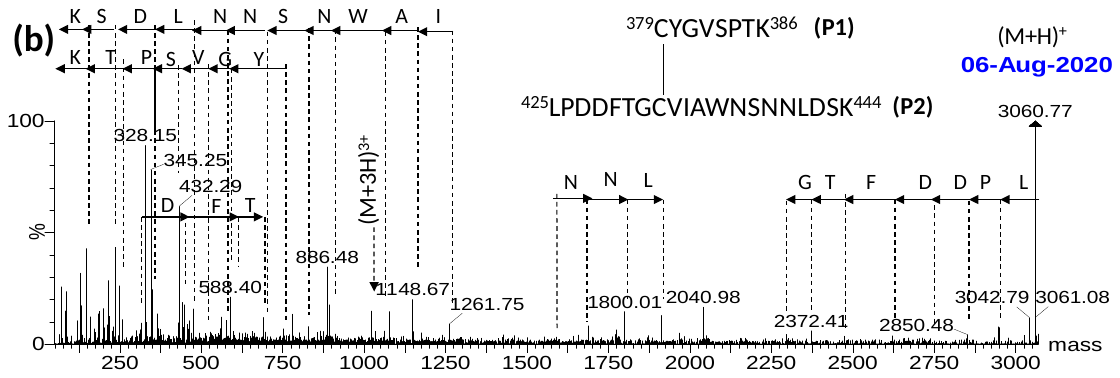

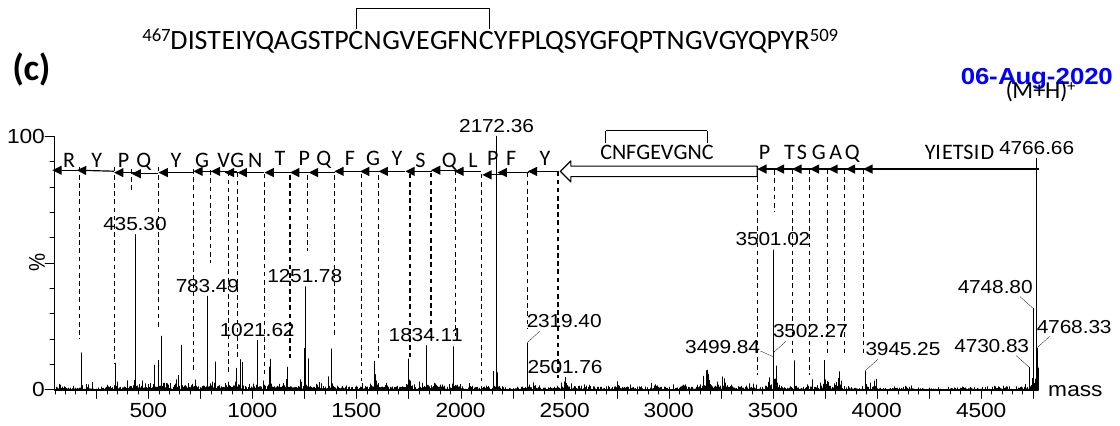

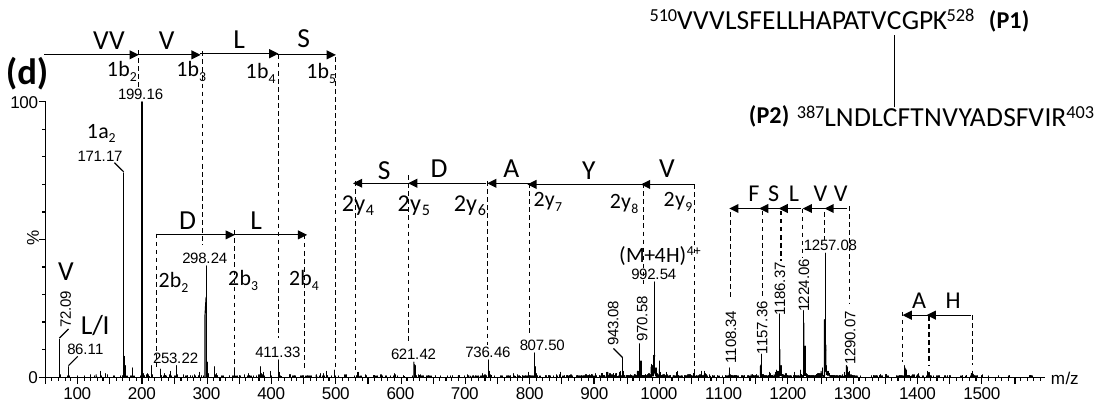

### **Fig. S2.** ESI-MS/MS spectra of the V^320^-R^328^ peptide containing the O-glycans HexNAc:Hex:NeuAc_2_, HexNAc:Hex:NeuAc and HexNAc-Hex detected in *RBD_(319-541)_-HEK_A_3_*.

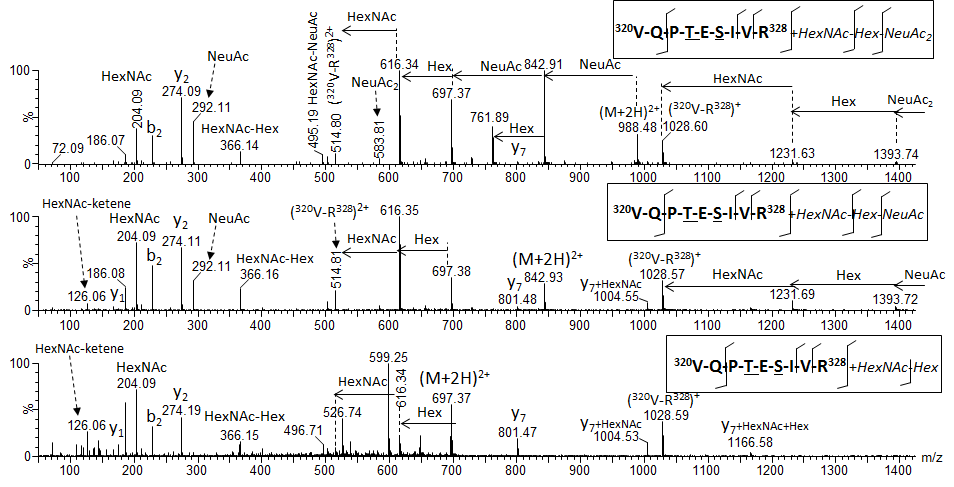

**Fig. S3.** ESI-MS/MS spectra of peptide ^538^CVNF^541^-*AAAHHHHHH* containing several known and unknown modifications at Cys_538_ detected in *RBD_(319-541)_-HEK_A3*. The nomenclature of fragment ions in peptides with disulfide bonds is in agreement with the proposed by Mormann et al (1).

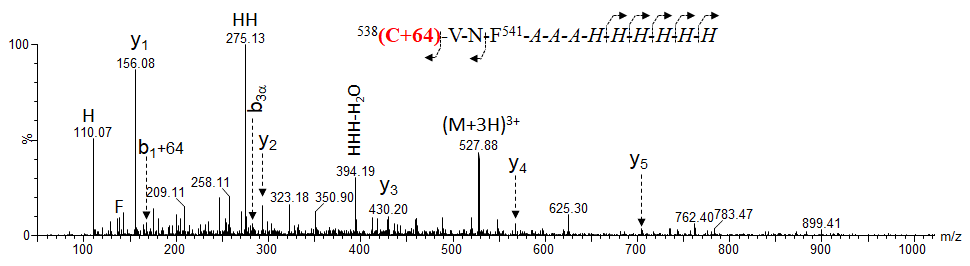

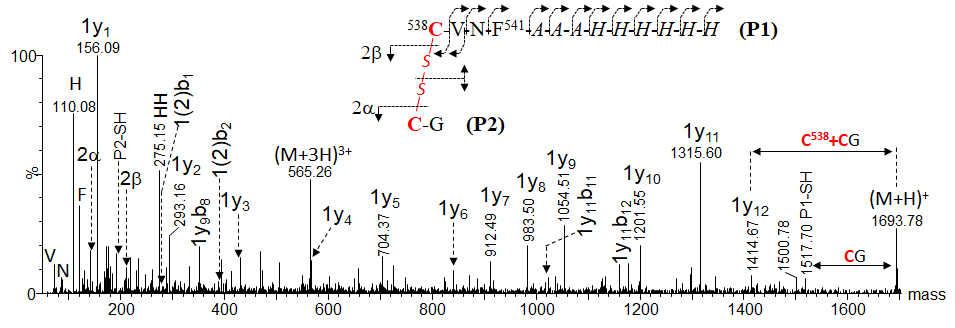

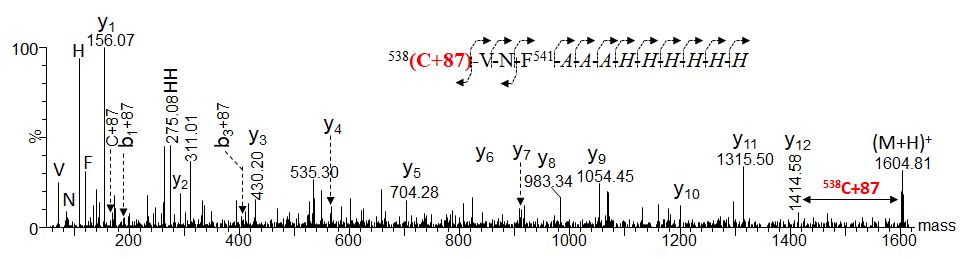

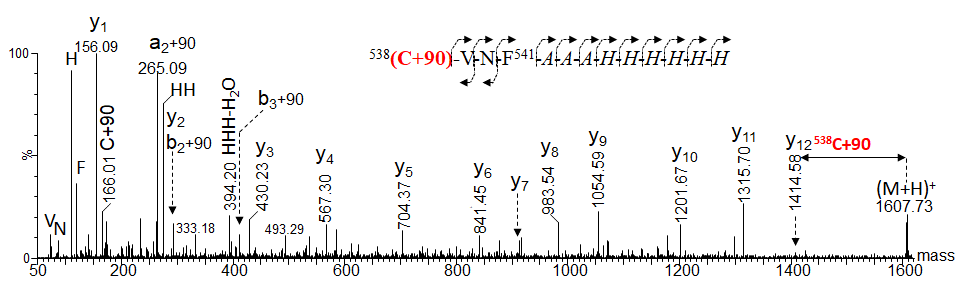

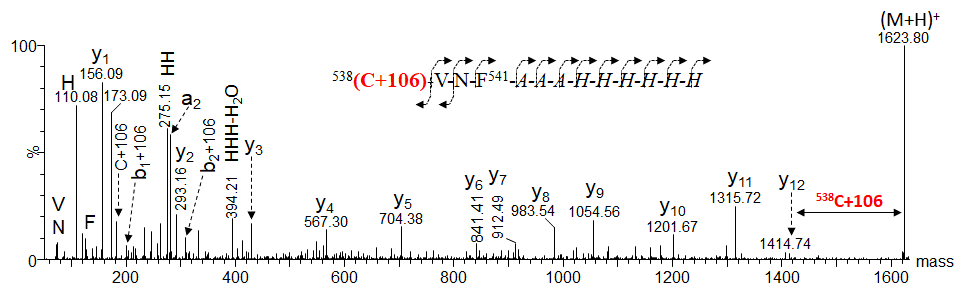

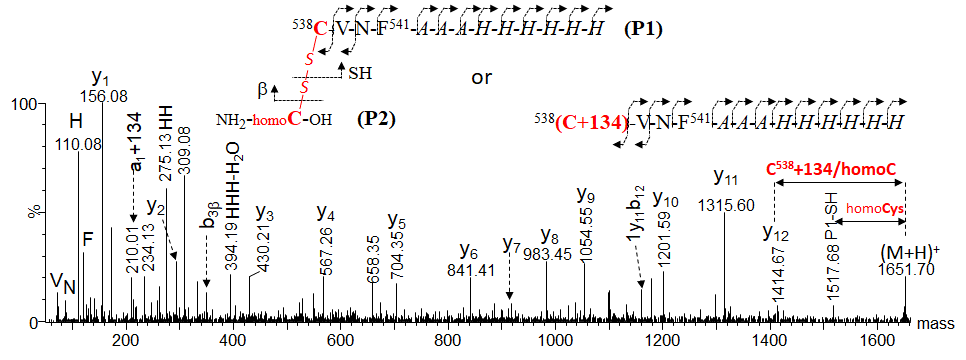

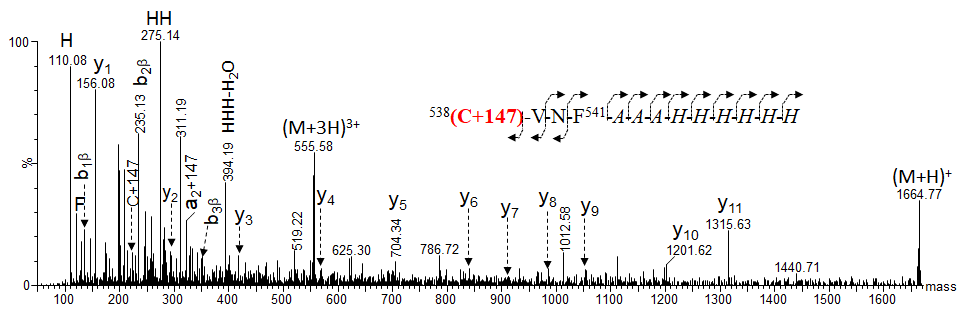

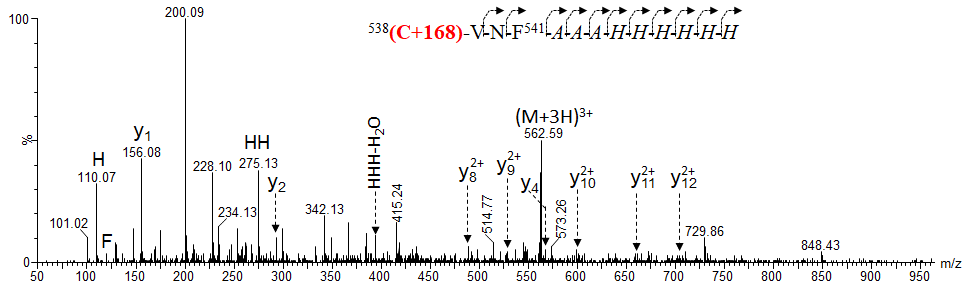

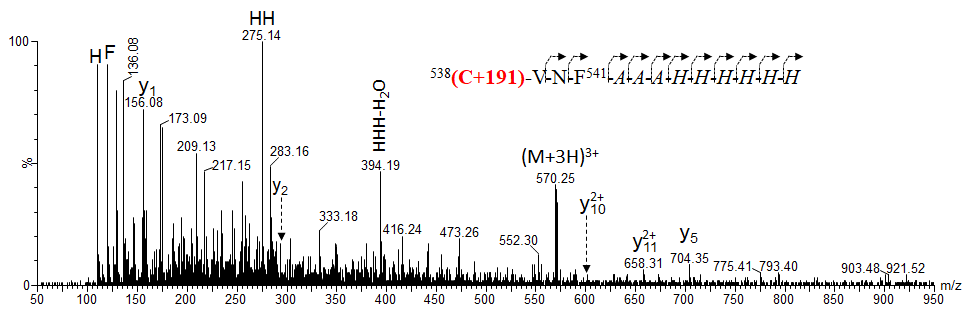

**Fig. S4.** ESI-MS/MS spectra of the low-abundance scrambled species of C_538_-C_379_ and C_538_-C_432_ detected in *RBD_(319-541)_-HEK_A3*. The nomenclature of fragment ions for disulfide bonds is in agreement with the proposed by Mormann et al (1).

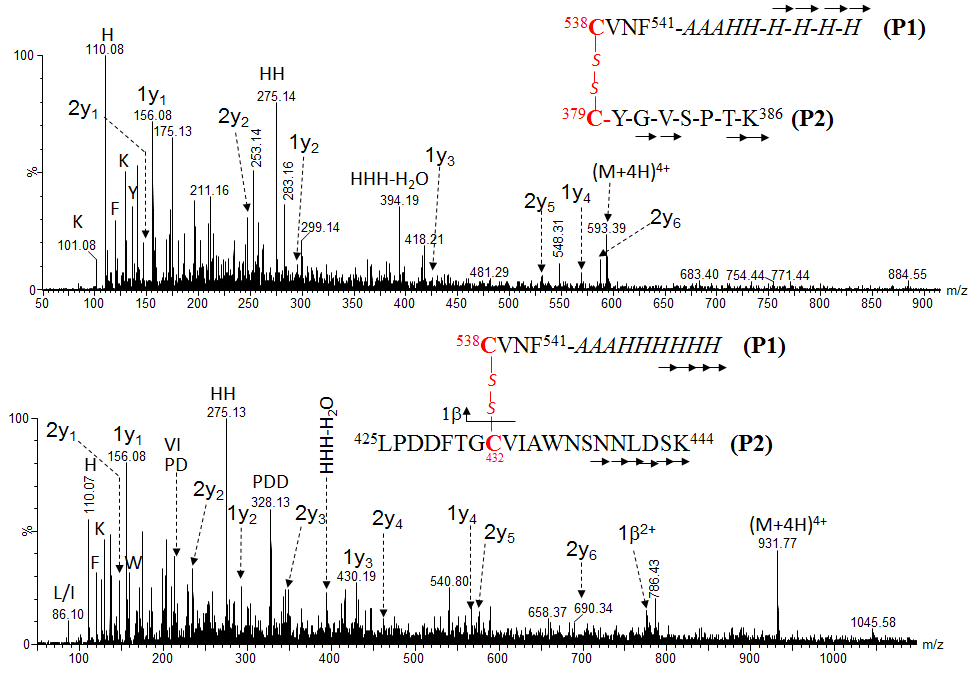

### **Fig. S5.** ESI-MS/MS spectra of peptides containing the free Cys_336_, Cys_391_, Cys_432_ and C_538_ detected in *RBD_(319-541)_-HEK_A3*.

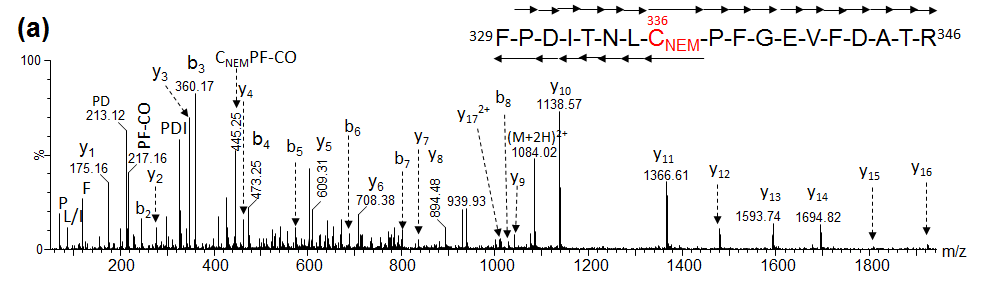

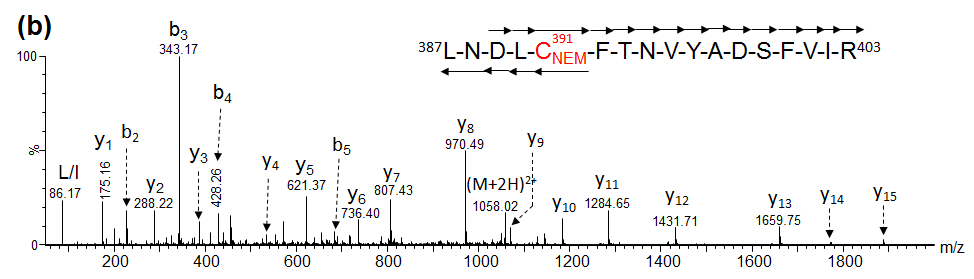

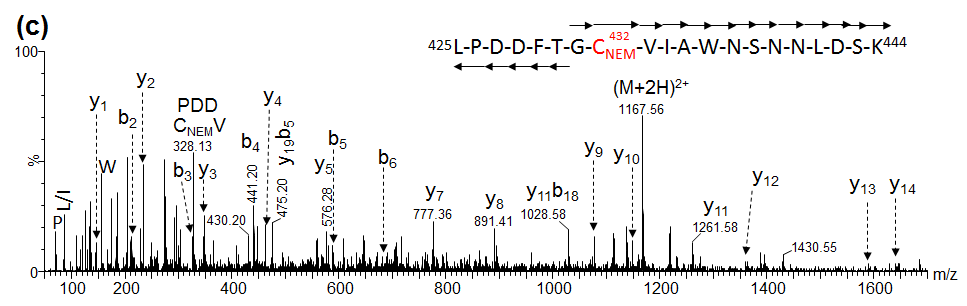

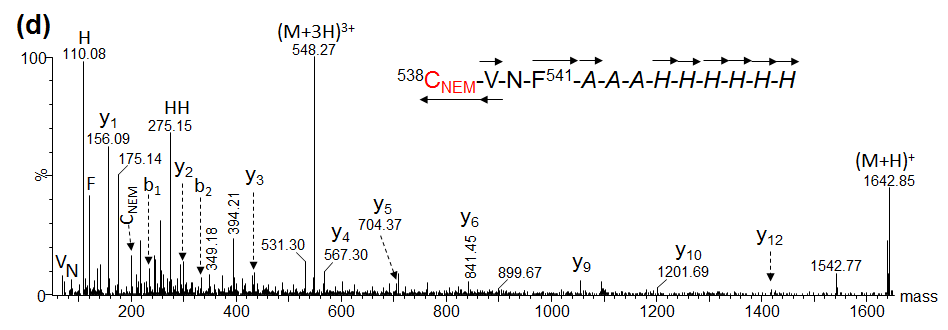

**Fig. S6.** ESI-MS multiply-charged (a) and deconvoluted with MaxEnt1 (b) of the molecular mass of the N-deglycosylated protein and the tryptic peptides obtained by using the in-solution standard (c) and BFD with ethanol (d) protocol for *RBD_(319-541)_-HEK*. The inset shown in (b), (c) and (d) corresponds to the expanded region in the range delimited by a broken line rectangle. Monosaccharide symbols follow the SNFG system (2) and the O-glycans structures as previously reported (3). The upper and lower mass spectra shown in (e), (f) and (g) correspond to expanded regions of the ESI-MS spectra shown in (c) and (d), respectively.
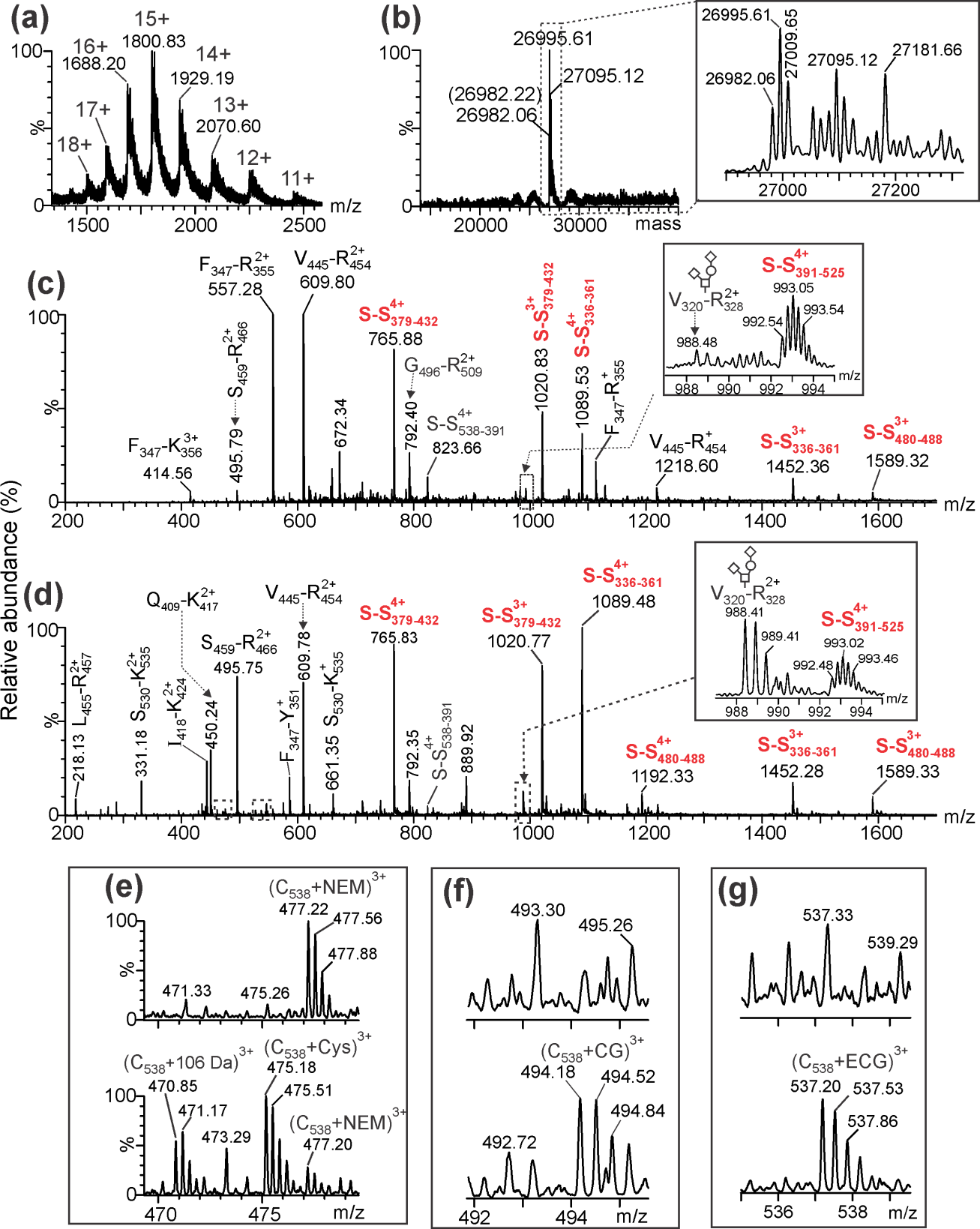

**Table S1.** Summary of the 100% sequence coverage assignment by ESI-MS of the tryptic digestion using the in-solution buffer-free (BFD) and 85% by the standard digestion (SD) protocol of *RBD_(319-541)_-HEK* expressed in HEK293T.

| **Code** | ***m/z*_Theor_** | **z** | ***m/z*_Exp_** | | **Assignment** |
| --- | --- | --- | --- | --- | --- |
|  |  |  | **BFD** | **SD** |  |
| V_320_-R_328_ | 514.79 | 2 | 514.76 | - | ^320^VQPTESIVR^328^ |
| F_347_-R_355_ | 557.28  1113.55 | 2 | 557.26  1113.47 | 557.28  1113.56 | ^347^FASVYAWNR^355^ |
| F_347_-K_356_ | 414.55 | 3 | 414.57 | 414.56 | ^347^FASVYAWNRK^356^ |
| K_356_-R_357_ | 303.21 | 1 | 303.20 | - | ^356^KR^357^ |
| G_404_-R_408_ | 575.28  288.14 | 1  2 | 575.25  288.13 | -  - | ^404^GDEVR^408^ |
| Q_409_-K_417_ | 899.50  450.25 | 1  2 | 899.44  450.24 | -  - | ^409^QIAPGQTGK^417^ |
| I_418_-K_424_ | 886.43  443.72 | 1  2 | 886.37  443.70 | -  ~~-~~ | ^418^IADYNYK^424^ |
| V_445_-R_454_ | 1218.59  609.80 | 1  2 | 1218.52  609.78 | 1218.60  609.80 | ^445^VGGNYNYLYR^454^ |
| L_455_-R_457_ | 435.27  218.14 | 1  2 | 435.25  218.13 | -  - | ^455^LFR^457^ |
| K_458_-R_466_ | 559.82  373.55 | 2  3 | 559.79  373.54 | -  373.55 | ^458^KSNLKPFER^466^ |
| S_459_-R_466_ | 495.77 | 2 | 495.75 | 495.79 | ^459^SNLKPFER^466^ |
| G_496_-R_509_ | 792.38 | 2 | 792.35 | 792.40 | ^496^GFQPTNGVGYQPYR^509^ |
| K_529_-K_535_ | 395.25 | 2 | 395.22 | - | ^529^KSTNLVK^535^ |
| S_530_-K_535_ | 661.39  331.20 | 1  2 | 661.35  331.18 | -  - | ^530^STNLVK^535^ |
| N_536_-K_537_ | 261.16 | 1 | 261.14 | - | ^536^NK^537^ |
| **O-glycopeptides** | | | | | |
| 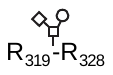 | 920.96  614.31 | 2  3 | 920.90  614.29 | -  - | ^319^RVQP**T**E**S**IVR^328^+HexNAc-Hex-NeuAc (Nt-free + O-glycosylation) |
| 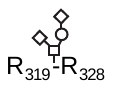 | 1066.50  711.34 | 2  3 | 1066.46  711.32 | 1066.52  711.35 | ^319^RVQP**T**E**S**IVR^328^+HexNAc-Hex-NeuAc_2_ (Nt-free + O-glycosylation) |
| 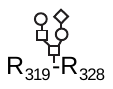 | 736.02  1103.52 | 3  2 | 735.98  1103.47 | 736.04  1103.53 | ^319^RVQPTESIVR^328^+ HexNAc_2_-Hex_2_-NeuAc (Nt-free + O-glycosylation) |
| 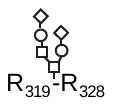 | 1249.07  833.05 | 2  3 | 1248.99  833.02 | 1249.07  833.07 | ^319^RVQPTESIVR^328^+ HexNAc_2_-Hex_2_-NeuAc_2_ (Nt-free + O-glycosylation) |
| 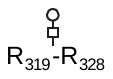 | 517.27 | 3 | 517.24 | 517.26 | ^319^RVQPTESIVR^328^ +HexNAc-Hex (O-glycosylation) |
| 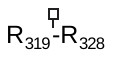 | 694.38 | 2 | 694.34 | 694.38 | ^319^RVQPTESIVR^328^ +HexNAc (O-glycosylation) |
| 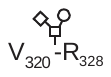 | 842.90 | 2 | 842.88 | - | ^320^VQP**T**E**S**IVR^328^+HexNAc-Hex-NeuAc (O-glycosylation) |
| 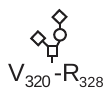 | 988.45 | 2 | 988.41 | 988.48 | ^320^VQP**T**E**S**IVR^328^+HexNAc-Hex-NeuAc_2_ (O-glycosylation) |
| 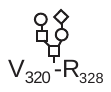 | 1025.47 | 2 | 1025.42 | - | ^320^VQPTESIVR^328^ +HexNAc_2_-Hex_2_-NeuAc (O-glycosylation) |
| 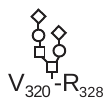 | 1171.02 | 2 | 1170.97 | - | ^320^VQPTESIVR^328^ +HexNAc_2_-Hex_2_-NeuAc_2_ (O-glycosylation) |
| 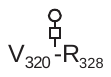 | 697.36 | 2 | 697.31 | - | ^320^VQPTESIVR^328^ +HexNAc-Hex (O-glycosylation) |
| 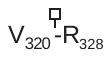 | 616.33 | 2 | 616.30 | - | ^320^VQPTESIVR^328^ +HexNAc (O-glycosylation) |
| **Native disulfide bonds** | | | | | |
| S-S_336-361_ | 1452.35  1089.51 | 3  4 | 1452.28  1089.48 | 1452.36  1089.53 | ^329^FP**D**ITNL**C**PFGEVF**D**ATR^346^  **____\|**  **\|** ^358^ISN**C**VADYSVLYNSASFSTFK^378^  **(Native C336-C361)** |
| S-S_379-432_ | 1020.81  765.86 | 3  4 | 1020.77  765.83 | 1020.83  765.88 | ^379^**C**YGVSPTK^386^  **\|**  ^425^LPDDFTG**C**VIAWNSNNLDSK^444^  **(Native C379-C432)** |
| S-S_491-525_ | 992.52 | 4 | 992.48 | 992.54 | ^387^LNDL**C**FTNVYADSFVIR^403^  **\|__________**  **\|**  ^510^VVVLSFELLHAPATV**C**GPK^528^  **(Native C391-C525)** |
| S-S_480-488_ | 1589.38  1192.29 | 3  4 | 1589.33  1192.33 | 1589.32  1192.33 | ^467^DISTEIYQAGSTP**C**NGVEGFN**C**YFPLQSYGFQPTNGVGYQPYR^509^ **\|________\|**  **(Native C480-C488)** |
| **Scrambled disulfide bonds** | | | | | |
| S-S_336_-_379_ | 965.12 | 3 | 965.06 | - | ^329^FP**D**ITNL**C**PFGEVF**D**ATR^346^  **\|**  ^379^**C**YGVSPTK^386^  **(Scrambling C336-C379)** |
| S-S_538_-_336_ | 836.63  669.51 | 4  5 | 836.61  669.50 | 836.66  669.52 | ^538^**C**VNF^541^*-HHHHHH*  **\|**  ^329^FP**D**ITNL**C**PFGEVF**D**ATR^346^  **(Scrambling C538-C336)** |
| S-S_538_-_361_ | 905.16 | 4 | 905.12 | - | ^538^**C**VNF^541^*-HHHHHH*  **\|**  ^358^ISN**C**VADYSVLYNSASFSTFK^378^  **(Scrambling C538-C361)** |
| S-S_538_-_379_ | 719.32  539.74 | 3  4 | 719.31  539.72 | 719.38  539.79 | ^538^**C**VNF^541^*-HHHHHH*  **\|**  ^379^**C**YGVSPTK^386^  **(Scrambling C538-C379)** |
| S-S_538_-_391_ | 823.63  659.11 | 4  5 | 823.61  659.09 | 823.66  659.12 | ^538^**C**VNF^541^*-HHHHHH*  **\|**  ^387^LNDL**C**FTNVYADSFVIR^403^  **(Scrambling C538-C391)** |
| S-S_538-432_ | 878.40 | 4 | 878.38 | - | ^538^**C**VNF^541^*-HHHHHH*  **\|**  ^425^LPDDFTG**C**VIAWNSNNLDSK^444^  **(Scrambling C538-C432)** |
| S-S_432-432_ | 1104.51 | 4 | 1104.46 | - | ^425^LPDDFTG**C**VIAWNSNNLDSK^444^  **\|**  ^425^LPDDFTG**C**VIAWNSNNLDSK^444^  **(Scrambling C432-C432)** |
| S-S_538-525_ | 821.16  657.13 | 4  5 | 821.14  - | 821.19  657.15 | ^538^**C**VNF^541^*-HHHHHH*  **\|**  ^510^VVVLSFELLHAPATV**C**GPK^528^  **(Scrambling C538-C525)** |
| S-S_538-538_ | 869.37  652.28 | 3  4 | 869.38  652.31 | -  - | ^538^**C**VNF^541^*-HHHHHH*  **\|**  ^538^**C**VNF^541^*-HHHHHH*  **(Homodimer C538-C538)** |
| **Free and modified cysteines** | | | | | |
| C_336_+NEM | 1084.01 | 2 | 1084.01 | 1084.03 | ^329^FP**D**ITNL**C_NEM_**PFGEVF**D**ATR^346^  **(C336+NEM, +125 Da)** |
| C_432_+NEM | 1167.54 | 2 | 1167.48 | 1167.55 | ^425^LPDDFTG**C_NEM_**VIAWNSNNLDSK^444^  **(C432+NEM, +125 Da)** |
| C_391_+NEM | 1058.02 | 2 | 1057.97 | 1058.04 | ^387^LNDL**C_NEM_**FTNVYADSFVIR^403^  **(C391+NEM, +125 Da)** |
| C_525_+NEM | 702.39 | 3 | 702.40 | 702.41 | ^510^VVVLSFELLHAPATV**C_NEM_**GPK^528^  **(C525+NEM, +125 Da)** |
| C_538_+NEM | 477.21 | 3 | 477.20 | 477.22 | ^538^**C_NEM_**VNF^541^-*HHHHHH*  (**Ct, C538+NEM, +125 Da**) |
| C_538_+Cys | 475.19 | 3 | 475.18 | - | ^538^**C**VNF^541^-*HHHHHH*  **\|**  NH_2_-**C**-COOH  (**Cys+C538, +119 Da**) |
| C_538_+ECG | 537.21  403.16 | 3  4 | 537.20  403.15 | -  - | ^538^**C**VNF^541^-*HHHHHH*  **\|**  NH_2_-E**C**G-COOH  (**C538+ECG, +305 Da**) |
| C_538_+CG | 494.20 | 3 | 494.18 | - | ^538^**C**VNF^541^-*HHHHHH*  **\|**  NH_2_-**C**G-COOH  (**C538+CG, +176 Da**) |
| C_538_+EC | 518.21 | 3 | 518.19 | - | ^538^**C**VNF^541^-*HHHHHH*  **\|**  NH_2_-E**C**-COOH  (**C538+EC, +248 Da**) |
| C_538_-34 Da | 424.20 | 3 | 424.19 | - | ^538^**C***VNF^541^-*HHHHHH*  (**C538 - 34 Da, -SH_2_**) |
| C_538_-49 Da | - | 3 | 419.17 | - | ^538^**C***VNF^541^-*HHHHHH*  (**C538 - 49 Da, unknown**) |
| C_538_+32Da | 446.19 | 3 | 446.19 | - | ^538^**C***VNF^541^-*HHHHHH*  (**C538 + 32 Da, possibly +[O]_2_**) |
| C_538_+64Da | 456.85 | 3 | 456.82 | - | ^538^**C***VNF^541^-*HHHHHH*  (**C538 + 64 Da, +S[O]_2_**) |
| C_538_+90 Da | - | 3 | 465.51 | - | ^538^**C***VNF^541^*-HHHHHH*  (**C538 + 90 Da, unknown**) |
| C_538_+106Da | - | 3 | 470.85 | ~~-~~ | ^538^**C***VNF^541^*-HHHHHH*  (**C538 + 106 Da, unknown**) |
| C_538_+147Da | - | 3 | 484.50 | - | ^538^**C***VNF^541^*-HHHHHH*  (**C538 + 147 Da, unknown**) |
| C_538_+191Da | - | 3 | 499.19 | - | ^538^**C***VNF^541^*-HHHHHH*  (**C538 + 191 Da, unknown**) |
| **Artifacts of the protocol** | | | | | |
| 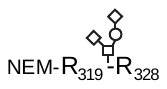 | 1129.03 | 2 | 1128.99 | 1129.05 | NEM -^319^RVQPTESIVR^328^ +HexNAc-Hex-NeuAc_2_ (Nt-NEM + O-glycosylation) |
| 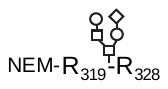 | 1166.05 | 2 | 1166.01 | 1166.07 | NEM -^319^RVQPTESIVR^328^ +HexNAc_2_-Hex_2_-NeuAc (Nt-NEM + O-glycosylation) |
| K_356_-R_357_^NEM^ | 214.63 | 2 | 214.63 | - | ^356^K_NEM_R^357^ |
| _NEM_  S-S_336-361_ | 1491.03  1120.78 | 3  4 | -  - | 1494.05  1120.80 | ^329^FP**D**ITNL**C**PFGEVF**D**ATR^346^  **___\|**  **\|** ^358^ISN**C**VADYSVLYNSASFSTFK^378^  (Native C336-C361, +NEM) |
| _NEM_  S-S_379-432_ | 1062.49  797.12 | 3  4 | -  - | 1062.51  797.12 | ^379^**C**YGVSPTK^386^  **\|**  ^425^LPDDFTG**C**VIAWNSNNLDSK^444^  (Native C379-C432, +NEM) |
| Q_409_-K_417_+NEM | 512.78 | 2 | - | 512.78 | ^409^QIAPGQTGK^417^ +NEM |
| I_418_-K_424_ +NEM | 1011.48  506.24 | 1  2 | -  - | 1011.50  506.26 | ^418^IADYNYK^424^ +NEM |
| V_445_-R_454_+NEM |  | 1  2 | -  - | 1343.64  672.34 | ^445^VGGNYNYLYR^454^ +NEM |
| L_455_-R_457_+NEM |  | 1 | - | 560.33 | ^455^LFR^457^ +NEM |
| K_458_-R_466_+NEM | 622.34 | 2 | - | 622.35 | ^458^KSNLKPFER^466^ +NEM |
| K_529_-K_535_+NEM | 457.77 | 2 | 457.75 | 457.78 | ^529^KSTNLVK^535^ +NEM |
| NEM-Cys+C_538_ | 516.88 | 3 | 516.85 | - | ^538^**C**VNF^541^-*HHHHHH*  **\|**  NEM-**C**-COOH  (NEM at Cys+C^538^, +244 Da) |
| OH-NEM-Cys+C_538_ | 522.88 | 3 | 522.85 | - | ^538^**C**VNF^541^-*HHHHHH*  **\|**  OH-NEM-**C**-COOH  (hydrolyzed NEM at Cys+C^538^, +262 Da) |

### **Table S2.** Summary of the 100% sequence coverage assignment by ESI-MS of the tryptic digestion using the in-solution buffer-free (BFD) and 80.6% by the standard digestion (SD) protocol of *(RBD_(319-541)_-CHO)_2_* expressed in CHO.

| **Code** | ***m/z*_Theor_** | **z** | ***m/z*_Exp_** | | **Assignment** |
| --- | --- | --- | --- | --- | --- |
|  |  |  | **BFD** | **SD** |  |
| R^319^-R^328^ | 592.84 | 2 | 592.82 | 592.84 | ^319^RVQPTESIVR^328^ |
| V^320^-R^328^ | 514.79 | 2 | 514.78 | - | ^320^VQPTESIVR^328^ |
| F^347^-R^355^ | 557.28  1113.55 | 2  1 | 557.28  1113.54 | 557.28  1113.50 | ^347^FASVYAWNR^355^ |
| F^347^-K^356^ | 621.33 | 2 | 621.30 | - | ^347^FASVYAWNRK^356^ |
| K^356^-R^357^ | 303.21 | 1 | 303.21 | - | ^356^KR^357^ |
| G^404^-R^408^ | 575.28  288.14 | 1  2 | 575.28  288.14 | -  - | ^404^GDEVR^408^ |
| Q^409^-K^417^ | 899.50  450.25 | 1  2 | 899.48  450.25 | -  - | ^409^QIAPGQTGK^417^ |
| I^418^-K^424^ | 886.43  443.72 | 1  2 | 886.41  443.72 | -  443.72 | ^418^IADYNYK^424^ |
| V^445^-R^454^ | 1218.59  609.80 | 1  2 | 1218.55  609.79 | 1218.56  609.80 | ^445^VGGNYNYLYR^454^ |
| L^455^-R^457^ | 435.27  218.14 | 1  2 | 435.27  218.14 | -  - | ^455^LFR^457^ |
| K^458^-R^466^ | 373.55 | 3 | 373.54 | 373.54 | ^458^KSNLKPFER^466^ |
| S^459^-R^466^ | 495.77 | 2 | 495.77 | 495.77 | ^459^SNLKPFER^466^ |
| K^529^-K^535^ | 395.25 | 2 | 395.24 | - | ^529^KSTNLVK^535^ |
| S530-K535 | 661.39  331.20 | 1  2 | 661.38  331.19 | -  - | ^530^STNLVK^535^ |
| N^536^-K^537^ | 261.16 | 1 | 261.15 | - | ^536^NK^537^ |
| **O-glycopeptides** | | | | | |
| 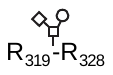 | 920.96  614.31 | 2  3 | 920.94  614.30 | 920.96  614.31 | ^319^RVQPTESIVR^328^+HexNAc-Hex-NeuAc (Nt-free + O-glycosylation) |
| 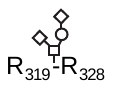 | 1066.50  711.34 | 2  3 | 1066.50  711.33 | 1066.50  711.34 | ^319^RVQPTESIVR^328^+ HexNAc-Hex-NeuAc_2_ (Nt-free + O-glycosylation) |
| 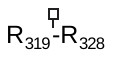 | 694.38 | 2 | 694.37 | 694.39 | ^319^RVQPTESIVR^328^ +HexNAc (O-glycosylation) |
| 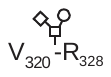 | 842.90 | 2 | 842.90 | 842.91 | ^320^VQPTESIVR^328^+HexNAc-Hex-NeuAc (O-glycosylation) |
| 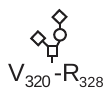 | 988.45 | 2 | 988.44 | 988.46 | ^320^VQPTESIVR^328^+HexNAc-Hex-NeuAc_2_ (O-glycosylation) |
| 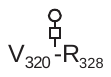 | 697.36 | 2 | 697.35 | - | ^320^VQPTESIVR^328^ +HexNAc-Hex (O-glycosylation) |
| 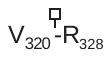 | 616.33 | 2 | 616.31 | - | ^320^VQPTESIVR^328^ +HexNAc (O-glycosylation) |
| **Expected disulfide bonds** | | | | | |
| **S-S_336-361_** | 1452.35  1089.51 | 3  4 | 1452.32  1089.50 | 1452.34  1089.51 | ^329^FP**D**ITNL**C**PFGEVF**D**ATR^346^  **____\|**  **\|** ^358^ISN**C**VADYSVLYNSASFSTFK^378^  (Native C^336^-C^361^) |
| **S-S_336-361_** | 1297.93 | 3 | - | 1297.95 | ^329^FP**D**ITNL**C**PFGEVF**D**ATR^346^  **\|**  ^358^ISN**C**VADYNSASF^370^  (Native C336-C361, *in-source fragmentation*) |
| **S-S_336-361_** | 975.11 | 3 | - | 975.12 | ^329^FP**D**ITNL**C**PFGEVF**D**ATR^346^  **\|**  ^358^ISN**C**VADY^365^  (Native C336-C361, *in-source fragmentation*) |
| **S-S_379-432_** | 1530.71  1020.81  765.86 | 2  3  4 | 1530.70  1020.80  765.86 | 1530.70  1020.81  765.86 | ^379^**C**YGVSPTK^386^  **\|**  ^425^LPDDFTG**C**VIAWNSNNLDSK^444^  (Native C379-C432) |
| **S-S_391-525_** | 992.52  794.22 | 4  5 | 992.51  794.20 | 992.52  - | ^387^LNDL**C**FTNVYADSFVIR^403^  **\|__________**  **\|**  ^510^VVVLSFELLHAPATV**C**GPK^528^  (Native C391-C525) |
| **S-S_480-488_** | 1589.38 | 3 | 1589.37 | 1589.39 | ^467^DISTEIYQAGSTP**C**NGVEGFN**C**YFPLQSYGFQPTNGVGYQPYR^509^ **\|________\|**  (Native C480-C488) |
| **S-S_538-538_** | 652.28  522.03 | 4  5 | 652.29  522.01 | -  - | ^538^**C**VNF^541^*-HHHHHH*  **\|**  ^538^**C**VNF^541^*-HHHHHH*  (Ct homodimer, C538-C538) |
| **Artifacts of the protocol** | | | | | |
| 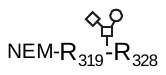 | 983.48 | 2 | 983.46 | - | NEM-^319^RVQPTESIVR^328^ +GalNAc-Gal-SA (Nt-NEM + O-glycosylation) |
| 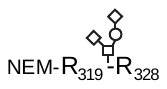 | 1129.03 | 2 | 1129.04 | - | NEM-^319^RVQPTESIVR^328^ +GalNAc-Gal-SA_2_ (Nt-NEM + O-glycosylation) |
| 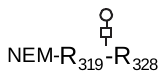 | 837.93 | 2 | 837.92 | - | NEM-^319^RVQPTESIVR^328^ +HexNAc-Hex (O-glycosylation) |
| 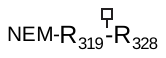 | 756.91 | 2 | 756.90 | - | NEM-^319^RVQPTESIVR^328^ +HexNAc (O-glycosylation) |
| V^445^-R^454^ +40 Da | 629.81 | 2 | 629.79 | - | ^445^VGGNYNYLYR^454^ +40 Da at Gly^446^ and Nt |
| K^458^-R^466^ +NEM | 622.34 | 2 | 622.35 | 622.34 | ^458^KSNLKPFER^466^ +NEM |
| K^529^-K^535^ +NEM | 457.77 | 2 | 457.75 | 457.76 | ^529^KSTNLVK^535^ +NEM |

### **Table S3.** Summary of the 99% sequence coverage assignment by ESI-MS of the tryptic digestion using the in-solution buffer-free (BFD) protocol of *RBD_(331-529)_-Ec* expressed in *E. coli*.

| **Code** | ***m/z*_Exp_** | **z** | ***m/z*_Theor_** | **Assignment** |
| --- | --- | --- | --- | --- |
| Nt-His_6_ | 884.93  590.29  442.97 | 2  3  4 | 884.93  590.29  442.97 | *GSSHHHHHHSSGLVPR*  (N-terminal end with His_6_-tag) |
| F_347_-R_355_ | 557.28 | 2 | 557.28 | ^347^FASVYAWNR^355^ |
| K_356_-R_357_ | 303.20 | 1 | 303.21 | ^356^KR^357^ |
| G_404_-R_408_ | 575.28  288.14 | 1  2 | 575.28  288.14 | ^404^GDEVR^408^ |
| Q_409_-K_417_ | 450.25  899.48 | 2  1 | 450.25  899.50 | ^409^QIAPGQTGK^417^ |
| I_418_-K_424_ | 443.72 | 2 | 443.72 | ^418^IADYNYK^424^ |
| V_445_-R_454_ | 609.80 | 2 | 609.80 | ^445^VGGNYNYLYR^454^ |
| L_455_-R_457_ | 435.27  218.14 | 1  2 | 435.27  218.14 | ^455^LFR^457^ |
| S_459_-R_466_ | 495.77  330.84 | 2  3 | 495.77  330.85 | ^459^SNLKPFER^466^ |
| **Native disulfide bonds** | | | | |
| **S-S_336-361_** | 1560.37  1170.58  936.61 | 3  4  5 | 1560.39  1170.55  936.64 | *GSHMAS*-^331^NITNL**C**PFGEVFNATR^346^  **_________\|**  **\|** ^358^ISN**C**VADYSVLYNSASFSTFK^378^  **(Native C^336^-C^361^)** |
| **S-S_379-432_** | 1530.72  1020.81  765.86 | 23  4 | 1530.71  1020.81  765.86 | ^379^**C**YGVSPTK^386^  **\|**  ^425^LPDDFTG**C**VIAWNSNNLDSK^444^  **(Native C^379^-C^432^)** |
| **S-S_480-488_** | 1590.05 | 3 | 1590.04 | ^467^DISTEIYQAGSTP**C**NGVEGFN**C**YFPLQSYGFQPTNGVGYQPYR^509^ **\|________\|**  **(Native C^480^-C^488^** **+ two deamidated residues at Asn^501^ and tentatively Asn^481^)** |
| **S-S_391-525_** | 992.51 | 4 | 992.52 | ^387^LNDL**C**FTNVYADSFVIR^403^  **\|**  ^510^VVVLSFELLHAPATV**C**GPK^528^  **(Native C^391^-C^525^)** |
| **Scrambled disulfide bonds** | | | | |
| S-S_336-336_ | 1183.06  946.66 | 4  5 | 1183.05  946.64 | *GSHMAS*-^331^NITNL**C**PFGEVFNATR^346^  **\|**  *GSHMAS*-^331^NITNL**C**PFGEVFNATR^346^  (**Homodimer scrambling C^336^-C^336^)** |
| S-S_336-379_ | 1073.17  805.14 | 3  4 | 1073.17  805.13 | *GSHMAS*-^331^NITNL**C**PFGEVFNATR^346^  **\|**  ^379^**C**YGVSPTK^386^  (**Scrambling C^336^-C^379^)** |
| S-S_336-391_ | 1451.71  1088.95  871.43 | 3  4  5 | 1451.69  1089.02  871.42 | *GSHMAS*-^331^NITNL**C**PFGEVFNATR^346^  **\|**  ^387^LNDL**C**FTNVYADSFVIR^403^  (**Scrambling C^336^-C^391^)** |
| S-S_336-432_ | 1524.71  1143.80  915.22 | 3  4  5 | 1524.71  1143.78  915.23 | *GSHMAS*-^331^NITNL**C**PFGEVFNATR^346^  **\|**  ^425^LPDDFTG**C**VIAWNSNNLDSK^444^  (**Scrambling C^336^-C^432^)** |
| S-S_361-379_ | 1056.50  792.63 | 3  4 | 1056.50  792.62 | ^358^ISN**C**VADYSVLYNSASFSTFK^378^  **\|**  ^379^**C**YGVSPTK^386^  (**Scrambling C^361^-C^379^)** |
| S-S_361-391_ | 1435.03  1076.53 | 3  4 | 1435.02  1076.52 | ^358^ISN**C**VADYSVLYNSASFSTFK^378^  **\|**  ^387^LNDL**C**FTNVYADSFVIR^403^  (**Scrambling C^361^-C^391^)** |
| S-S_361-432_ | 1507.96  1131.33 | 3  4 | 1508.03  1131.28 | ^358^ISN**C**VADYSVLYNSASFSTFK^378^  **\|**  ^425^LPDDFTG**C**VIAWNSNNLDSK^444^  (**Scrambling C^361^-C^432^)** |
| S-S_361-525_ | 1074.03 | 4 | 1074.05 | ^358^ISN**C**VADYSVLYNSASFSTFK^378^  **\|____________**  **\|**  ^510^VVVLSFELLHAPATV**C**GPK^528^  (**Scrambling C^361^-C^525^)** |
| S-S_379-391_ | 947.81  711.08 | 3  4 | 947.79  711.10 | ^379^**C**YGVSPTK^386^  **\|**  ^387^LNDL**C**FTNVYADSFVIR^403^  (**Scrambling** **C^379^-C^391^)** |
| S-S_379-525_ | 708.66 | 4 | 708.63 | ^379^**C**YGVSPTK^386^  **\|**  ^510^VVVLSFELLHAPATV**C**GPK^528^  (**Scrambling C^379^-C^525^)** |
| S-S_391-391_ | 995.00 | 4 | 994.99 | ^387^LNDL**C**FTNVYADSFVIR^403^  **\|**  ^387^LNDL**C**FTNVYADSFVIR^403^  (**Homodimer** **scrambling C^391^-C^391^)** |
| S-S_391-432_ | 1399.38  1049.75 | 3  4 | 1399.33  1049.75 | ^387^LNDL**C**FTNVYADSFVIR^403^  **\|**  ^425^LPDDFTG**C**VIAWNSNNLDSK^444^  **(Scrambling C^391^-C^432^)** |
| S-S^4^_32-525_ | 1396.00  1047.28 | 3  4 | 1396.04  1047.28 | ^425^LPDDFTG**C**VIAWNSNNLDSK^444^  **\|**  ^510^VVVLSFELLHAPATV**C**GPK^528^  (**Scrambling C^432^-C^525^)** |
| **Artifacts of the protocol** | | | | |
| V_445_-R_454_ +40Da | 629.80 | 2 | 629.80 | ^445^VGGNYNYLYR^454^ +40 Da at Gly^446^ |

**Fig. S7.** ESI-MS/MS spectra of the scrambled species detected in *RBD_(331-529)_-Ec* (**Table S3**). The nomenclature of fragment ions for disulfide bonds is in agreement with the proposed by Mormann et al (1).

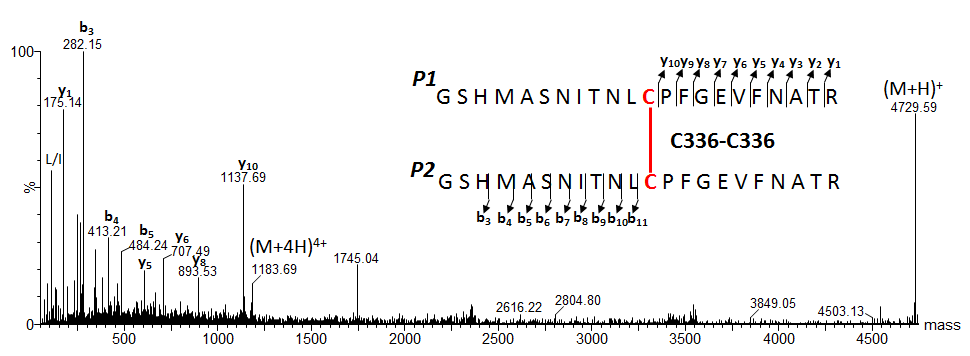

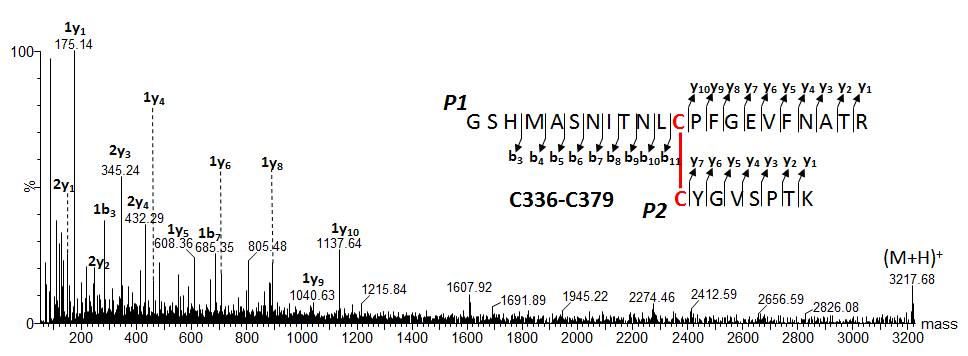

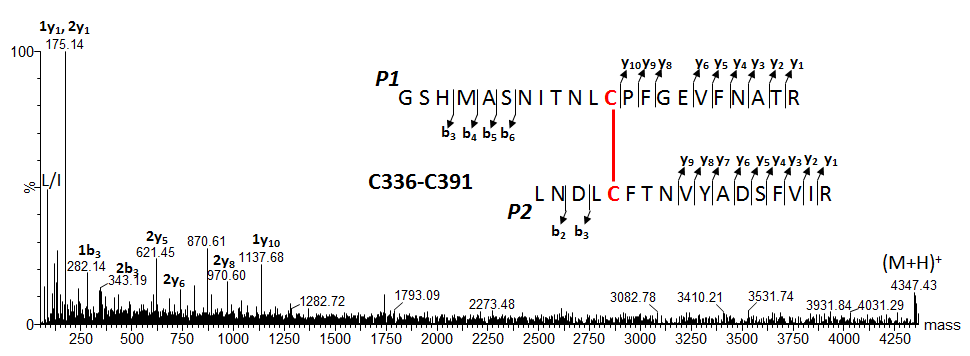

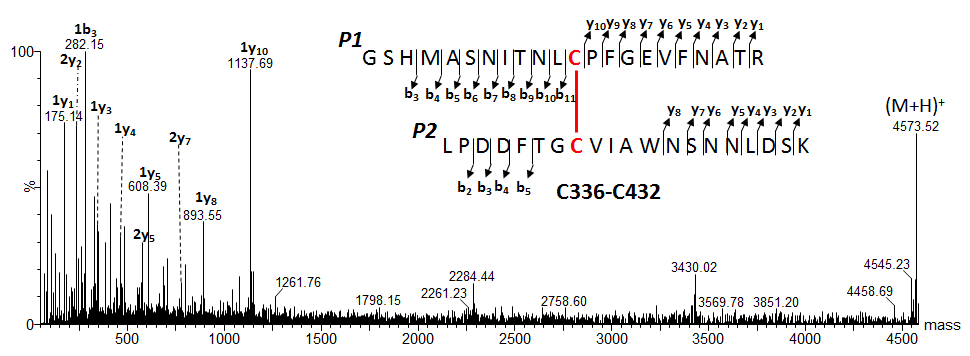

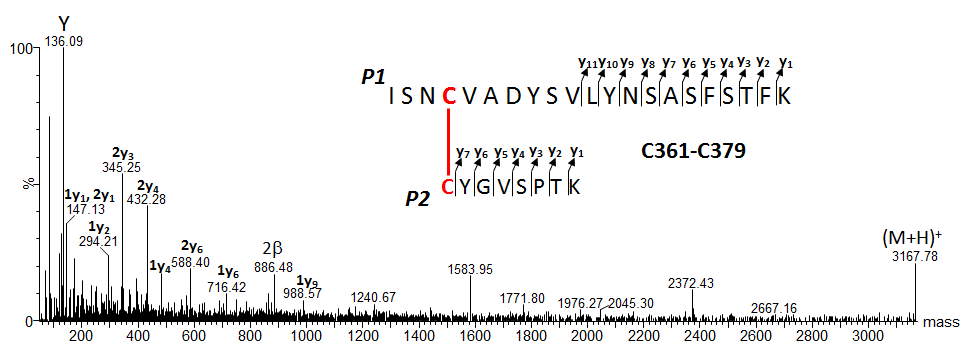

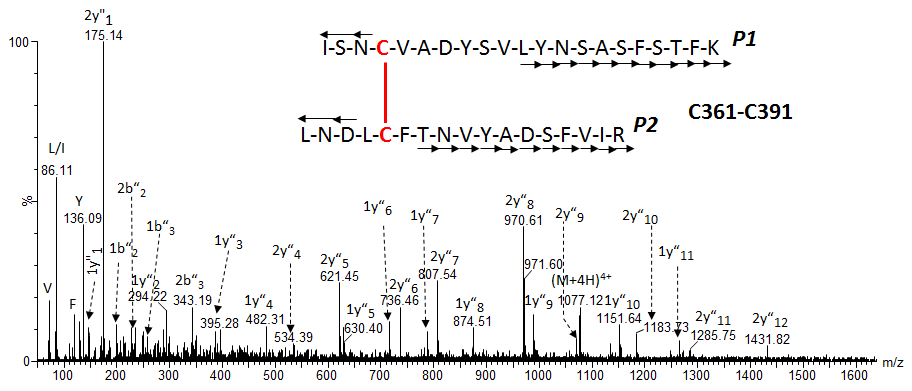

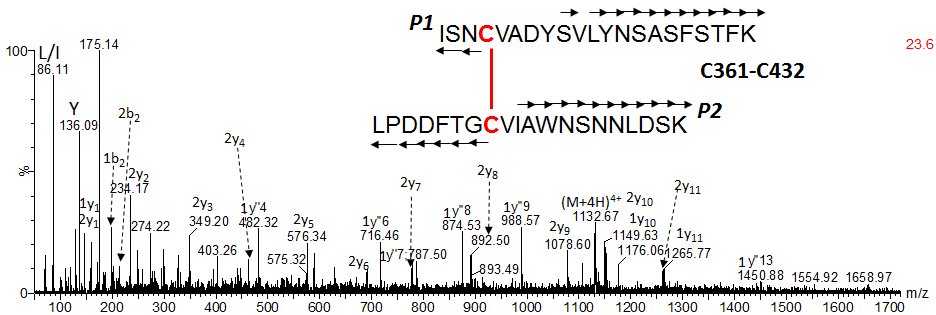

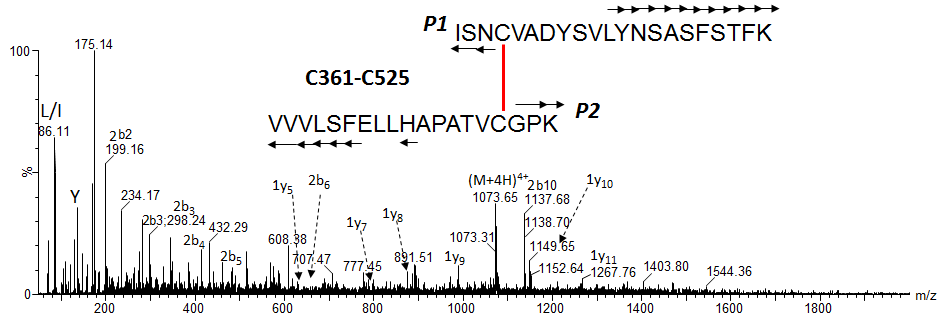

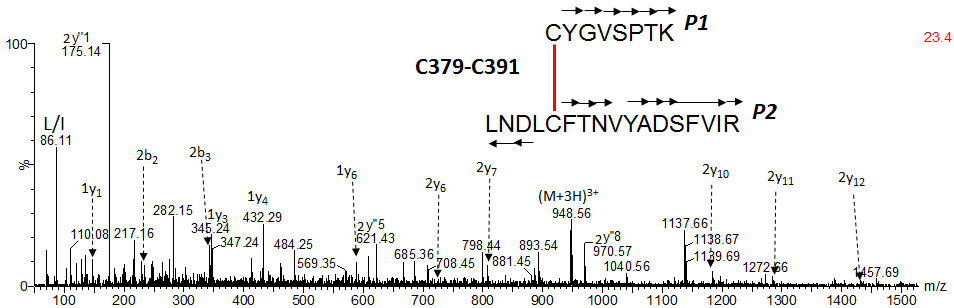

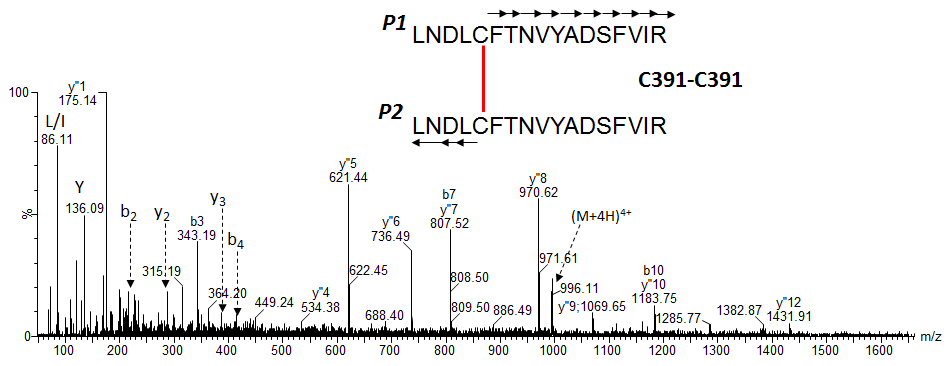

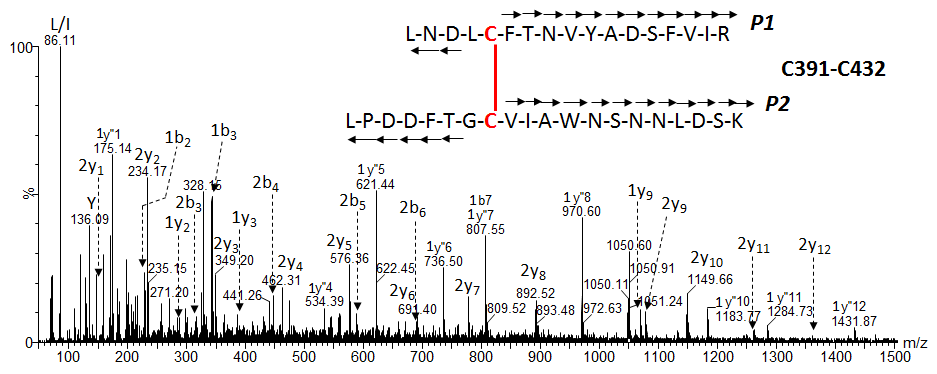

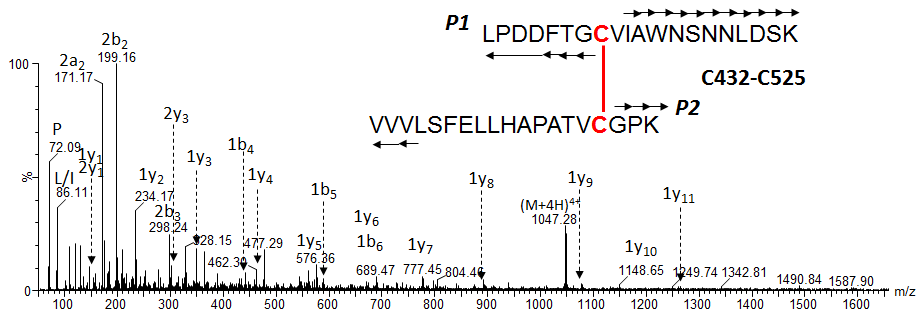

### **Fig. S8.** ESI-MS spectra of the N-deglycosylated protein before (a) and after (b) deconvolution with MaxEnt1 for the *RBD_(333-527)_-C1* (**Table 2**).

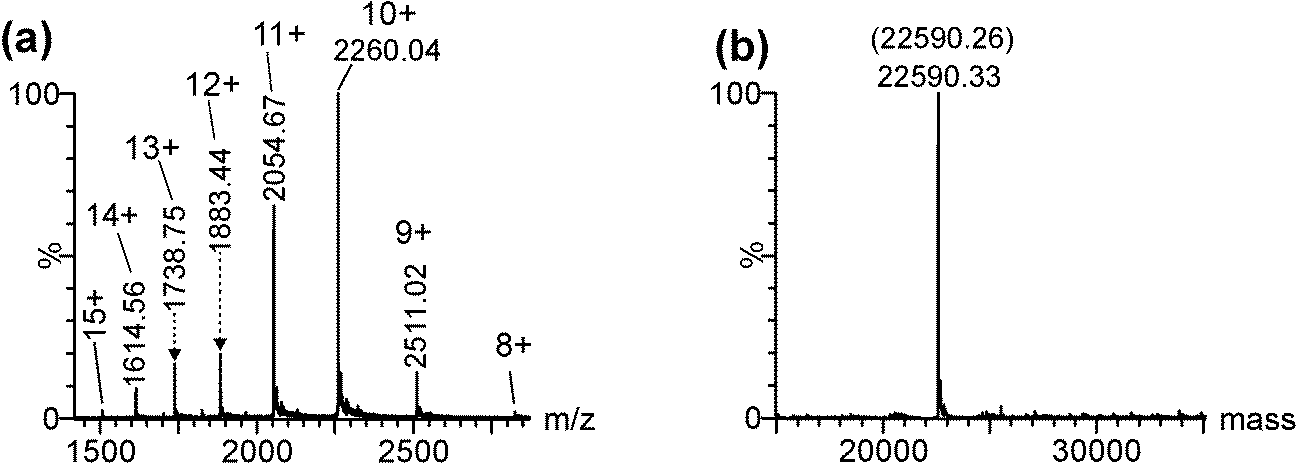

### **Table S4.** Summary of the 100% sequence coverage assignment by ESI-MS of the tryptic digestion using the in-solution BFD protocol of *RBD_(333-527)_-C1* expressed in *T. heterothallica* C1.

| **Code** | ***m/z*_Exp_** | **z** | ***m/z*_Theor_** | **Assignment** |
| --- | --- | --- | --- | --- |
| F_347_-R_355_ | 557.26 | 2 | 557.28 | ^347^FASVYAWNR^355^ |
| K_356_-R_357_ | 303.21 | 1 | 303.21 | ^356^KR^357^ |
| G_404_-R_408_ | 575.26  288.13 | 1  2 | 575.28  288.14 | ^404^GDEVR^408^ |
| Q_409_-K_417_ | 899.47  450.24 | 1  2 | 899.50  450.25 | ^409^QIAPGQTGK^417^ |
| I_418_-K_424_ | 886.40  443.71 | 1  2 | 886.43  443.72 | ^418^IADYNYK^424^ |
| V_445_-R_454_ | 609.78  1218.54 | 2  1 | 609.80  1218.59 | ^445^VGGNYNYLYR^454^ |
| L_455_-R_457_ | 218.13  435.25 | 2  1 | 218.14  435.27 | ^455^LFR^457^ |
| K_458_-R_466_ | 559.79  373.54 | 2  3 | 559.82  373.55 | ^458^KSNLKPFER^466^ |
| S_459_-R_466_ | 495.75  330.84 | 2  3 | 495.77  330.85 | ^459^SNLKPFER^466^ |
| **Native disulfide bonds** | | | | |
| **S-S_336-361_** | 1941.86  1294.92  971.44 | 2  3  4 | 1941.90  1294.94  971.46 | ^333^TNL**C**PFGEVFDATR^346^ ^(a)^  **\|**  ^358^ISN**C**VADYSVLYNSASFSTFK^378^  (N-terminal end, Native C_336_-C_361_, Asn_343_🡪Asp) |
| **S-S_379-432_** | 1530.70  1020.78  765.83 | 23  4 | 1530.71  1020.81  765.86 | ^379^**C**YGVSPTK^386^  **\|**  ^425^LPDDFTG**C**VIAWNSNNLDSK^444^  (Native C_379_-C_432_) |
| **S-S_480-488_** | 1589.36  1192.26 | 3  4 | 1589.38  1192.29 | ^467^DISTEIYQAGSTP**C**NGVEGFN**C**YFPLQSYGFQPTNGVGYQPYR^509^ **\|_________\|**  (Native C_480_-C_488_) |
| **S-S_391-525_** | 1527.41  1145.79 | 3  4 | 1527.42  1145.82 | ^387^LNDL**C**FTNVYADSFVIR^403^  **\|**  ^510^VVVLSFELLHAPATV**C**GP^527^-*GGGGSEPEA*  (C-terminal end, Native C_391_-C_525_) |
| **Free-Cys and scrambled disulfide bonds of N-deglycosylated protein** | | | | |
| C_336_+NEM | 847.87 | 2 | 847.90 | ^333^TNL**C_NEM_**PFGEVFDATR^346 (b)^ C_336_+NEM |
| C_432_+NEM | 1167.52 | 2 | 1167.54 | ^425^LPDDFTG**C_NEM_**VIAWNSNNLDSK^444 (b)^ C_432_+NEM |
| S-S_336-391_ | 889.72 | 4 | 889.68 | ^333^TNL**C**PFGEVFDATR^346^ ^(a)^  **\|**  ^387^LNDL**C**FTNVYADSFVIR^403^  **Scrambling C_336_-C_391_** |
| S-S_336-432_ | 944.43 | 4 | 944.44 | ^333^TNL**C**PFGEVFDATR^346^ ^(a)^  **\|**  ^425^LPDDFTG**C**VIAWNSNNLDSK^444^  **Scrambling C_336_-C_432_** |
| S-S_361-432_ | 1131.26 | 4 | 1131.28 | ^358^ISN**C**VADYSVLYNSASFSTFK^378^  **\|**  ^425^LPDDFTG**C**VIAWNSNNLDSK^444^  **Scrambling C_361_-C_432_** |
| **Glycopeptides of RBD with Cys reduced and carbamidomethylated** ^(a, b)^ | | | | |
| N_343_+M3 | 1259.58 | 2 | 1259.55 | ^333^TNLC_CAM_PFGEVFNATR^346^+GlcNAc_2_Man_3_ |
| N_343_+M3A1 | 1361.10 | 2 | 1361.09 | ^333^TNLC_CAM_PFGEVFNATR^346^+GlcNAc_2_Man_3_GlcNAc |
| N_343_+M4 | 1340.58 | 2 | 1340.58 | ^333^TNLC_CAM_PFGEVFNATR^346^+GlcNAc_2_Man_4_ |
| N_343_+M4A1 | 1442.11  961.76 | 2  3 | 1442.12  961.75 | ^333^TNLC_CAM_PFGEVFNATR^346^+GlcNAc_2_Man_4_GlcNAc |
| N_343_+M5A1 | 1523.13  1015.81 | 2  3 | 1523.14  1015.76 | ^333^TNLC_CAM_PFGEVFNATR^346^+GlcNAc_2_Man_5_GlcNAc |
| N_343_+M6 | 1502.58 | 2 | 1502.63 | ^333^TNLC_CAM_PFGEVFNATR^346^+GlcNAc_2_Man_6_ |
| N_343_+M7 | 1583.70 | 2 | 1583.66 | ^333^TNLC_CAM_PFGEVFNATR^346^+GlcNAc_2_Man_7_ |
| N_343_+M7A1 | 1123.79 | 3 | 1123.80 | ^333^TNLC_CAM_PFGEVFNATR^346^+GlcNAc_2_Man_7_GlcNAc |
| N_343_+M8 | 1110.12 | 3 | 1110.12 | ^333^TNLC_CAM_PFGEVFNATR^346^+GlcNAc_2_Man_8_ |
| N_343_+M8A1 | 1177.84 | 3 | 1177.82 | ^333^TNLC_CAM_PFGEVFNATR^346^+GlcNAc_2_Man_8_GlcNAc |
| N_343_+M9 | 1164.10 | 3 | 1164.14 | ^333^TNLC_CAM_PFGEVFNATR^346^+GlcNAc_2_Man_9_ |
| C_361_+CAM | 1187.04  791.92 | 2  3 | 1187.06  791.71 | ^358^ISNC_CAM_VADYSVLYNSASFSTFK^378^ |
| C_379_+CAM | 911.44  456.22 | 1  2 | 911.43  456.22 | ^379^C_CAM_YGVSPTK^386^ |
| C_391_+CAM | 1024.02 | 2 | 1024.00 | ^387^LNDLC_CAM_FTNVYADSFVIR^403^ |
| C_432_+CAM | 1133.52  756.03 | 2  3 | 1133.53  756.02 | ^425^LPDDFTGC_CAM_VIAWNSNNLDSK^444^ |
| C_480_+CAM, C_488_+CAM | 1628.07  1221.31 | 3  4 | 1628.07  1221.30 | ^467^DISTEIYQAGSTPC_CAM_NGVEGFNC_CAM_YFPLQSYGFQPTNGVGYQPYR^509^ |
| C_525_+CAM | 1325.65  884.12 | 2  3 | 1325.66  884.11 | ^510^VVVLSFELLHAPATVC_CAM_GP^527^-*GGGGSEPEA* |
| **Artifacts of the protocol** | | | | |
| V_445_-R_454_+40Da | 629.79  1258.57 | 2  1 | 629.81  1258.62 | ^445^VGGNYNYLYR^454^ +40Da at Gly^446^ |

(a) Underlined Asp (D) indicates the conversion of Asn to Asp due to N-deglycosylation of PNGase F, while underlined Asn (N) indicates the occupy N-glycosylation site of Asn^343^.
(b) C_NEM_: cysteine N-ethylmaleimydated; C_CAM_: cysteine reduced and carbamidomethylated; Man: Mannose; GlcNAc: N-acetylglucosamine.

**Fig. S9.** ESI-MS/MS spectra of the free Cys and scrambled disulfide bonds species detected in *RBD_(333-527)_-C1* (**Table S4**). The nomenclature of fragment ions for disulfide bonds is in agreement with the proposed by Mormann et al (1).

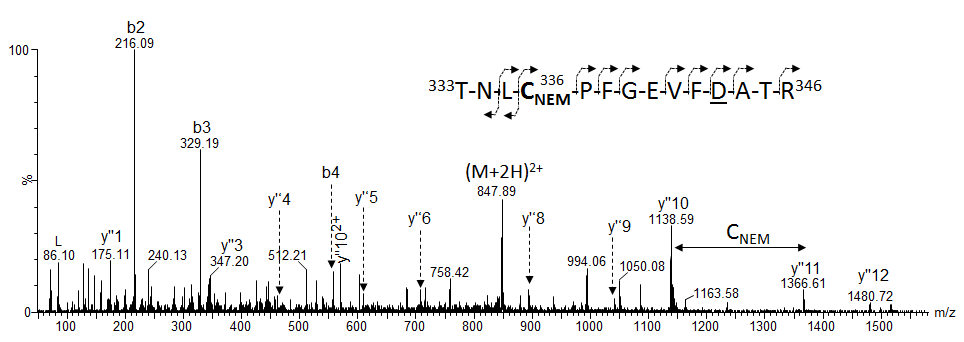

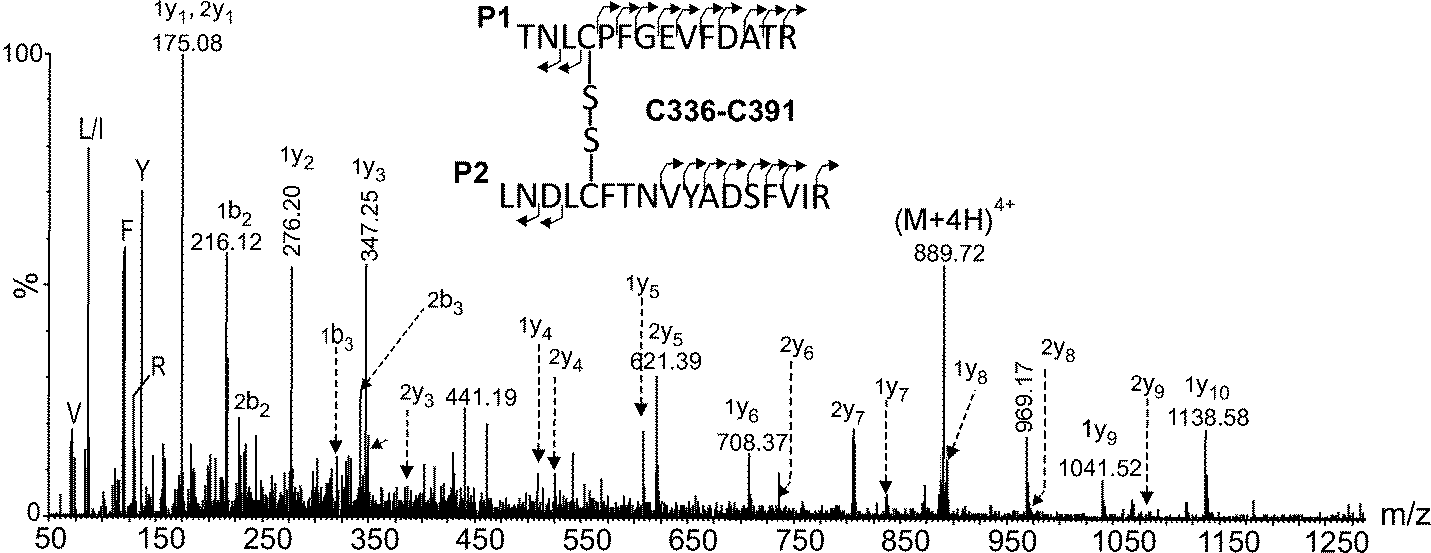

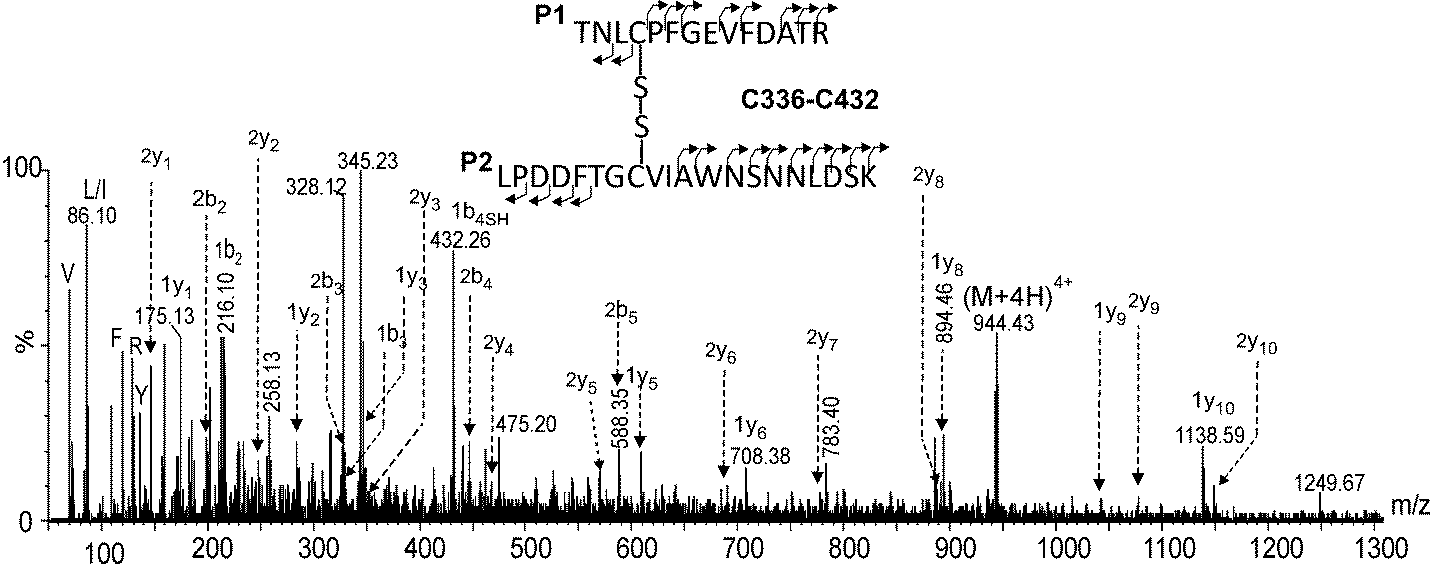

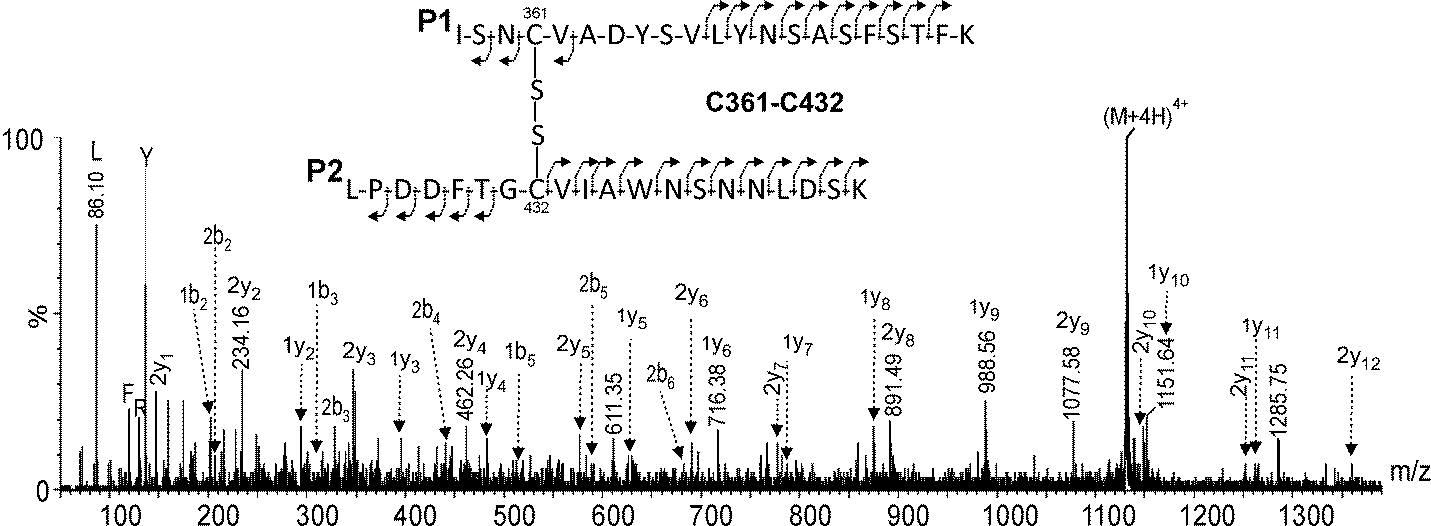

### **Fig. S10.** ESI-MS/MS spectra of the carbamidomethylated Cys peptides and the N-terminal peptide T_333_-R_346_ containing several N-glycoforms detected in *RBD_(333-527)_-C1* (**Table S4**).

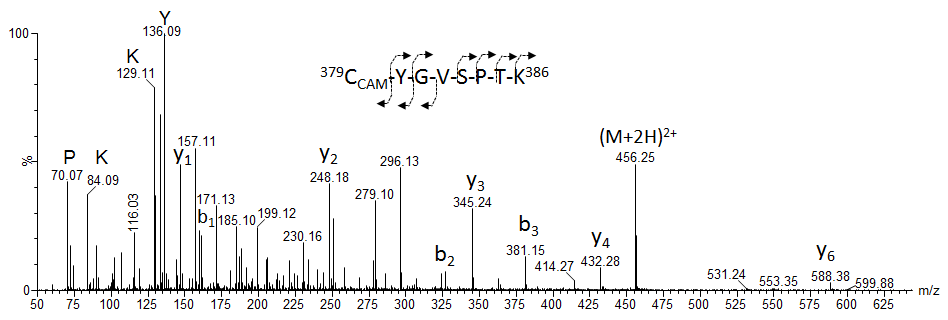

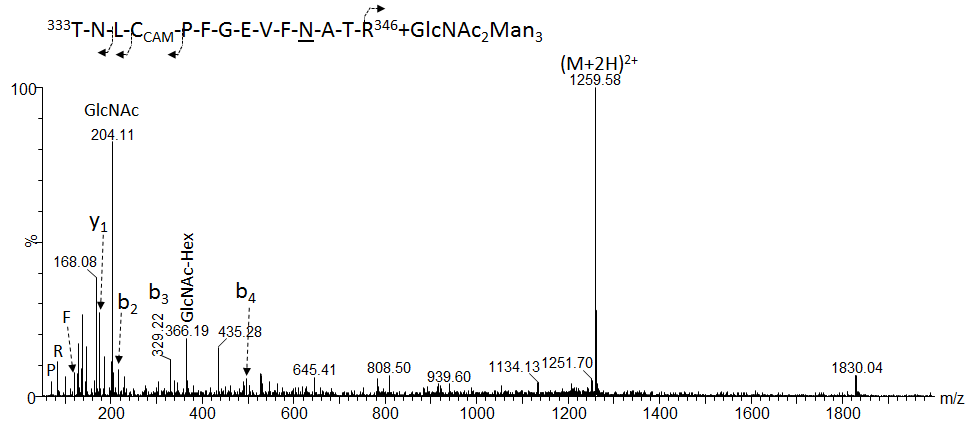

### **Table S5.** Summary of the 99% sequence coverage assignment by ESI-MS of the tryptic digestion using the in-solution BFD protocol of *RBD_(331-530)_-Cmyc-Pp* expressed in *P. pastoris*.

| **Code** | ***m/z*_Exp_** | **z** | ***m/z*_Theor_** | **Assignment** |
| --- | --- | --- | --- | --- |
| F_347_-R_355_ | 557.28 | 2 | 557.28 | ^347^FASVYAWNR^355^ |
| K_356_-R_357_ | 303.21 | 1 | 303.21 | ^356^KR^357^ |
| G_404_-R_408_ | 575.28  288.15 | 1  2 | 575.28  288.14 | ^404^GDEVR^408^ |
| Q_409_-K_417_ | 899.48  450.26 | 1  2 | 899.50  450.25 | ^409^QIAPGQTGK^417^ |
| I_418_-K_424_ | 886.44  443.72 | 1  2 | 886.43  443.72 | ^418^IADYNYK^424^ |
| V_445_-R_454_ | 609.81 | 2 | 609.80 | ^445^VGGNYNYLYR^454^ |
| L_455_-R_457_ | 218.15  435.28 | 2  1 | 218.14  435.27 | ^455^LFR^457^ |
| S_459_-R_466_ | 495.78  330.85 | 2  3 | 495.77  330.85 | ^459^SNLKPFER^466^ |
| S_530_-R_531_ | 262.15 | 1 | 262.15 | ^530^S*R*^531^ |
| EQK | 404.22 | 1 | 404.21 | *EQK* |
| Ct-His_6_ | 709.67  532.50 | 3  4 | 709.66  532.50 | *LISEEDLNSAVDHHHHHH*  (C-terminal end with His_6_-tag) |
| **Native disulfide bonds** | | | | |
| **S-S_336-361_** | 1625.42  1219.32 | 3  4 | 1625.41  1219.31 | *EAEAEFS*-^331^DITNL**C**PFGEVFDATR^346^  **___________\|**  **\|** ^358^ISN**C**VADYSVLYNSASFSTFK^378^  (N-terminal end + EAEA, Native C^336^-C^361^ and Asn^331^, ^343^🡪Asp) |
| **S-S_379-432_** | 1530.73  1020.81  765.87 | 23  4 | 1530.71  1020.81  765.86 | ^379^**C**YGVSPTK^386^  **\|**  ^425^LPDDFTG**C**VIAWNSNNLDSK^444^  (Native C^379^-C^432^) |
| **S-S_480-488_** | 1589.40  1192.30 | 3  4 | 1589.38  1192.29 | ^467^DISTEIYQAGSTP**C**NGVEGFN**C**YFPLQSYGFQPTNGVGYQPYR^509^ **\|______ ____\|**  (Native C^480^-C^488^) |
| **S-S_391-525_** | 992.49 | 4 | 992.52 | ^387^LNDL**C**FTNVYADSFVIR^403^  **\|**  ^510^VVVLSFELLHAPATV**C**GPK^528^  (Native C^391^-C^525^) |
| **Artifacts of the protocol** | | | | |
| V_445_-R_454_+40Da | 629.82  1258.60 | 2  1 | 629.81  1258.62 | ^445^VGGNYNYLYR^454^ +40Da at Gly^446^ |

### **Fig. S11.** ESI-MS/MS spectra of the NEM-R^319^-R^328^ peptide containing NEM at N-terminal end and the O-glycans HexNAc-Hex-NeuAc (a) and HexNAc-Hex-NeuAc_2_ (b) detected in *RBD_(319-541)_-HEK_A3*.

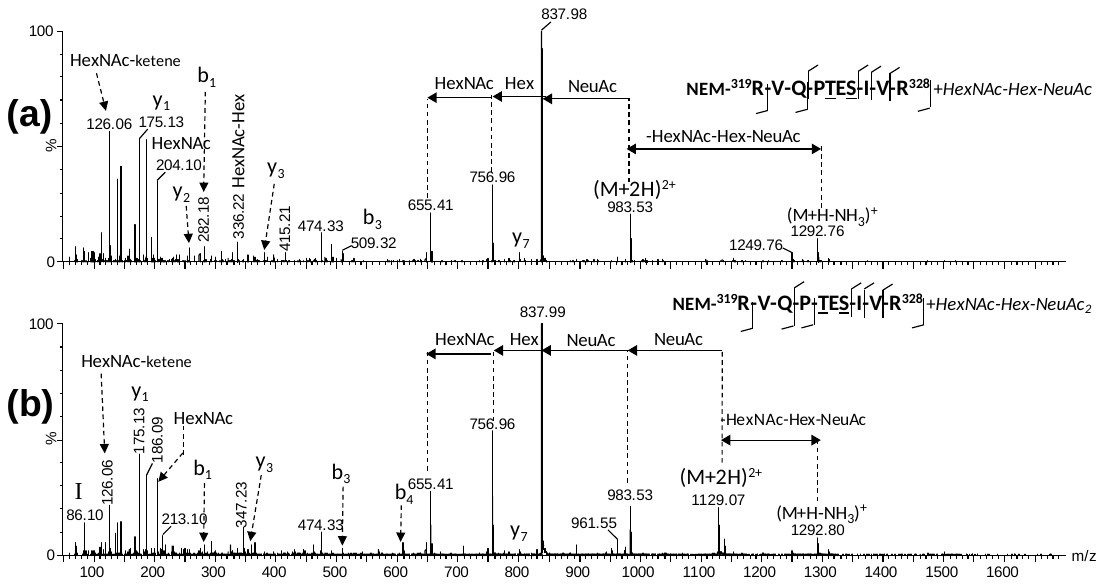

### **Fig. S12.** ESI-MS/MS spectrum of the peptides K_458_-R_466_ (*m/z*_Exp_ 622.36, 2+) and K_529_-K_535_ (*m/z*_Exp_ 457.78, 2+) containing NEM at Lys_458_ and Lys_529_ detected in RBD_(319-541)_-HEK_A3.

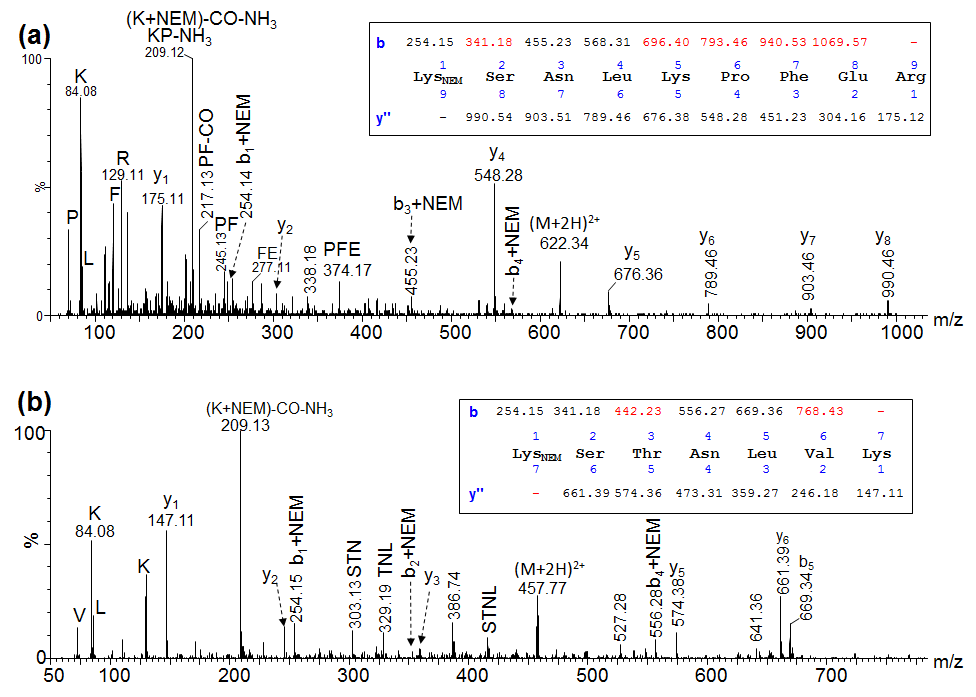

### **References**

1.	Mormann M, Eble J, Schwöppe C, Mesters RM, Berdel WE, Peter-Katalinić J, et al. Fragmentation of intra-peptide and inter-peptide disulfide bonds of proteolytic peptides by nanoESI collision-induced dissociation. Analytical and bioanalytical chemistry. 2008;392(5):831-8.

2.	Varki A, Cummings RD, Aebi M, Packer NH, Seeberger PH, Esko JD, et al. Symbol nomenclature for graphical representations of glycans. Glycobiology. 2015;25(12):1323-4.

3.	Shajahan A, Supekar NT, Gleinich AS, Azadi P. Deducing the N- and O-glycosylation profile of the spike protein of novel coronavirus SARS-CoV-2. Glycobiology. 2020;30(12):981-8.
